# Discovery and isolation of novel capsaicinoids and their TRPV1-related activity

**DOI:** 10.1101/2024.10.29.620944

**Authors:** Joshua David Smith, Vendula Tvrdoňová Stillerová, Martin Dračinský, Martin Popr, Hannah Lovinda Angermeier Gaustad, Quentin Lorenzi, Helena Smrčková, Jakob K. Reinhardt, Marjorie Anna Liénard, Lucie Bednárová, Pavel Šácha, Tomáš Pluskal

## Abstract

Chilis contain capsaicin and other structurally related molecules known as capsaicinoids. Capsaicin’s target protein, the transient receptor potential cation channel subfamily V member 1 (TRPV1), has been linked to many post-activation effects, including changes in metabolism and pain sensation. Capsaicinoids also bind to TRPV1, but current studies often disregard non-capsaicin interactions. To fill in these gaps, we screened 40 different chili varieties derived from four *Capsicum* species by means of untargeted metabolomics and a rat TRPV1 (rTRPV1) calcium influx activation assay. The resulting capsaicinoid profiles were specific to each variety but only partially corresponded with species delimitations. Based on rTRPV1 activation elicited by crude chili extracts, capsaicinoids act in an additive manner and a capsaicinoid profile can serve as a gauge of this activation. In addition, we isolated eighteen capsaicinoids, including five previously unreported ones, and confirmed their structure by NMR and MS/MS. We then tested rTRPV1 activation by 23 capsaicinoids and three related compounds. This testing revealed that even slight deviations from the structure of capsaicin reduce the ability to activate the target, with a mere single hydroxylation on the acyl tail reducing potency towards rTRPV1 by more than 100-fold. In addition, we tested how rTRPV1 activity changes in the presence of capsaicin together with non-activating capsaicin analogs and weakly activating capsaicinoids and found both classes of molecules to positively modulate the effects of capsaicin. This demonstrates that even such compounds have measurable pharmacological effects, making a case for the use and study of natural chili extracts.

**Graphical Abstract:** 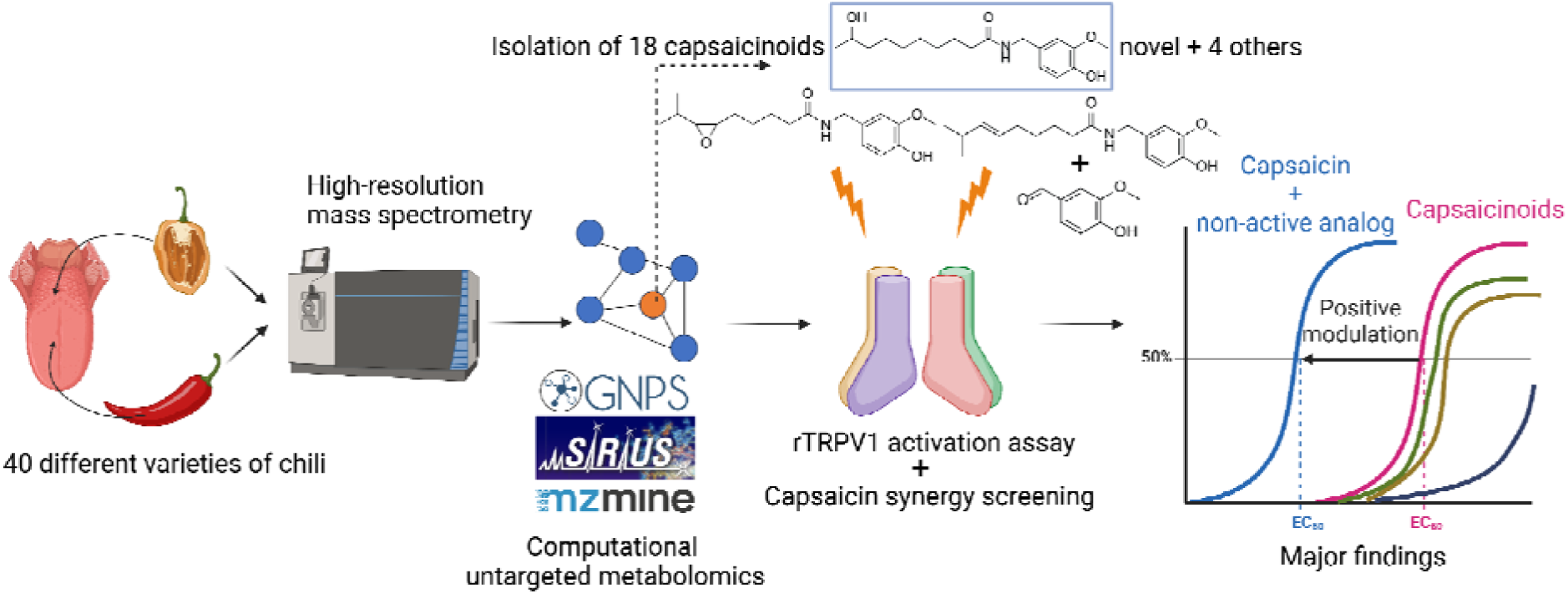

Created in BioRender. Smith, J. (2025) https://BioRender.com/a61h668

**Highlights:**

- 5 novel capsaicinoids (vanilloids) and 13 other capsaicinoids were isolated from chilis.
- The slightest deviation in the capsaicin structure results in decreased TRPV1 activity.
- A single hydroxylation can reduce potency to TRPV1 100-fold.
- Non-pungent vanilloids modulate capsaicin-TRPV1 activity.

## 1. Introduction

Chilis are a broad term for variously spicy fruits produced by different species of *Capsicum.* There are 26 known species of chili peppers with many varieties and cultivars, but only five species have been domesticated: *C. annuum* var. *annuum*, *C. chinense*, *C. frutescens*, *C. baccatum* var. *pendulum*, and *C. pubescens* (Liu et al., 2023). Some of the world’s hottest chilis, such as Habanero, Carolina Reaper and Pepper X, are varieties of the species *C. chinense*. At the time of our study, Carolina Reaper was established as the spiciest pepper in the world, but it has since been dethroned by Pepper X (Atwal, 2024). Chilis are ranked based on a measurement system called the Scoville scale, which originally utilized human taste subjected to series of sugar and ethanol dilutions of chili pepper extracts; the number of dilutions needed to neutralize the spice were reported as the number of Scoville Heat Units (SHU) (Scoville, 1912). Currently, Scoville measurements are typically conducted by determining the content of capsaicin and dihydrocapsaicin using high-performance liquid chromatography (HPLC) or gas chromatography (“How Do You Measure the ‘Heat’ of a Pepper?,” 2022; Woodbury, 1980). Modern computational metabolomics allows for more thorough investigations of biological samples compared to traditional mass spectrometry (MS) analysis. In particular, untargeted metabolomics combined with tandem mass spectrometry (MS/MS) has burgeoned in the past decade with the development of computational tools such as MZmine, SIRIUS and GNPS. (Schmid et al., 2023; Nothias et al., 2020; Dührkop et al., 2019). Despite these technological advancements, capsaicinoids have so far been investigated only in a targeted manner.

Capsaicin and dihydrocapsaicin are typically the two most abundant capsaicinoids found in chilis; however, inconsistencies in capsaicinoid profiles across and within varieties have been reported (Zewdie and Bosland, 2001). All capsaicinoids share a common structural scaffold consisting of a vanilloid head attached via an amine bond (“neck”) to an acyl tail of varying length (Fig. 1*A*). To date, about 25 different capsaicinoids have been reported (Fig. 1*A* and Supplementary Table S1) (Attuquayefio and Buckle, 1987; Constant et al., 1996; Higashiguchi et al., 2006; Ishimov et al., 2011; Jurenitsch et al., 1979; Kobata et al., 2010, 1998; Ochi et al., 2003; Wang et al., 2020). For some of these compounds, only tentative structures without NMR confirmation are available or have been published without corresponding NMR spectra (Ishimov et al., 2011; Jurenitsch et al., 1979). Therefore, not all data regarding capsaicinoids are up to current “FAIR – Findable, Accessible, Interoperable, and Reusable” standards (Wilkinson et al., 2016).

**Fig. 1.**
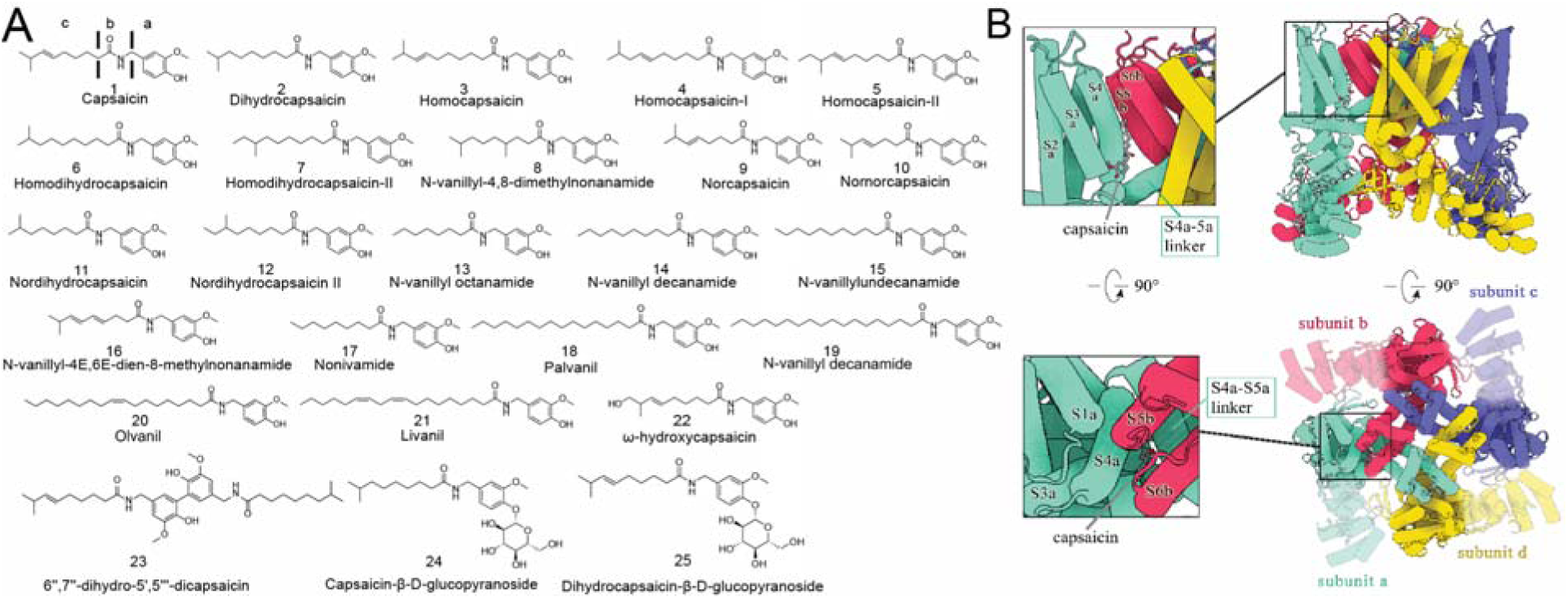
Capsaicinoids and the TRPV1 receptor. (A) Previously isolated capsaicinoids. All capsaicinoids can be divided into three parts (highlighted on compound **1** – capsaicin): a – vanilloid head; b – amide bond (neck) region; and c – alkyl tail (Elokely et al., 2016). (B) Cryo-EM structure of TRPV1 in complex with capsaicin (PDB: 7LPB)(Kwon et al. 2021). The TRPV1 structure and related images were generated using Protein Imager (Tomasello et al. 2020).

Chili extracts are offered as over-the-counter dietary supplements, as well as analgesic creams and patches containing varied mixtures of capsaicinoids. However, most studies involving capsaicinoids have focused solely on capsaicin or dihydrocapsaicin (Attuquayefio and Buckle, 1987; Kurian and Starks, 2002; Stipcovich et al., 2018), which limits our understanding of how capsaicinoids function as a group. The relevance of this understanding is exemplified in *Capsicum* extracts administered in clinical trials, for example NCT06363305 or NCT03489226 (“Capsimax Effect on Metabolic Rate, Satiety and Food Intake,” 2018; “Impact of Sex in the Effect of Dietary Capsaicin on Cardiovascular Health,” 2024). Capsaicinoids have been of relevance due to their ability to bind to the transient receptor potential cation channel subfamily V member 1 protein (TRPV1), found in nociceptive neurons (Caterina et al., 1997). TRPV1 is a homotetramer membrane channel activated by voltage, heat, capsaicin, the double-knot toxin, and the centipede toxin RhTx and each of these stimulants activates TRPV1 through a specific part of the channel (Cao et al. 2013; Bae et al. 2015; Yang et al. 2015). The capsaicin-binding pocket is surrounded by helixes S4, S6 and the S4–S5 linker (Fig. 1*B*). When capsaicin binds to TRPV1, the channel transition starts by gating and proceeds through changes in the selectivity filter (Chu et al., 2020; Dong et al., 2019; F. Yang et al., 2015). The fully activated channel results in a flux of cations (preferentially divalent) (Caterina et al. 1997) and a plethora of downstream biological effects, including a sensation of pain and heat (Caterina and Julius, 2001; Patapoutian et al., 2003).

Despite the use of chili extracts in over-the-counter products and in clinical trials, we have a relatively limited understanding of how extracts from different *Capsicum* varieties, containing diverse mixtures of capsaicinoids, can influence TRPV1 activity. Furthermore, many identified capsaicinoids have not been individually tested for their effects on TRPV1. We therefore set out to examine how different capsaicinoid profiles and individual capsaicinoids influence TRPV1 activation. In addition, suspecting that the breeding of ever-hotter varieties might have resulted in TRPV1 agonists even more potent than capsaicin or dihydrocapsaicin, we performed an untargeted metabolomic screening of 40 different chili samples together with an rTRPV1 activation assay.

## 2. Methods

### 2.1 Plant collection and sample preparation

In obtaining material for this study, we collaborated with the Botanical Garden of the Faculty of Tropical AgriSciences of the Czech University of Life Sciences in Prague. Each year, the garden regrows chilis to maintain a collection of chili seeds. Chili sowing begins between January and February and continues into summer and early fall months. In cases of increased pest infestation, plants are subjected to chemical treatment with acetamiprid, a common insecticide. The botanical garden has confirmed the varieties in its collection by comparing their morphological traits with those of plants grown at other botanical gardens or by verified dealers. For our present study, we harvested ripe chilis on several occasions between the months of August and October. Ripeness was assessed based on fruit coloration (Po et al., 2018), (Fig. 2*A*). In total, we collected the fruits of 40 different varieties covering a wide range of heat levels (Supplementary Table S2). Unfortunately, at the time of collection, we were unable to obtain fruits of the species *C. pubescens*, which has been reported to have atypical capsaicinoid fingerprints, and we did not include wild chilis (Zewdie and Bosland, 2001) in the scope of our study.

**Fig. 2.**
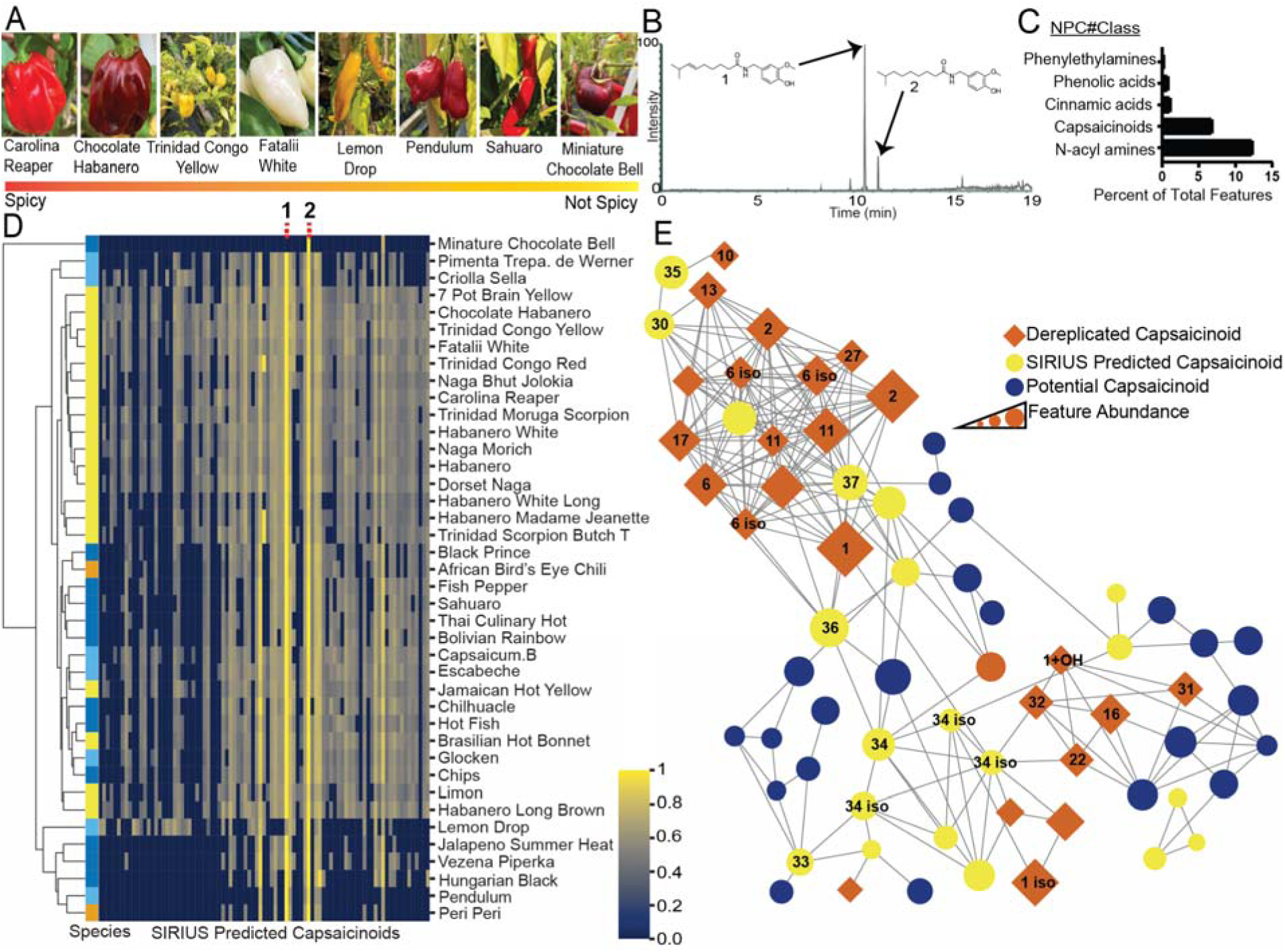
Untargeted metabolomic analysis highlighting features related to capsaicinoid composition. (A) Pictures of chilis before harvesting. (B) Representative example trace of an LC–MS base peak chromatogram of a Carolina Reaper extract. (C) Top five molecular classes predicted by SIRIUS based on the MS data of crude extracts. (D) Clustering and heatmap of all capsaicinoid-like compounds predicted by SIRIUS, capsaicin **1** and dihydrocapsaicin **2** are highlighted. Columns correspond to a specific feature with each cell representing the peak area found in a given variety (i.e., the row). Peak areas were log_10_ transformed and then normalized by rows to the most abundant feature per variety. The resulting normalized abundance of a feature is highlighted on the color scale from 0 to 1. Species are color-coded as follows: yellow (#F0E442) – *C. chinense*, blue (#0072B2) – *C. annuum*, sky blue (#56B4E9) – *C. baccatum*, orange (#E69F00) – *C. frutescens*. (E) Main capsaicinoid network based on all 40 varieties. Most labeled nodes are confirmed using reference standards with retention time (RT). Dereplicated nodes are features with predicted chemical formulas matching the chemical formulas of known capsaicinoids. Some nodes are labeled **iso**, meaning they have an MS/MS match for the same compound but have a different RT. Other molecules marked with **+OH** are indicative of hydroxylation. The strong homology of these compounds highlights the importance of having either a standard or NMR data for structure confirmation. Compounds denoted as potential capsaicinoids contained fragments indicative of a vanilloid head but were not predicted to be capsaicinoids by CANOPUS.

There, chili plants analyzed in our study were sown in winter months, from December to February, transplanted to pots throughout March and finally replanted into polycarbonate-covered flower beds or backup containers at the end of May. The chilis were grown in an ordinary horticultural substrate fertilized with cow manure. Chilis and sometimes stems were collected when ripe – based on size, shape and color – and immediately placed on dry ice. They were then stored at −80□°C until further processing, in which they were crushed to a powder with a pestle and mortar under liquid nitrogen, lyophilized and again stored at−80□°C until analysis and experimentation. While each variety had several chilis collected either from an individual plant or from several plants, they were combined as a single, pooled biological replicate. In certain cases, several samples were recollected to confirm the presence of a capsaicinoid or for isolation purposes only.

### 2.2 Chili extraction

Approximately 1 g of chili powder was used for extraction with 25 ml of ethyl acetate (EtOAc, Thermo Fisher) and agitation provided by an IKA dispenser T18 ultra-turrax (IKA, Staufen, Germany) for 1 minute. This process was completed twice for each variety. Pooled extracts from each variety were evaporated and then subjected to acetonitrile:hexane partitioning with a ratio of 1:1. The acetonitrile portion was dissolved to 1 mg/ml in acetonitrile (ACN), spun down at high speeds. A portion of the spun-down sample was diluted to 0.2 mg/ml and analyzed by a single injection (i.e., a single technical replicate) on UPLC-HRMS (details in section 2.4). Individual capsaicinoids were isolated with maceration in EtOAc, fractionation on normal-phase flash chromatography and final isolation of compound with HPLC; details can be found in Supplementary Methods 1.2.

### 2.3 Dereplication of capsaicinoids

We also applied our dereplication pipeline to aid in the search for novel capsaicinoids. By querying Reaxys and Wikidata, we were able to identify 25 previously reported capsaicinoids from *Capsicum* (Fig. 1*A*). Our output from the search was greater than 25 compounds with some inconsistencies in the literature and the storage of data. Capsaicin was sometimes stored with a SMILES string indicative of a mixture of *cis* and *trans* isomers despite natural capsaicin existing almost exclusively in its trans-form. The carbon–carbon double bond adjacent to the terminal isopropyl group is speculated to be formed by the enzyme desaturase (Blum et al., 2003). While many desaturases result in *cis* configuration, plants are known to be home of many atypical fatty acids with those in the *trans* configuration (Cerone and Smith, 2022; Millar et al., 2000). However, this is yet to be confirmed experimentally. We also had to contend with the known problem concerning the nomenclature of the different compounds referred to as homocapsaicin (Thompson, 2007). To avoid confusion, we refer to the structure/name pairs used in the original publication (Jurenitsch et al., 1979). The confusion regarding homocapsaicin is caused by the fact that homocapsaicin-II is reported as homocapsaicin in the works of Wang et. al. (Wang et al., 2020). Furthermore, many studies reporting on the structural confirmation or isolation of capsaicinoids relied solely on MS characterization without library matching or orthogonal confirmation. Although the structures themselves are still reliable for dereplication purposes, the interpretation of analytical results concerning capsaicinoids should be taken with caution, especially with suspected or known alkyl tail isomers. Lastly, two other capsaicinoids, linvanil and myrvanil, have been suggested to be found in *Capsicum* oleoresin; however, these have never been isolated from this extract, so they were left off the list (Kobata et al., 2010).

### 2.4 UPLC–HRMS analysis

An Orbitrap ID-X mass spectrometer (Thermo Fisher, Waltham, MA, USA) coupled to a Vanquish Duo UHPLC system (Thermo Fisher) was used. The temperature of the column compartment was set to 40□°C. A Waters Acquity BEH C18 150□×□2.1 mm, 1.7 μM, 130 A column (Waters corporation, Milford, MA, USA) was used. Samples were injected in 1 μl quantities. A Waters Vion mix (Waters corp.) was injected as a separate instrument QC sample. Mobile phase A: LC–MS-grade water + 0.1% MS-grade formic acid. Mobile phase B: LC–MS-grade acetonitrile + 0.1% MS-grade formic acid (Thermo Fisher). The flow rate was set to 0.350 ml/min. 0 minutes 5%B → 0.5 minutes 5%B → 15.5 minutes 100%B → 17.3 minutes 100%B →17.3 minutes 5%B → 19.3 minutes end of run. The ion source was set to 3000 V, the gas mode was set to static, 50 sheath gas (arb), 10 aux gas (arb), 1 sweep gas (arb). The temperature of the ion transfer tube was 325°C and the vaporizer temperature was 350°C. The Orbitrap resolution was set to 60,000 with a scan range of 100–1000 m/z, the rf lens was set to 45%, and the maximum injection time was set to 118 seconds. The MS/MS was set to 0.6-second cycles, with 1.0□×□10^5^ signal intensity triggering the MS/MS scan. Fragmented ions were added to an exclusion list with a 5 ppm tolerance for 2 seconds after initial triggering. Isotopes were also excluded. Apex detection was used with a FWHM value of 4 and a 50% window. The MS/MS isolation window was 0.8 m/z, with HCD fragmentation with a fixed energy mode at 35%. MS/MS detection took place in the Orbitrap, with a resolution of 15,000 and 80 ms maximum injection time. Capsaicinoid standards (MedChemExpress, Monmouth Junction, NJ, USA) were diluted to 10 μM, capsiate (MedChemExpress) and vanillin (Fluka, Charlotte, USA) to 100 μM, and 4-hydroxy-3-methoxybenzyl alcohol (Alfa Aesar, Haverhill, USA) to 1 mM, all with LC-MS grade ACN.

### 2.5 Untargeted metabolomics data processing

Thermo Fisher .raw files obtained from the UPLC–HRMS run were converted to .mzML files using MSconvert. The MSconvert settings were set to values recommended in MZmine documentation (Li-chin, 2023). The converted files were then processed using MZmine version 4.0.8 (Heuckeroth et al., 2024; Schmid et al., 2023). Processed MZmine data and the corresponding feature table was exported to .mgf files using both the SIRIUS and GNPS export. Exported files were processed on GNPS’s feature-based molecular networking platform and SIRIUS version 4.13.3. All specific parameters for mzmine processing can be found in Supplementary Methods 1.1.

All exports from SIRIUS, GNPS and MZmine were post-processed using Python scripts (GitHub) for database dereplication and statistics.

### 2.6 UV and CD

The UV absorption of isolated capsaicinoids used for testing were measured using a Thermo Fisher Nanodrop One^c^ spectrophotometer. DMSO was used as a blank. ECD and absorption spectra were measured on a Jasco 1500 spectrometer (Jasco, Tokyo, Japan) over a spectral range of 180 nm to 350 nm in dACN (1.5 mM solutions). Measurements were made in a cylindrical quartz cell with a 0.02 cm path length, using a scanning speed of 10 nm/min, a response time of 8 seconds, standard instrument sensitivity and 3 accumulations. After a baseline correction, the resulting spectra were expressed in terms of differential molar extinction (Δε) and molar extinction (ε), respectively.

### 2.7 NMR

NMR spectra were measured on a 500 or a 600 MHz Bruker Avance III HD spectrometer (Billerica, MA, USA) (^1^H at 500.0 or 600.1 MHz and ^13^C at 125.7 or 150.9 MHz) in CD_3_CN or DMSO-*d*_6_. The spectra were referenced to solvent signals (CD_3_CN: δ = 1.94 and 1.32 ppm for ^1^H and ^13^C, respectively; DMSO-*d*_6_: δ = 2.50 and 39.52 ppm for ^1^H and ^13^C, respectively). The signal assignment was performed using a combination of a 1D (^1^H and ^13^C) experiment and 2D correlation experiments (H,H-COSY, H,C-HSQC and H,C-HMBC). In some cases, the sample quantity was insufficient for the detection of some or all ^13^C signals in the 1D experiments. The structural analysis and signal assignment was then based on the 2D correlation experiments. All NMR data were deposited in the NP-MRD (see Data Availability); all ID’s for individual compounds can be found in Supplementary File S2.

### 2.8 QC of Isolated compounds

Samples designated for the TRPV1 activity assay were submitted to the quality control (QC) procedure to confirm the integrity of each sample using an automated high-throughput LC-MS protocol on the ACQUITY UPLC I-Class (Waters). The analytes were applied on a reversed phase column (ACQUITY UPLC BEH C18 1.7μm, 2.1×50mm, Waters) using a fast (1.5 min) gradient elution with 10-95% acetonitrile. The exact mass of the molecule was looked for in the signal from the mass spectrometry detector to confirm the identity, while purity was determined by integrating the area under peaks in the signal from the diode array detector (DAD) chromatogram.

### 2.9 TRPV1 activity assay

Purchased capsaicinoid standards, capsiate (both MedChemExpress), vanillin (Honeywell Fluka) and 4-hydroxy-3-methoxybenzyl alcohol (Alfa Aesar) and AMG-517 (TargetMol, Boston, MA, USA) were all dissolved in DMSO to 10 mM as stock solutions. Stably transfected irTRPV1-FlpIn293 cells with Tet-on system (procedure described in Supplementary Methods 1.3) were grown to confluence in Advanced Dulbecco’s Modified Eagle Medium (Adv-DMEM/F12, #12634) supplemented with 10% FBS, 1% Pen Strep, 1% Glutamax (all Gibco, Thermo Fisher Scientific). Forty-eight hours before the experiment, doxycycline (final concentration 1 μg/mL)(Sigma-Aldrich, St.Luid, Mi., USA) and 10 μM ruthenium red (RR; Merck KGaG, Darmstadt, Germany) were added to the media to induce TRPV1 expression. After 24 hours, the cells were washed and detached with 0.5% trypsin–EDTA in PBS (Gibco). Cells resuspended in blocking media (Adv-DMEM/F12 + 10% FBS + 10 μM RR) were counted and then seeded into a 384-well plate (3674, Corning, Corning, NY, USA) in the amount of 3×10^4^ per well. Twenty-four hours after seeding, a 2× Fluo-4 direct dye solution (Invitrogen, Thermo Fisher Scientific), dissolved in HHBS buffer, pH ∼ 7.4, was added to well with cells and incubated for 30 minutes at 37°C and then for 40 minutes at room temperature. Once incubated with the dye, the supernatant was removed using BlueWasher (BlueCatbio NH Inc., Lebanon, NH, USA) and changed to an HHBS + 10 μM sulfinpyrazone (Merck KGaG) solution added by Certus (Fritz Gyger AG, Gwatt, Switzerland), followed by 10 minutes of incubation in FDSS μCell (∼ 30°C; Hamamatsu, Hamamatsu city, Japan). A separate compound plate was prepared using an Echo dispenser 650 (Beckman Coulter Inc.,Brea, CA, USA). The final concentration was reached by adding HHBS + 10 μM sulfinpyrazone as above by Certus. The rTRPV1 activity measurement started by recording the baseline (40 s, sample interval 0.339 s). Then the compounds/compound mixture were transferred to the dedicated wells by tips while the recording was running for the next 5 minutes. The 1% TritonX-100 (Merck KGaG) was used for normalization of the data by an internal laboratory management system ScreenX (version 2.0, developed at Institute of Molecular Genetics, Prague, Czechia) and GraphPad Prism (version 10.2.2 (397), GraphPad Software). All dose/concentration-response curves were plotted as a four parametric logistic function made using least-square fit with separately handled replicates with the following equation:

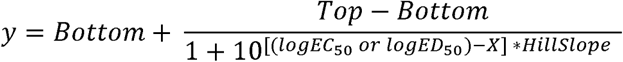

The number of replicates and other statistical relevant data are linked with relevant tables (Supplementary Table S3-S5).

## 3. Results and Discussion

### 3.1 Untargeted metabolomics of chili varieties reveals novel candidate capsaicinoids

To explore the chemical space of capsaicinoids contained in chili peppers, we performed an untargeted metabolomic investigation of 40 chili varieties covering a wide range of heat levels (Fig. 2*A* and Supplementary Table S2), which are in long-term cultivation at the Botanical Garden of the Faculty of Tropical AgriSciences of the Czech University of Life Sciences in Prague.

#### 3.1.1 SIRIUS and CANOPUS investigation of data based on predicted compound classes

Despite extensive research in the area, studies on capsaicinoids have mostly been limited to targeted analysis of already known compounds (Kozukue et al., 2005; Peña-Alvarez et al., 2009; Reilly et al., 2001; Sganzerla et al., 2014; Wu et al., 2019). To start exploring the chemical space and distribution of capsaicinoids amongst different chili varieties, and to explain differences in TRPV1 activation observed between varieties, we first screened 40 chili varieties, listed in Supplementary Table S2, in an untargeted manner by means of UPLC–MS/MS (Fig. 2*B*). To obtain a holistic understanding of the chili metabolome, we then leveraged MZmine for raw data processing and “feature-finding” (i.e., feature–a “meaningful” MS signal produced by a small molecule occurring in the sample) (Heuckeroth et al., 2024; Schmid et al., 2023), GNPS for feature-based molecular networking (Nothias et al., 2020), and SIRIUS for feature annotation. To help identify capsaicinoids, we used CANOPUS, a part of SIRIUS, which allows for compound class prediction and SIRIUS molecular formula prediction (Dührkop et al., 2020; Ludwig et al., 2020). Results from CANOPUS indicated “N-acyl amines” as the largest compound class that accounted for 12.3% percent of all features found across all varieties (Fig. 2*C*). N-acyl amines are the necessary precursors to the acyl moieties of all capsaicinoids (Fig. 1*A*). By contrast, features predicted to be phenylethylamines, simple phenolic acids and derivatives of cinnamic acid, such as the capsaicinoid precursors vanillylamine and vanillin, accounted for about 1–2% of all features. Furthermore, we did not detect vanillylamine in any chili sample. It has been reported that low levels of phenylethylamines are due to their quick conversion into capsaicinoids (Ogawa et al., 2015). The excess of N-acyl amines could further indicate that these phenylethylamines are a limiting factor in capsaicinoid biosynthesis and diversity. As expected, capsaicinoids were one of the largest predicted classes of molecules at 6.85% (Fig. 2*C*). Other major compound classes present are not considered in this paper.

The initial screening of 40 chili varieties revealed only 1530 features, which is a relatively small number for that amount of samples, indicating that many features are shared by the varieties. However, there was no discernible intraspecies clustering based on capsaicinoids profiles and the hierarchical clustering of metabolomic features (Fig. 2*D*). One example that highlights this irregular pattern is the variety Lemon Drop, which contains more dihydrocapsaicin than capsaicin. This observation is consistent with previous targeted capsaicinoid experiments, which indicated that the composition of capsaicinoids is not a good chemotaxonomic identifier (Zewdie and Bosland, 2001) and was specific to each variety. It is important to note, a single pooled biological replicate (i.e., several chilies of the same variety were combined) was used for crude chili extracts.

#### 3.1.2 Exploration for novel capsaicinoids and capsinoids

Contrary to expectation, we found no capsiate or other capsinoid in our initial screening. Capsiate is a non-pungent analog of capsaicin found in *C. annuum* (Fig. *S1*, compound **39**) (Kobata et al., 1998). The main difference between the two molecular classes lies in their precursors, which are vanillyl alcohol and vanillylamine, respectively (Sano et al., 2022). To form capsinoids, the vanillyl alcohol condenses into an ester bond which has decreased stability in polar solvents when compared to our extraction solvent (ethyl acetate) (Sutoh et al. 2001). It is speculated that a putative aminotransferase (pAMT) in capsaicinoid biosynthesis requires a point mutation to produce vanillyl alcohol (Lang et al., 2009) and downstream capsinoids. More recently, it has been suggested that a cinnamyl alcohol dehydrogenase (CAD) produces vanillyl alcohol and not the pAMT mutation (Sano et al. 2022). We identified small amounts of vanillyl alcohol in some varieties (1000-fold lower signal intensity than a 1 mM standard), which would support the hypothesis of a CAD vanillyl alcohol formation as we have both vanillyl alcohol and capsaicinoids in the same extract. Lastly, our ionization method tailored to capsaicinoids is unfavorable for capsinoids due to in-source fragmentation. For example, capsiate’s base peak ion corresponds to a vanilloid head and contains only a small [M+H]+ peak (Table1). If capsinoids are present at a low concentration, the in-source fragmentation can result in [M+H]+ ions at intensities too low for feature identification or MS/MS for molecular networking. Future studies should consider milder ionization strengths to minimize in-source fragmentation.

To identify possible novel capsaicinoids, we constructed a feature-based molecular network on the GNPS platform, which groups clusters of molecules based on similarities in their MS/MS (Fig. 2*E* *and S2*). Capsaicinoids are identifiable by CANOPUS molecular class annotation and verifiable based on the presence of capsaicin and other nodes with the vanilloid head MS/MS fragment of m/z = 137 (Wu et al., 2019). Using this network, we selected several targets for isolation based on mass or fragmentation indicative of hydroxylation, tail length, or dimerization, assuming that the target compounds would produce a spectrum of TRPV1 activation.

### 3.2 Isolation and purification of capsaicinoids resulted in five novel structures

In order to investigate the effects of both crude extracts and individual capsaicinoids on the TRPV1 receptor, we proceeded to purify the target compounds from our initial screening. A pooled selection of lyophilized chilis were macerated in ethyl acetate and pre-fractionated by flash-chromatography prior to the isolation of individual compounds using semi-preparative HPLC (Fig. 3*A* and 3*B*). In total, we isolated 18 capsaicinoids, including five novel molecules and several others that were previously only obtained through synthesis (Fig. 3*C* and Table 1). Information about the MS/MS fragmentation of each isolated compound can be found in Supplementary File S1, and NMR data are provided in Supplementary File S2.

**Fig. 3.**
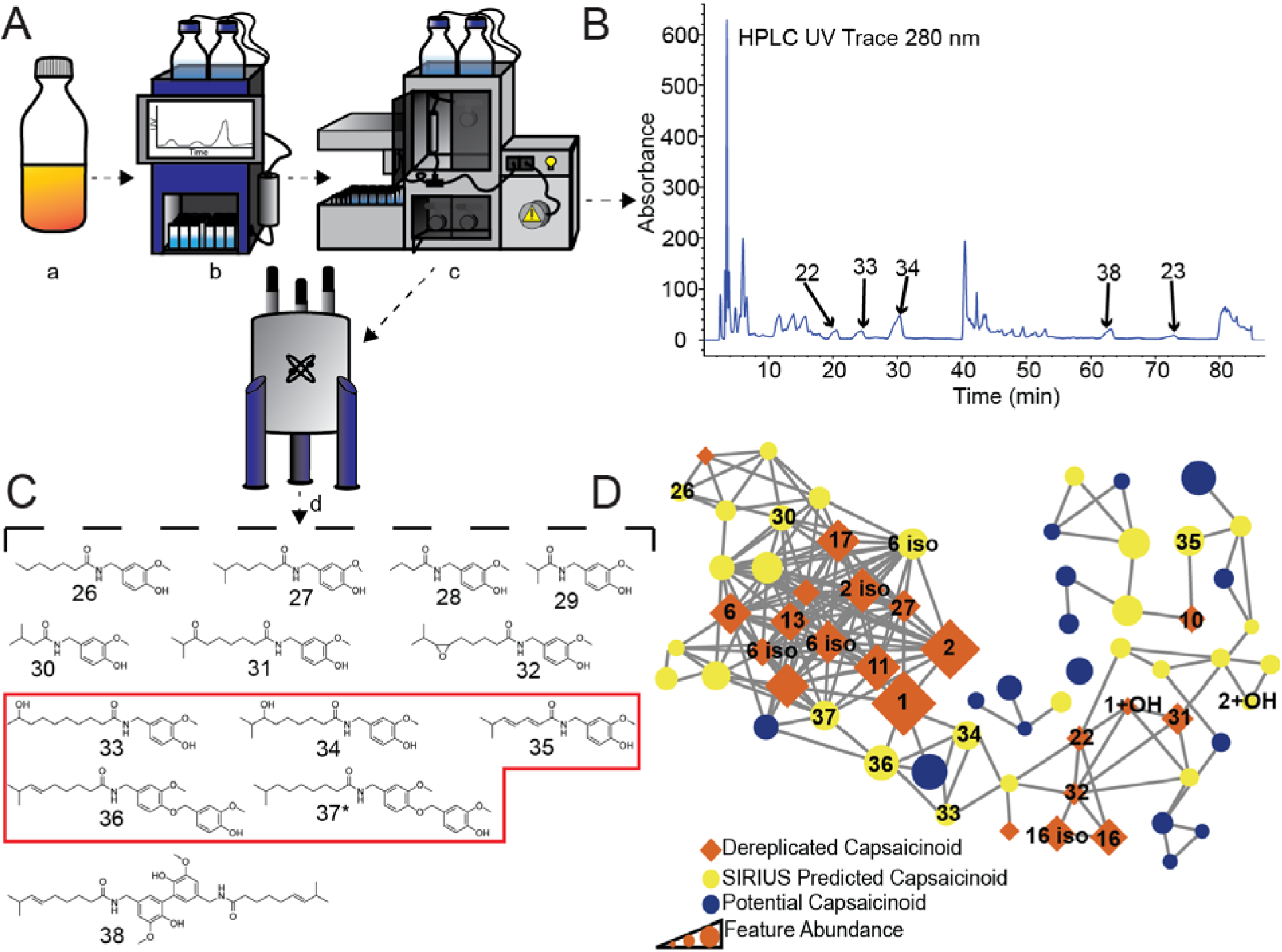
Isolation of capsaicinoids. (A) Overall schema of the workflow: maceration (a) of chilis was followed by normal-phase flash-chromatography (b), isolation of individual compounds on HPLC (c) and lastly structure confirmation with NMR (d). (B) HPLC chromatogram from one of the pooled flash fractions highlighting the isolation of individual compounds. (C) Structures of newly isolated compounds. The red box highlights the novel capsaicinoids. *Compound **37** was identified based on the NMR structure of **36** followed by MS/MS analysis. (D) Molecular network used for compound identification before isolation. Most labeled nodes are nodes confirmed based on a reference standard or isolation. Some labeled nodes, such as **6 iso** or **1 + OH**, are nodes that had a library match to a compound with the same precursor match but a different RT. The node corresponding to capsaicinoid **34** was identified as **33** with a .002 better cosine score; however, the RT did not match. Capsaicinoids **28** and **29** were not in the initial network because of their low intensity, and dimers (**23** and **38**) were also in a separate network. Denoted as potential capsaicinoids are compounds that contain an MS/MS fragment indicative of a vanilloid head but predicted to be a different molecular class from capsaicinoids.

**Table 1.**
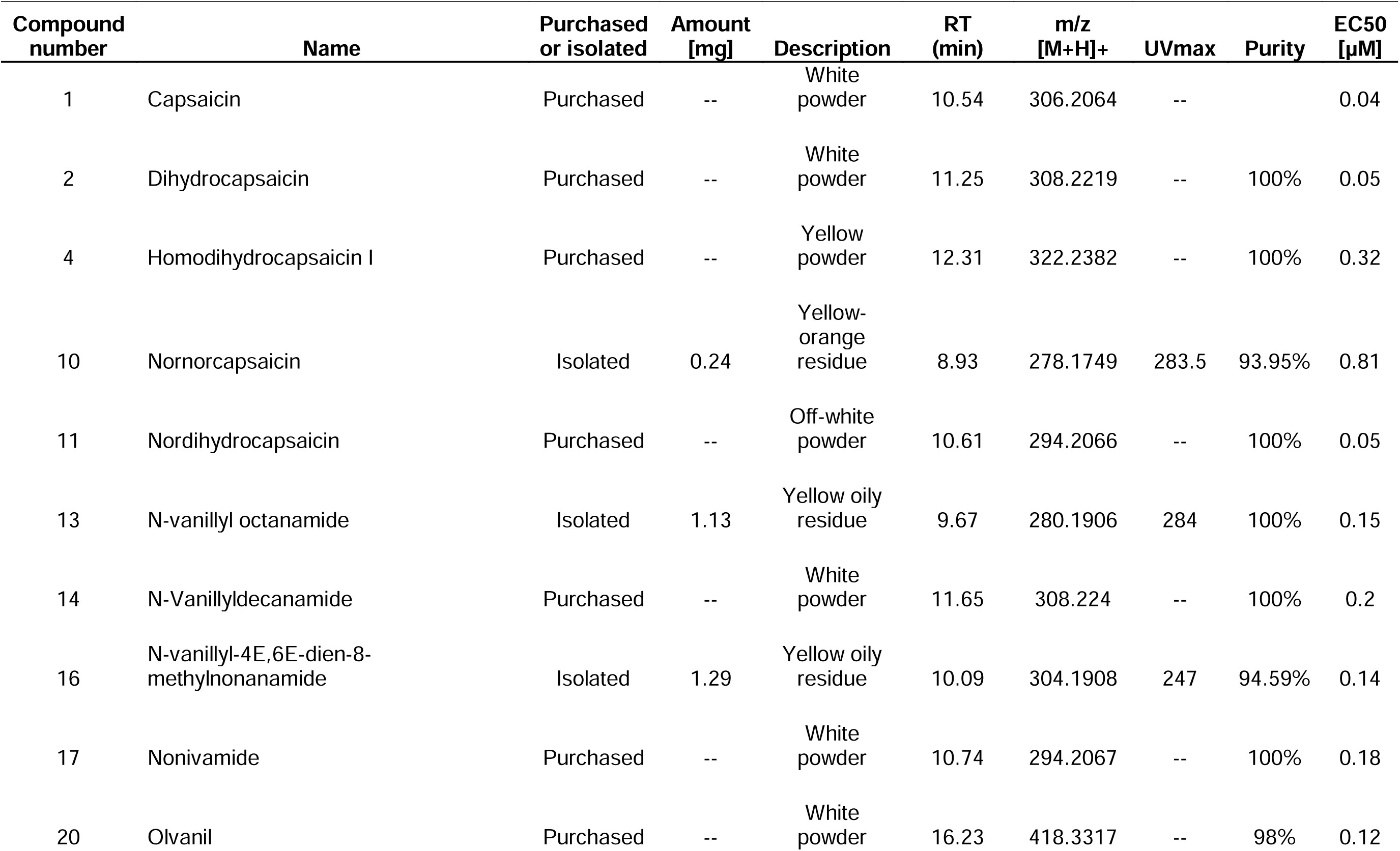
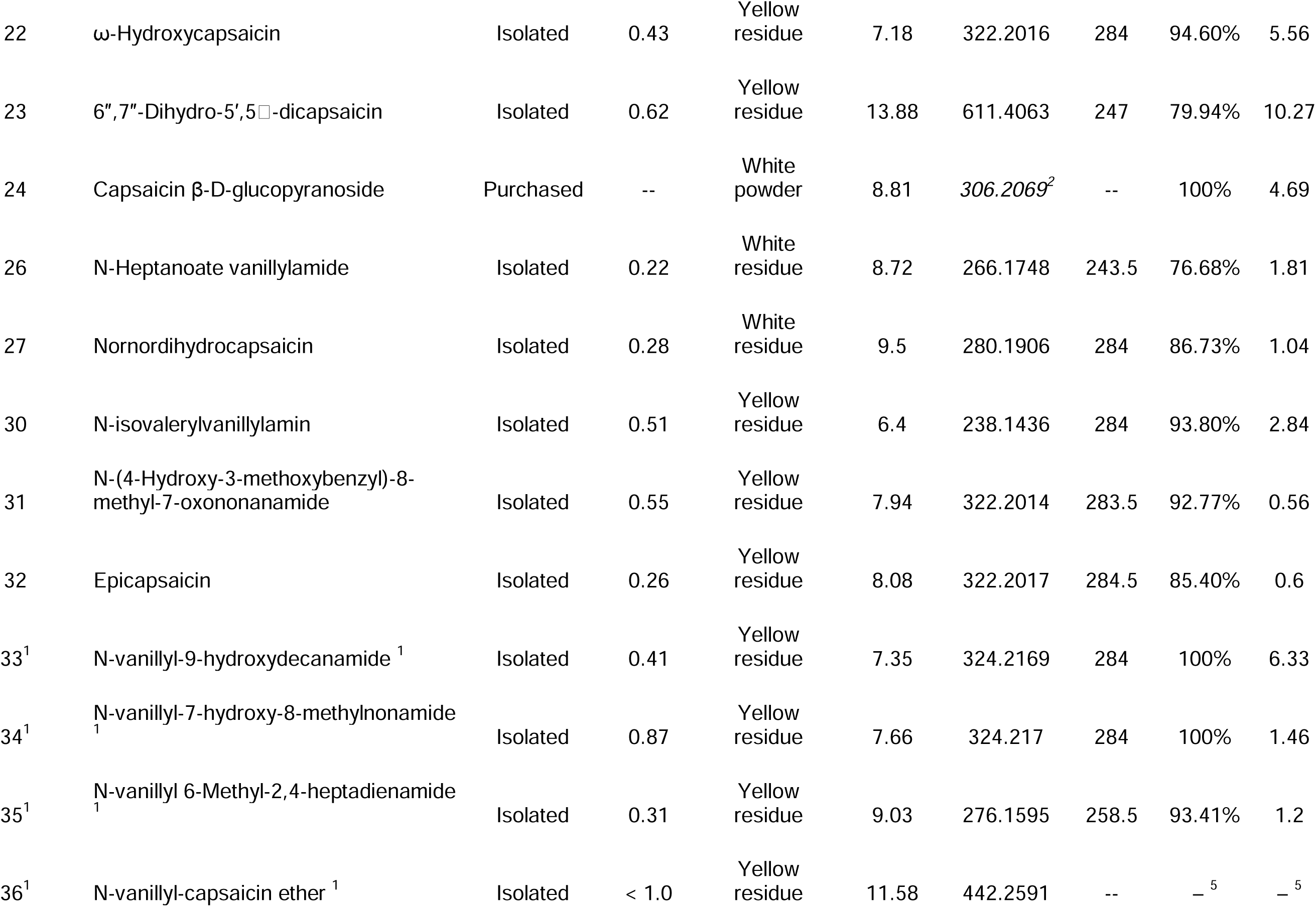
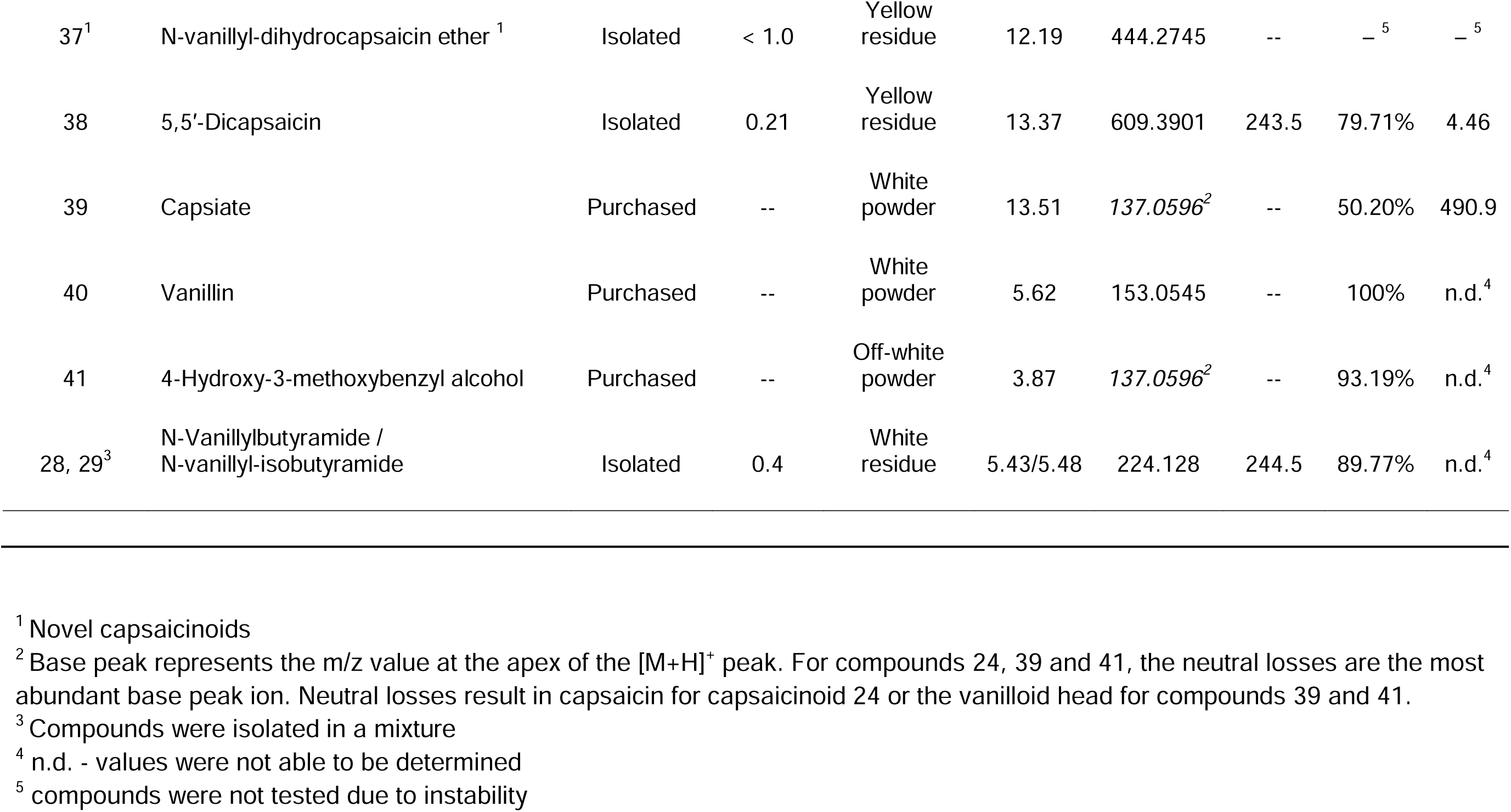
Summary of properties of used compounds; RT – UPLC-HRMS retention time, m/z [M+H]^+^ – base peak mass^2^, UV_max_ – maximum UV absorbance, EC_50_ - median effective concentration on TRPV1.

We successfully annotated, with a high degree of confidence, 26 out of 50 nodes (52%) predicted to be capsaicinoids in our final network of pooled chilis by comparing the retention times and characteristic ions of isolated compounds with those of purchased reference standards (capsaicinoids **1**, **2**, **4**, **11**, **14**, **17**, **20** and **24**) (Fig. 3*D*). Nodes predicted to be capsaicinoids that are left unannotated correspond to substances of varying tail length or differing in hydroxylation. We removed all redundant adducts and other artifacts from the network (Fig.□*S3*).

Among those, capsaicinoids **10**, **13** and **26–30** with varying tail length were expected based on SIRIUS formula prediction and manual inspection of MS/MS spectra. Capsaicinoids **10** and **13** were previously reported (Gannett et al., 1988; Wang et al., 2020). Interestingly, capsaicinoids **26**, **29** and **30** have only been synthesized non-enzymatically and have never been isolated from a chili species (Ježo, 1975; Nelson, 1919; F. Yang et al., 2015) whereas capsaicinoid **28** has been synthesized only enzymatically (Kobata et al., 1998). Capsaicinoid **27** has been synthesized and only tentatively isolated from *C. chinense* and *C.annuum* without any NMR or reference standard confirmation (Buitimea-Cantúa et al., 2020; Stipcovich et al., 2018; Sweat et al., 2016; Takahashi et al., 1976).

Capsaicinoids **22**, **31**, **32**, **33**, **34** are tail-oxidized. ω-Hydroxycapsaicin, capsaicinoid **22**, was first identified as a microbial metabolite (Reilly et al., 2003), later isolated from *C. annuum* (Ochi et al., 2003) and then as a product of capsaicin p450 oxidation(Deng et al., 2022). Likewise, capsaicinoid **31** has previously been identified only as a product of human cell line oxidation, and capsaicinoid **32** has hitherto only been known as a product of microbial p450 enzyme catalysis and epiphytic processing (Halme et al., 2016; Ma et al., 2021; Migglautsch et al., 2018; Xin et al., 2017). Lastly, to our knowledge, capsaicinoids **33** and **34** are reported here for the first time.

Only two capsaicinoids, **16** and **35**, were doubly reduced. Capsaicinoid **16** was isolated previously (Wang et al., 2020) whereas capsaicinoid **35** is a new structure first reported here.

Capsaicinoids **36** and **37** are both also newly identified and present a unique structure indicative of S_N_2 chemistry on the vanilloid hydroxyl group. It is unlikely that capsaicinoids **36** and **37** are a photocatalyzed product, because these reactions typically result in insertion-like products, as seen in capsaicinoid **23**. The structure of capsaicinoid **36** was confirmed by NMR, which enabled us to determine capsaicinoid **37** by MS/MS analysis. Although both compounds can be isolated and were found in several harvests, they are markedly unstable based on repeated MS and NMR measurements. There were other nodes also indicative of possible acyl modifications to the vanillyl head of the capsaicin molecule, but these features were less pronounced and harder to generate a structure for with only MS/MS.

Two capsaicinoid dimers, **23** and **38,** were isolated. We obtained reliable NMR structural confirmation for 5,5’-dicapsaicin, capsaicinoid **38**. Capsaicinoid **23** was previously isolated from *C. annuum* (Ochi et al., 2003) and was confirmed based on MS/MS differences compared to capsaicinoid **38**. Interestingly, capsaicinoid **38** has not been previously identified as a direct product of chili extraction, however it has been identified as a product of different reactions (Tateba and Mihara, 1991;Bernal and Barceló, 1996;Henderson et al., 1999;Reilly et al., 2013; Tian et al., 2019;Martínez-Juárez et al., 2004).

It was not possible to determine the absolute configuration of capsaicinoids with chiral centers, **22, 33** and **34,** using standard circular dichroism (CD) experiments. This inability to determine their absolute configuration was likely caused by the flexibility between the chromophore and the chiral center itself (Saito and Schreiner, 2020). Because of the limited amount of material available, CD spectroscopy structural assignment was not tested.

### 3.3 Capsaicinoids and related molecules modulate capsaicin-induced TRPV1 activation

With the individual capsaicinoids at hand, we investigated the effects of both the isolated compounds and crude extracts on TRPV1 activation. Among laboratory model organisms, the human TRPV1 receptor (hTRPV1) shares the greatest degree of homology with the rat TRPV1 receptor (rTRPV1): 83% sequence similarity (Neuberger et al., 2023). In addition, rTRPV1 and hTRPv1 have been reported to have similar capsaicin-elicited activity, only rTRPV1 is less sensitive to capsazepine and ruthenium red, two common antagonists of TRPV1 (McIntyre et al., 2001). We used the Flp-In T-REx-293 cell line along with the rTRPV1 receptor and tetracycline repressor to create an inducible line denoted as irTRPV1-FlpIn293. Surface receptor expression was confirmed by immunocytochemistry staining and functionality (Fig. S4). Our irTRPV1–FlpIn293 system under doxycycline control can express an excessive amount of receptors, which makes the sensitivity of our model close to that of radioactivity assays (Caterina et al., 1997; Lee et al., 2003).

#### 3.3.1 Activation of rTRPV1 by crude chili extracts correlates with their capsaicinoid content

We began screening all crude chili extracts against our rTRPV1 assay (example traces of receptor responses, Fig. 4*A*). Most extracts activated the receptor with a response 60% or greater. The crude Miniature Chocolate Bell (S25) extract elicited a slight partial activation at the highest concentration tested, which may likely be caused by its lower capsaicinoid content (Supplementary Table S2), although the presence of an inhibitor or negative modulator in the mixture cannot be ruled out. See Supplementary Table S2 and S3 for all values pertaining to crude extracts.

**Fig. 4.**
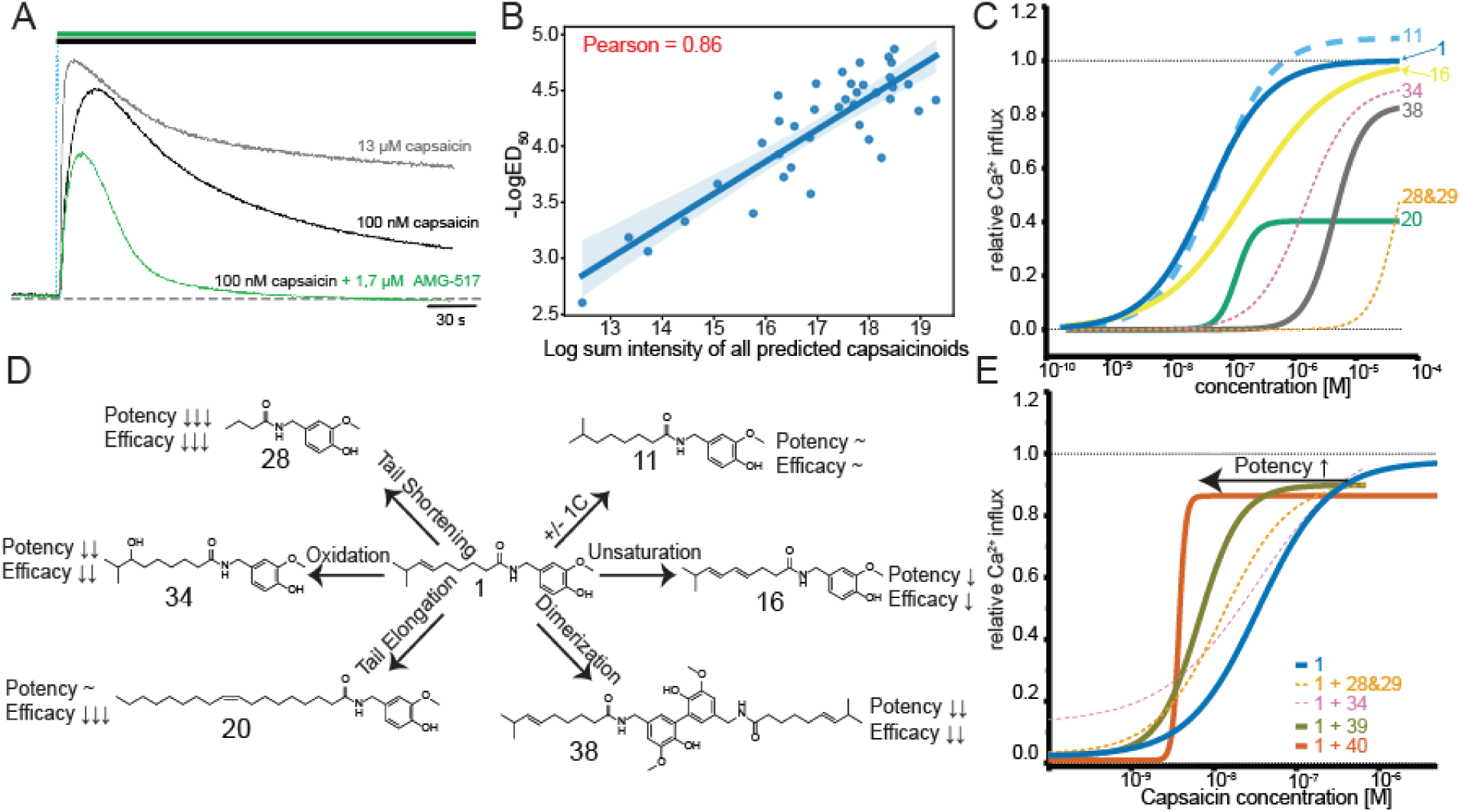
Effects of chili extracts and pure capsaicinoids on rTRPV1 activation. (A) Example recorded traces from the rTRPV1 activity assay used for investigation of activity of capsaicinoids. All the compounds/compound mixtures were applied in parallel and not washed (top bars: black - first compound, green - second compound). (B) Relation between total capsaicinoid content and TRPV1 activation. Each point corresponds to one chili variety. For each variety, the sum intensity of MS peak area for predicted capsaicinoids is plotted against its corresponding −LogED_50_ value. S7 and S25 were removed from the data analysis, see Supplementary Table S2 and S3. Removal of the two varieties did not change the overall positive trend. (C) Activation curves of capsaicinoids representative of their group. All values were normalized against that of capsaicin. Complete data of the curves are listed in Supplementary Table S4 (D) Modifications of capsaicinoids and how they affect rTRPV1 response. Relative to capsaicin, most modifications result in decreased potency and efficacy. (E) Capsaicin-induced rTRPV1 activation modulated by a non-potent analog or precursor. Concentrations of capsaicinoids added: **28** and **29** – 1.7 μM, **34** – 0.2 μM, **39** – 1.7 μM, **40** – 1 μM. These single concentrations were added to a dilution series representing a full dose– response curve of capsaicin. A full description of the curves for combinations of capsaicinoids is provided in Supplementary Table S5.

As stated in the previous section, we see that capsaicinoids are not a good taxonomic indicator; however, we were curious if there would be a relation between capsaicinoid composition and rTRPV1 activation. Indeed, our data revealed a strong correlation (Pearson = 0.86) between the capsaicinoid composition (i.e MS sum intensity of all capsaicinoid predicted features, Supplementary Table S2) of a given chili variety and the median effective dose (ED_50_) at values of the corresponding crude extracts (Fig. 4*B*, Supplementary Table S3). We expanded this comparison with different MS feature groupings and ED_50_ values (Fig. *S5A-D*). When we exclude the capsaicin and dihydrocapsaicin features, the correlation remains strong (Fig. *S5B*). This led us to believe that rTRPV1 activation may have an additive effect, i.e., more capsaicinoids results in greater activation. When focused on remaining features, Pearson correlations greater than 0.7 still exist, alluding to the possibility of a non-vanilloid feature influencing the rTRPV1 receptor activity. In future studies, the collection size could be increased to obtain a more jointly normally distributed representation (Schober et al. 2018) by increasing both the number of varieties and replicates. Overall, we considered this correlation as a suggestion that all capsaicinoids present should be accounted for and not just a select few. To confirm this, we tested both capsaicinoids individually and in pairs with capsaicin on rTRPV1 activation.

#### 3.3.2 Overview of activity of the tested capsaicinoids and related compounds

It is known that the vanilloid head of capsaicin is the most sensitive to atomic structure modification (Walpole et al., 1996). In addition, modification to the neck can critically alter the recognition and potency of the compound (Vu et al., 2020; F. Yang et al., 2015), as seen also with piperine (Dong et al., 2019; McNamara et al., 2005), gingerols (Yin et al. 2019; Dedov et al. 2002) and synthetic capsaicin analogs (Vu et al., 2020). It is also established that TRPV1 activation decreases as the tail of the capsaicinoid becomes shorter (F. Yang et al., 2015). Although these trends are known and many other synthetic agonists have been tested on TRPV1 (Lee et al., 2004, 2003; Park et al., 2004; Ryu et al., 2008), there is still a large knowledge gap regarding the potency of naturally occurring capsaicinoids. We have filled this gap by testing 24 capsaicinoids and three other related compounds in an rTRPV1 activation assay (Table 1).

Of the 18 isolated compounds, only compounds **13**, **26** and **28** were, to the best of our knowledge, previously tested for TRPV1 activation (F. Yang et al., 2015). Of the purchased standards, other than capsaicin, capsaicinoids **2**, **17** and **20** were tested on dorsal root ganglia that express TRPV1, with respectively slightly higher median effective concentration (EC_50_) values of 0.19 μM, 0.55 μM and 0.17 μM (Walpole et al., 1993). Differences are explainable due to different testing systems. Capsaicinoids **14** and **17** were also previously tested, yielding EC_50_ values of 0.061 μM and 0.072 μM, respectively, in a ^45^Ca^2+^ uptake assay utilizing rTRPV1-expressing Chinese hamster ovary cells (Cho et al., 2012). Our EC_50_ value for capsaicinoid **14** indicated almost three-fold lower potency, but within a range expected considering the differences in assay methods and expression systems. As far as we have been able to ascertain, no other capsaicinoid standards have been tested with a TRPV1 expression system.

Most compounds tested were 90% pure or more. Impurities typically consisted of unknown non-capsaicinoid compounds, possible isomers of the target compound, or capsaicinoids usually at 100-1000 times less intense than the target compounds based on inspection of the raw UPLC-MS/MS data. Compounds 23 and 38, the dimers, contained trace amounts of monomers, which may inflate the reported EC_50_ values. The purity of capsiate was the largest concern, which showed a 50% purity despite manufacturer reporting >94% (Alomone Labs). We attribute this lower purity to the storage of capsiate in DMSO, which has been shown to reduce capsinoid stability (Sutoh et al. 2001). QC testing occurred several months after initial capsiate testing with TRPV1; however, we recommend finding an alternative solvent or to prepare capsinoid solutions fresh before each assay. In general, all compounds were of high purity and should produce comparable EC_50_ values in future studies. As this was a discovery based study, we would recommend synthesizing compounds reported in this manuscript for future testing to ease subsequent purification.

#### 3.3.3 Effect of tail modifications on rTRPV1 activation

Whereas the head is important for TRPV1 receptor gating, in both a positive (Chu et al., 2020; Dong et al., 2019; F. Yang et al., 2015) and negative (McDonnell et al., 2002; Thomas et al., 2011) manner, the exact length of the tail has been shown to have an effect on capsaicin’s strong potency (F. Yang et al., 2015). The tail of capsaicin is arranged by series of van der Waal interactions with the S6 helix of the capsaicin-binding pocket (see Fig. 1*B*) of the TRPV1 receptor (Vu et al., 2020; F. Yang et al., 2015; Yang and Zheng, 2017) followed by the movement of the S4–S5 linker and the S6 helix caused by the binding of the vanilloid head (Yang et al., 2018). Capsaicinoids **2**, **16** and **17** had the same number of carbons as capsaicin but differed either in their connectivity or in the number of hydrogens. Unsaturation, saturation or linearization of the tail while maintaining the same number of carbons as capsaicin only slightly reduced the potency of the molecule (Fig. 4*C* and 4*D*). Because the capsaicin tail interacts in the TRPV1 hydrophobic pocket through a series of van der Waals interactions, small losses of activity are consistent with small changes in carbon chain length and similar tail polarity. Capsaicinoids **22** and **31–34** also contained the same number of carbon atoms but were oxidized somewhere on the tail, and depending on the position of the oxygenation, EC_50_ values were reduced between 10- and 150-fold. Based on these results, any oxidative process, even on the tail, can be linked to lower TRPV1 activity. We attribute the reduction of potency due to tail-hydroxylation residing in a predominantly hydrophobic core and its interaction with the S6 helix. The pocket likely struggles to accommodate the polar oxygen and restricts the movement of the S6 helix upon vanilloid binding. Only three capsaicinoids, **4**, **11** and **17**, which differed by a single carbon atom, had similar EC_50_ values and were 1–6 times less potent than capsaicin. Nordihydrocapsaicin, **11**, had the most comparable TRPV1 activation concentration to capsaicin and dihydrocapsaicin of 0.05 μM (Fig. 4*C* and 4*D*).

Other capsaicinoids with 2–5 fewer carbons on the tail and those that had varying connectivity were markedly less potent than capsaicin, exhibiting 15-fold lower to almost zero activity. The sole exception, capsaicinoid **13** (N-vanillyl octanamide) was only about three times less potent than capsaicin. Olvanil (**20**) and N-vanillylbutyramide/N-vanillyl-isobutyramide (**28** and **29**) both differed by six carbons. These three compounds were all partial agonists, but olvanil had a similar EC_50_ value to capsaicin (Fig. 4*C* and 4*D*). Short-tailed capsaicinoids have been investigated previously (F. Yang et al., 2015) and are known to be ineffective at opening TRPV1. Conversely, olvanil might be able to adopt a different position in which the binding of the vanilloid is less permissive due to the bulky tail interacting with the pulling of the S6 helix (Vu et al., 2020). The complete list of obtained EC_50_ values is in Table 1. The detailed concentration–response curve data are presented in Supplementary Table S4. We further confirmed that the measured responses were due to rTRPV1 activation by using the competitive TRPV1 antagonist AMG-517 (Doherty et al., 2007) (Fig. *S6*).

#### 3.3.4 Modulation of rTRPV1 activity by not-potent capsaicinoids

Capsaicin, other than in the form of a pure active ingredient, is always found in a mixture of capsaicinoids. The complexity of these mixtures varies from variety to variety and from extraction to extraction. Based on our observations with the crude extracts (Fig. 4*B*), we asked if any of the capsaicinoids (Table 1) could also act in an additive or synergistic manner and wanted to better understand the role of non-potent analogs. We performed a series of experiments using EC_10_, EC_50_ and EC_90_ concentrations of capsaicin together with several concentrations of isolated capsaicinoids or purchased standards (data not shown). By comparing the curves of individual capsaicinoids to curves that represent a combination of capsaicinoid with capsaicin, we selected combinations at concentrations that indicated possible modulation. Following, we measured full concentration-response curves for capsaicin applied together with denoted compounds (detailed data are shown in Supplementary Table S5). We considered strength (i.e., synergistic or additive) of interaction based on the shape and left-shift of the curve (Zhao et al. 2010; Tallarida 2011). While some weakly activating capsaicinoids, as illustrated by the curve of capsaicinoid **34** in Fig. 4*C&E*, might simply be additive to strong TRPV1 agonists, the results indicate that less potent capsaicinoids, in general, tend to act as positive modulators and increase the overall potency of the capsaicin / capsaicinoid mixture (Fig. 4*E* and Supplementary Table S5). Non-activating compounds, namely capsaicinoids **28** and **29**, capsiate **39** and the precursor vanillin **40**, resulted in the greatest positive modulation of capsaicin induced TRPV1 activation.

A similar leftward shift of the capsaicin concentration-response curve was observed when TRPV1 expressing cells were pretreated with adenosine-5’-triphosphate (ATP) before agonist application (Tominaga et al. 2001). ATP modulation of TRPV1 was originally linked to G-protein coupled purinergic receptors (Tominaga et al. 2001). Our irTRPV1–FlpIn293 is based on a HEK293 cell line, therefore we also expect the expression of purinergic receptors and a number of other endogenously expressed receptors (Atwood et al. 2011). The number of endogenously expressed receptors compared to induced rTRPV1 in our system is very low. Since we applied capsaicinoid mixtures contemporaneously and examined only response peaks (Fig. 4*A*), we do not attribute positive modulation as downstream effects from receptors other than rTRPV1. However, this cannot be excluded without further studies.

Capsaicin-induced TRPV1 channel opening begins with the binding of the vanilloid head (Yang et al. 2018) and a single capsaicin pocket occupancy is insufficient to evoke full receptor activation (Liu et al. 2019). This is in line with the calculated probability of a TRPV1 channel opening for capsaicin, which is 4.58% for one bound capsaicin molecule and 36% per two bound molecules (Li and Zheng 2023). The critical part that limits the opening is the S6 helix (Jara-Oseguera et al. 2019; Yang et al. 2018) close to the receptor gate, not the selectivity filter at the top of the receptor. Therefore, a single molecule of capsaicin might be sufficient to evoke full channel response, but when in combination with another vanilloid, regardless of tail length, if it can at least influence the transition of the receptor’s gate. In addition to the capsaicin pocket, the TRPV family of receptors are known to share 16 different ligand-binding sites, some with potential allosteric activities (Yelshanskaya and Sobolevsky 2022). Although the transition of one subunit doesn’t decrease the energy necessary for transition of another subunit (Liu et al., 2019; Yang et al., 2018; Zhang et al., 2021), the cooperativity from a different activating site is still possible and these small non-activating molecules, such as vanillin **40** or capsaicinoids **28** and **29**, might bind to a known, for example in ankyrin domain (Lishko et al. 2007) or unknown allosteric site. Taken together, non-pungent vanilloids (i.e., capsaicinoids **28** and **29** and related vanilloids **39** and **40**) appear to collectively facilitate the transition of the TRPV1 channel or increase its stability, most probably utilizing the vanilloid head. The exact mechanism (effect on the gate by a vanilloid head, allosteric modulation or other options) needs to be addressed by further studies.

## 4. Conclusions

In the process of exploring capsaicinoids, we have assembled the most comprehensive library of naturally occurring capsaicinoids with NMR structure confirmation. Our data show that most variations of capsaicinoid molecules reside in varying carbon tail length, sometimes with different degrees of oxidation or reduction, in line with findings of previous studies. Some of the capsaicinoids we discovered bear modifications to their vanilloid heads. However, these compounds seem to be unstable, based on repeated NMR and MS measurements, so we excluded them from further testing.

Previously, it was speculated, based on targeted analysis, that capsaicinoids were of no use for chemotaxonomic identification (Zewdie and Bosland, 2001). Our untargeted analysis corroborates this suspicion because the capsaicinoid fingerprints obtained did not allow for reliable species identification. Variation in the composition of capsaicinoid mixtures is important when considering chilis for use in medicinal products. Furthermore, CANOPUS predicted compound classes resulted in few identified vanilloid precursors indicating that these molecules may be a limiting factor in the biosynthesis and diversification of capsaicinoids.

Capsaicinoid mixtures from natural chili extracts, although not species-specific, were found highly effective to gauge rTRPV1 activation. Mixtures acted synergistically and were not dependent on capsaicin or dihydrocapsaicin as the sole driving force of rTRPV1 activation. The potency of most capsaicinoids is proportional to their similarity to capsaicin. Oxidation of the tail greatly reduces the potency of capsaicinoids, so protection against oxidation is recommended for maintaining their potency. Capsaicinoids with long acyl tails have similar potency but are less efficient compared to capsaicin. Conversely, shorter-tailed capsaicinoids are progressively less active and those with a tail about two carbons long are inactive altogether. Nevertheless, despite the loss in individual potency, these capsaicinoids act together synergistically.

Additionally, non-activating compounds can function as positive modulators of capsaicin activity. The non-activating precursor vanillin, which is found in smaller concentrations in chilis, can add to the overall synergistic effect of rTRPV1 activation. Capsaicin and dihydrocapsaicin are typically the most abundant capsaicinoids, but they should not be the only compounds considered with respect to TRPV1 activation. Ultimately, all chili derived products that use normalized extracts can experience batch-to-batch variation in capsaicinoid profiles.

### 4.1 Limitations of this study

We studied only four chili species and the varieties we selected tended to be on the spicier side. For a full understanding of the chemical space, less spicy varieties and wild cultivars should be included in future studies. All capsaicin modulation and antagonism experiments began with premixing all compounds and delivering them in parallel. Experiments could also be completed with pretreatment of compounds; however, this is unlikely to change any of the conclusions drawn from our results. Although our data document synergistic activity between capsaicin, related non-activating compounds and other capsaicinoids, we have not investigated the exact mechanism of this synergism. Lastly, all the experiments were done on the rat, not human, TRPV1 receptor, and their results may not translate between the two proteins completely despite their high homology.

## Supporting information

SI_File_1_MSMS

SI_File_2_NMR

## CRediT authorship contribution statement

**Joshua Smith**: Writing - original draft, Conceptualization, Data curation, Methodology, Formal Analysis, Funding Acquisition, Investigation, Writing -review and editing **Vendula Tvrdo**ň**ová Stillerová**: Conceptualization, Methodology, Investigation, Formal analysis, Data curation, Writing - original draft, Writing -review and editing. **Tomáš Pluskal**: Conceptualization, Funding Acquisition, Supervision, Writing -review and editing. **Pavel Šácha**: supervision. **Martin Dra**č**inský**: Investigation, Formal Analysis, Writing -review and editing. **Martin Popr**: Investigation and Formal analysis, Writing -review and editing. **Hannah Lovinda Angermeier Gaustad**: Investigation. **Quentin Lorenzi**: Investigation, **Helena Smr**č**ková**: Investigation. **Jakob K. Reinhardt**: Investigation, Methodology, Supervision. **Marjorie Anne Liénard**: Resources, Writing - original draft. **Lucie Bednárová**: Investigation

## Acknowledgments

J.D.S. was supported by the Charles University Grant Agency (GA UK) and the First Faculty of Medicine of Charles University, project ID: 252182 376322 Smith. T.P. was supported by the Czech Science Foundation (GA CR) grant 21-11563M and by the European Union’s Horizon 2020 research and innovation programme under Marie Skłodowska-Curie grant agreement No. 891397. V.T.S. and M.P. were supported by the Ministry of Education, Youth and Sports of the Czech Republic (LM2023052). We want to greatly thank The Botanical Garden of the Faculty of Tropical AgriSciences of the Czech University of Life Sciences Prague (CZU) for the donation of the chilis and the continuous collaboration. Special thanks are due to the Pluskal Group, Teo Hebra for the continuous support, and Roman Bushuiev for help with the Python scripts. Lastly, we want to thank Frederick Rooks for the comprehensive edits and suggestions on the manuscript.

## Data Availability

The Python scripts can be found at https://github.com/jdsmith145/Capsaicinoids. All raw and processed data files and MSconvert batch files, detailing the MSconvert settings used, are deposited in the MassIVE repository under reference number MSV000096223. NMR data has been deposited to NP-MRD (np-mrd.org); individual identifiers can be found in Supplementary Table S1. All data from both MassIVE and NP-MRD can also be found at 10.5281/zenodo.15088148. Data from TRPV1 experiments can be found at 10.5281/zenodo.14204350.

