## Supplementary material for "Discovery and isolation of novel capsaicinoids and their TRPV1-related activity": SI_Figures

### Supplementary data

#### 1.Supplementary Methods

##### 1.1 Metabolomics parameters

Mass detection was performed for centroided MS and MS/MS signals separately, with “Noise Levels” set to $1.0 \times 10^4$ and $1.0 \times 10^3$ signal intensity, respectively. Chromatogram building was carried out utilizing the ADAP Chromatogram Builder Module, using the following parameters: a minimum of 6 consecutive scans, a minimum group intensity of $5.0 \times 10^4$, a minimum absolute height of $1.0 \times 10^5$ and m/z tolerance for scan-to-scan matching of 0.001 m/z or 3.5 ppm. Chromatograms were smoothed using the Savitzky–Golay filter with retention time smoothing and 5 points. A local minimum resolver was used for feature deconvolution, with the following parameters: retention time dimension, a chromatographic threshold of 0.95%, a minimum search range of 0.05 (retention time or mobility), 5.0% minimum relative height, a minimum peak height of $1.0 \times 10^5$, a minimum ratio of peak top/edge of 3.00, a peak duration of 0–2 and a minimum of 6 scans. “MS/MS scan pairing” was also checked in the deconvolution step using the following parameters: MS1 to MS2 precursor tolerance of 0.0020 or 5 ppm, feature edges as a retention time filter, 25.0 as a minimum relative feature height, 1 minimum required signal and 1% minimum signal intensity. Isotopic patterns were grouped using the $^{13}C$ isotope filter. The grouping was conducted with a 0.001 m/z tolerance or 2 ppm and a retention time tolerance of 0.03 minutes. A monotonic shape constraint was applied, with the most intense isotope selected as the representative one. Other isotopic peaks were identified: 0.0010 m/z or 2 ppm tolerance, maximum charge of isotope m/z to 1 and search for single most intense scans. Join aligner was utilized to align detected features across samples: alignment was performed with an m/z tolerance of 0.010 or 3.50 ppm and a retention time tolerance of 0.07 min; m/z and RT weights were set to 1. Duplicate peak filter was set to new average mode, 0.0010 m/z and 3.0 ppm tolerance and a 0.01 retention time tolerance. Feature list blank subtraction required 1 detection in blank, based on peak area with maximum ratio type and features only above a 300%-fold change were kept. Correlated features were determined with 0.01 RT tolerance, 0 and 0 were used for minimum feature height and intensity. A minimum of 1 sample was required with 60% overlap and all gap-filled features were excluded. Six points, with 3 points on the edge, were required for feature shape correlation, which was based on Pearson and required 85%. Adducts were identified with ion identity networking using 0.0010 m/z or 3 ppm mass tolerance and a 0 height. Smaller networks without major ion, delete smaller networks: link threshold (set to 4) and delete networks without monomer were all enabled. The ions used were: [M]+, [M+H]+, [M+NH4]+, [M+Na]+, [M+K]+, [M+2H]2+ and [M-H20]+]. An additional ion was manually added to the MZmine list, [M+ethylamine]+ (46.0651). Feature lists were filtered by rows requiring 1 isotope feature, MS/MS scan and feature IDs were reset. Features were then matched with the capsaicinoid library using merged MS/MS spectra, precursor tolerance of 0.0010 or 5 ppm, spectral tolerance of 0.0015 m/z or 10 ppm, removed precursor, minimum of 5 matched signals, weighted cosine similarity based on MassBank and a minimum score of 0.8; all unmatched signals were kept and matched to zero.

Exported .mgf files were then uploaded to GNPS library (40 varieties : ID=109485b44f8b437caf0db5ffe480e7af and Large-scale extraction: ID=ffca33c3005c4251b8bc9e064f383fa8). The GNPS precursor and fragment mass tolerance was set to 0.0075 Da. Minimum cosine pairing was set to 0.65, with 5 minimum matched fragments, 500 Da maximum shift in precursor. Network TopK was set to 15 and the component size was 150. All other parameters were set to their default values. For SIRIUS, version 4.13.3, the mass analyzer parameter was changed to “orbitrap”, the “bio databases” option was selected, and values greater than 800 Da were excluded from the analysis; all other SIRIUS parameters were kept at their default values.

#### 1.2 Isolation of individual capsaicinoids

*First large-scale extraction*. Different varieties of lyophilized chilies (see Supplementary Table 2 for the specific varieties) were mixed together to yield a total of about 50 g. The mixed chili powder was submerged in an excess of EtOAc and stirred for two hours at room temperature. The EtOAc was decanted, filtered through Whitman No. 1 filter papers and collected into a round-bottom flask. The EtOAc was then removed under vacuum via rotary evaporator(Buchi, Uster, Switzerland) and reused for subsequent extractions. The remaining plant material was extracted two more times in a similar manner, but with 4 hours and then 16 hours of stirring. The extracts were pooled together. This crude extract mixture was then fractionated by normal-phase flash chromatography on a CombiFlash Rf 200 system (Teledyne, Thousand Oaks, CA., USA) at a flow rate of 30 ml/min. Mobile phase A: Cyclohexane. Mobile phase B: EtOAc. 0 → 45 minutes of isocratic 45% B; 45 → 75 minute gradient from 45% B to 100% B; 75 → 85 minutes 100% B. The fractions were pooled into 7 fractions based on LC–MS analysis. These pooled fractions were isolated using a 1260 Infinity II Preparative LC system (Agilent) with a 19 mm × 250 mm Waters XBridge C18 column using a Milli-Q water:ACN(Fisher) gradient. Gradients were 55 minutes and started at either 25% or 30% ACN and went to 100% ACN.

*Second large-scale extraction*. The second large-scale extraction followed similar procedures as the first, only with some optimization to the LC methods. EtOAc in excess was added to about 89 g of pooled lyophilized chili powder. Each sample was macerated with stirring for 1 hour and 30 minutes. The extract was then filtered into a round-bottom flask, and the EtOAc was evaporated under vacuum (rotary evaporator, Buchi). The EtOAc was then reused to extract the chili sample again. This was completed six times. On the last maceration, the sample was stirred overnight at room temperature. After this overnight maceration, the extract was filtered and pooled with the previous extracts. The remaining plant material was washed with a small portion of EtOAc and added to the pooled extract. The pooled extract was then filtered through Whitman No. 1 filter papers. Normal-phase flash chromatography (Teledyne) was then repeated with an optimized gradient for better peak separation. Mobile phase A: Cyclohexane. Mobile phase B: EtOAc. 0 → 8.4 minutes isocratic 10% B, 8.4 → 16.8 minutes 10% → 20% B,16.8 → 34.9 minutes isocratic 20%B, 34.9 → 36.9 minutes 20% → 40%B, 36.9 → 56.9 minutes isocratic 40%B, 56.9 → 70 minutes 40% → 100%B, 70 → 78 minutes 70 → 78 minutes. The fractions were checked by LC–MS and pooled to maintain the highest purity of the target compound. Pooled fractions were further separated by preparative HPLC (Agilent) equipped with a 10 mm × 250 mm Waters XBridge C18 column. The LC methods were optimized using an XBridge C18 3.5 μm column (Waters) and then transferred to an XBridge C18 10 mm × 250 mm (Waters) and then fine-tuned. Gradients were optimized for each pooled flash fraction and used two isocratic hold steps for the isolation of as many target capsaicinoids as possible. Depending on the fraction, a different percentage of ACN was used for the hold step.

#### 1.3 Inducible TRPV1 Cell Line

The full-length open reading frame for rat TRPV1 was amplified by PCR in a Veriti thermal cycler (Applied Biosystems, Thermo Fisher Scientific, MA, USA) from pcDNA3-rTRPV1-N604S (generously provided from Rachel Gaudet), using gene-specific oligonucleotide primers encompassing AsiSI and MluI restriction sites and Advantage Polymerase mix (Takara Bio, USA). The AsiSI-MluI-digested rTRPV1 DNA fragment was ligated into the pcDNA™5/FRT/TO plasmid (Invitrogen, Thermo Fisher Scientific), modified to contain AsiSI/MluI cloning sites, and transformed into Stbl3 *E. coli* cells, prior to plasmid DNA purification and Sanger sequencing verification. The final pFRT-TO-rTRPV1-FLAG construct was preceded by a cytomegalovirus (CMV) promoter containing a tetracycline-controlled transcriptional repressor (2 × 19-bp $TetO_2$) sequence. Downstream of the ORF, the final expression cassette was engineered to contain a Ser-Gly (AGTGGA) peptide linker followed by a FLAG-epitope sequence (ATTACAAGGATGACGACGATAAGGTT) and a stop codon.

Flp-In T-REx-293 parental cells containing a single integrated Flp Recombination Target (FRT) site (Invitrogen) were cultured in a HERAcell 150i $CO_2$ incubator at 37 °C (Thermo Fisher Scientific) in a 10 cm tissue culture dish (VWR, Radnor, PA, USA) in Advanced Dulbecco's Modified Eagle Medium (Adv-DMEM/F12) (Life Technologies, Thermo Fisher Scientific) supplemented with 10% fetal bovine serum (FBS, Gibco, Thermo Fisher Scientific), 16 μg/mL Blasticidin and 100 μg/mL Zeocin (both Life Technologies). Flp-In™ T-REx-293™ were seeded in a 6-well tissue culture dish (VWR) at a density of $0.3 \times 10^6$ cells in 1.5 mL Adv-DMEM/F12+FBS without antibiotics. After 48 hours, cells in three separate wells were co-transfected (day 0) with 4.5 μg endotoxin-free pOG44, 1 μg endotoxin-free rTRPV1-FLAG plasmid DNAs and 10 μL Lipofectamine (L2000, Invitrogen) diluted in 300 μL Opti-MEM I Reduced Serum (Life Technologies). The next day (day 1) the transfection medium was gently decanted and replaced with 2 mL Adv-DMEM/F12+FBS for 48 hrs.

On day 3 post transfection, cells from each transfection were trypsinized using 0.5 mL Trypsin-EDTA 1X 0.25% (Corning, Corning, NY, USA) and replated in a 6-cm culture dish in 5mL Adv-DMEM/F12+FBS and 200 µg/mL Hygromycin B (Life Technologies) to start selecting for transformants. The selection medium was decanted on day 5, day 8 and day 12 post transfection and replaced with fresh Adv-DMEM/F12+FBS and 200 µg/mL Hygromycin, for a total of 3 changes. Single-cell transformants observed on the experimental transfections plates were allowed to expand for an additional 5 days, after which lines containing individual Flp-In T-REx transformant colonies were established in presence of Hygromycin and Blasticidin. Final monoclonal lines (irTRPV1-FlpIn293) were validated via qPCR and western blot after induction for 48h with doxycycline 1 mg/mL.

#### 2. Supplementary Figures

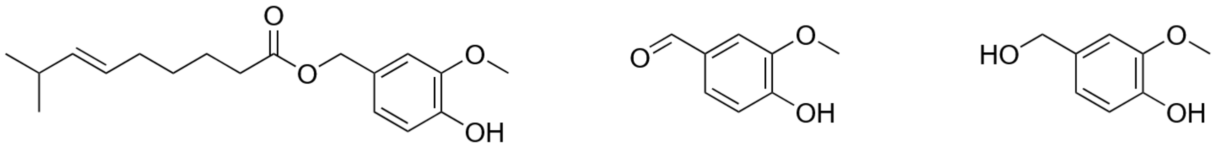


39 40 41

Fig. S1. Three compounds used in rTRPV1 testing but are not capsaicinoids. **39** is capsiate, **40** is vanillin and **41** is vanillyl alcohol.

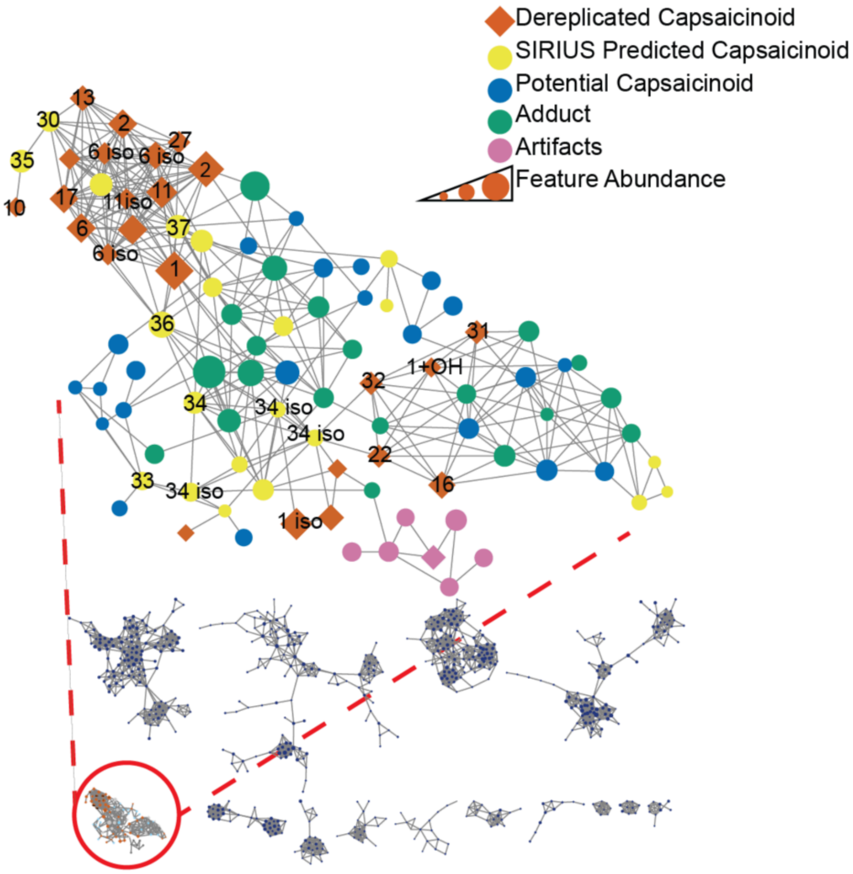


Fig. S2. Unprocessed network from the initial 40 chili extractions. Potential capsaicinoids are features indicative of a structure similar to a vanilloid head based on MS/MS spectra but were not identified as capsaicinoids by SIRIUS. Adducts are ions of repeated nodes with different counter ions, e.g. [M+H]+, [M+Na],[M+ethylamine]+ and [2M+H]+. Artifacts are features unrelated to capsaicinoids but were clustered there as a result of the networking process.

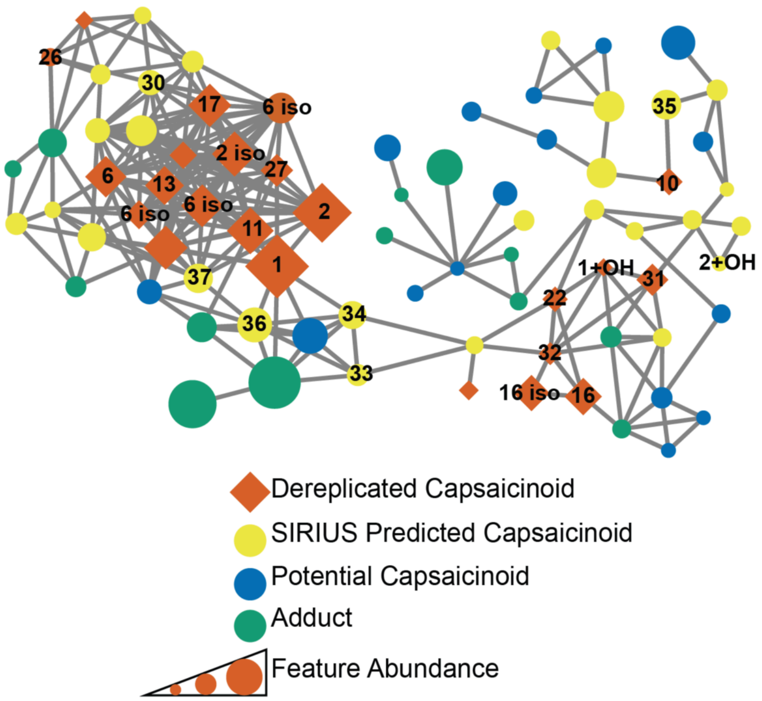


Fig. S3. Unprocessed network from the large-scale chili extraction. Potential capsaicinoids are features indicative of a structure similar to a vanilloid head based on MS/MS spectra but were not identified as capsaicinoids by SIRIUS. Adducts are ions of repeated nodes with different counter ions, e.g. [M+H]+, [M+Na],[M+ethylamine]+ and [2M+H]+.

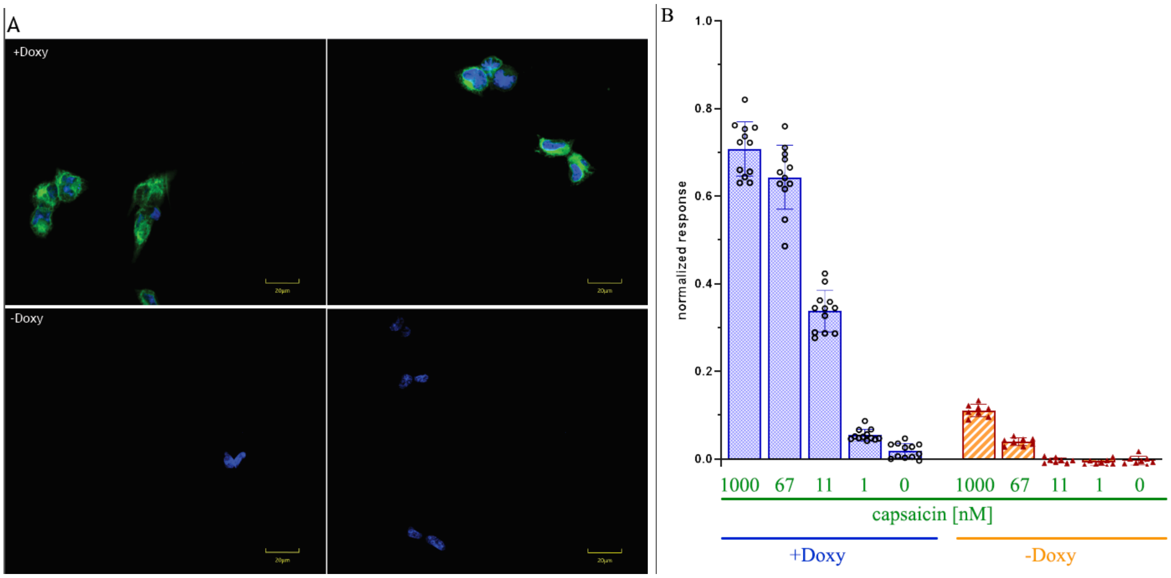


Fig. S4. Comparison of irTRPV1-FlpIn293 cell-line with and without induction by doxycycline (Doxy). (A) Immunocytochemical staining of irTRPV1-FlpIn293+Doxy (top row) and -Doxy (bottom row). Nucleus (blue; DAPI), rTRPV1 receptor (green; Santa Cruz sc-398417 (1:200, 1 hour, RT) + Abcam ab150115 (1:800, 1 hour, RT)). Recorded by Yokogawa CV8000, 60× objective. The non-induced cells alone express an enhanced number of receptors, which is not obvious from the images, because the gain was adjusted to the induced (+Doxy) cells with an excessive number of receptors. (B) Calcium influx after the application of capsaicin to induced (+Doxy, blue) and non-induced cells (-Doxy, orange), according to the protocol described in the Methods. The peak values are normalized to 1% TritonX-100.

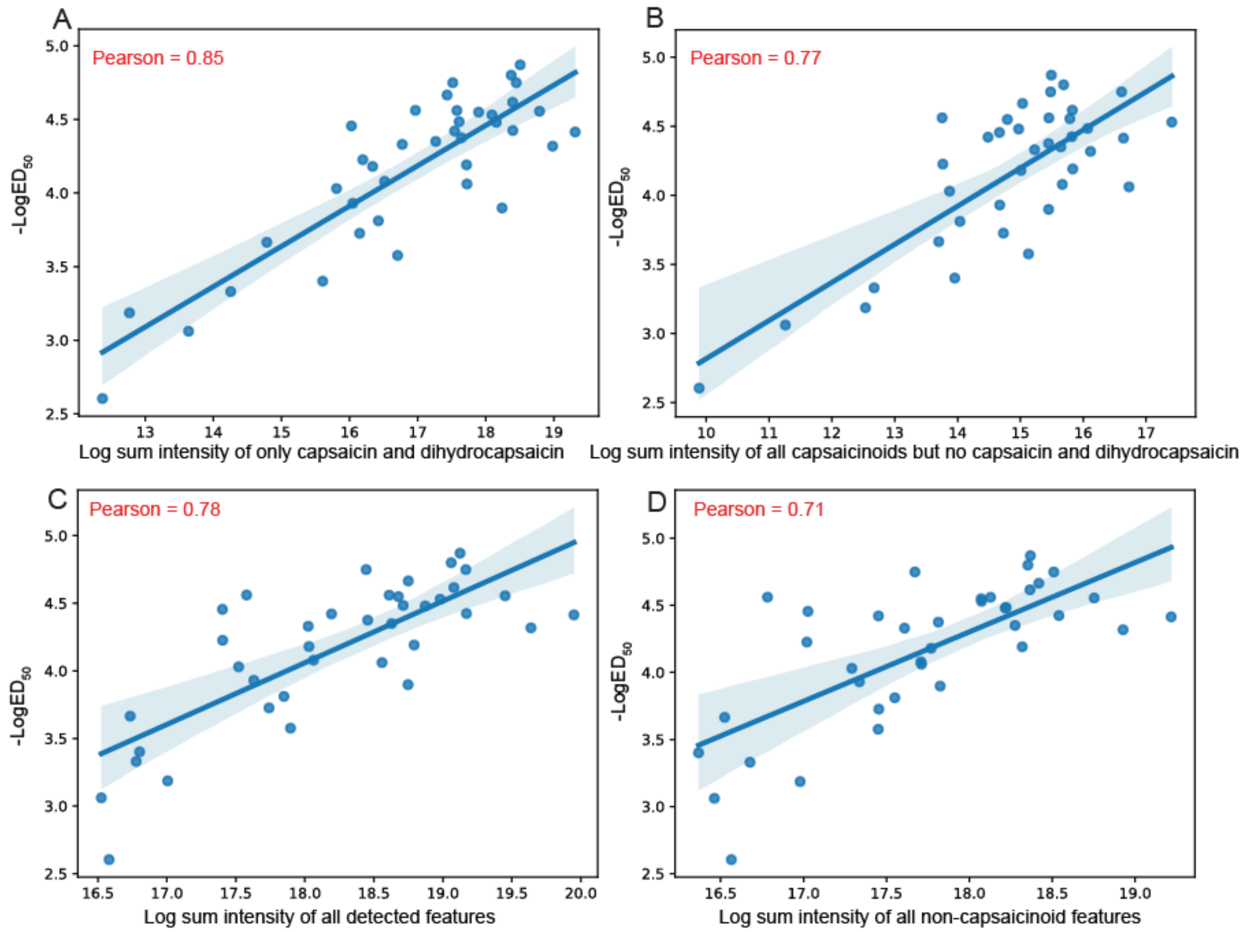


Fig. S5. Multiple comparisons of variety extract features to $ED_{50}$ values. Black Prince (S7) and Miniature Chocolate Bell (S25) were not plotted in any of the panels. (A) Sum feature area for capsaicin and dihydrocapsaicin, all other compounds were removed, Pearson = 0.85. Capsaicin and dihydrocapsaicin sums account for [M+H]+, [M+Na], [M+ethylamine]+ and [2M+H]+ adducts. (B) All capsaicinoid predicted features, but sum area for capsaicin and dihydrocapsaicin were removed. A strong correlation persists even without capsaicin and dihydrocapsaicin. (C) Pearson correlation of sum intensities of all features in a given variety to its $ED_{50}$ value, Pearson = 0.78. (D) Same correlation as previous panel, but feature area of all SIRIUS predicted capsaicinoids is removed, Pearson = 0.71.

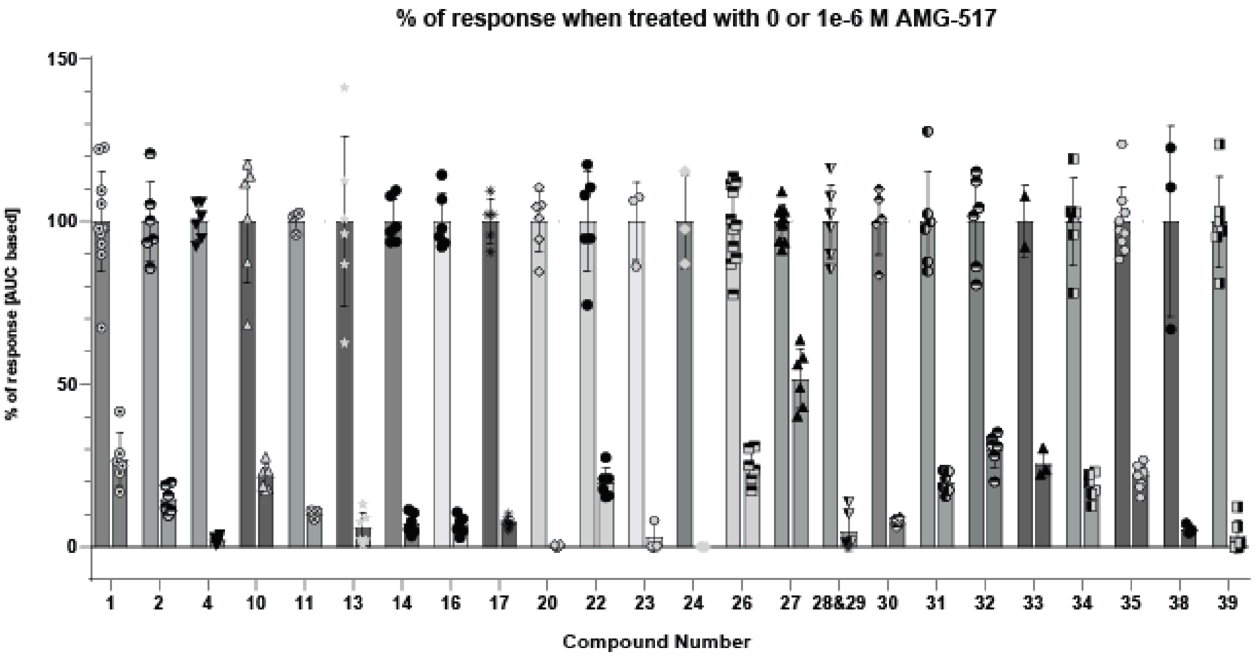


Fig. S6. Overview of inhibition of respective compounds by AMG-517. Compounds were added in a concentration close to their $EC_{50}$ along with 1.7 µM AMG-517 or without it to determine inhibition. The area under the peak was used to evaluate the inhibition. Some compounds showed greater inhibition than others. Because the respective capsaicinoid and the inhibitor were added simultaneously, it is possible that the capsaicinoid was outcompeted in a different way. This can reflect the different binding modes of the capsaicinoids.
