## Supplementary material for "Discovery and isolation of novel capsaicinoids and their TRPV1-related activity": SI_File_1_MSMS

### MSMS Annotation of Isolated Capsicinoids

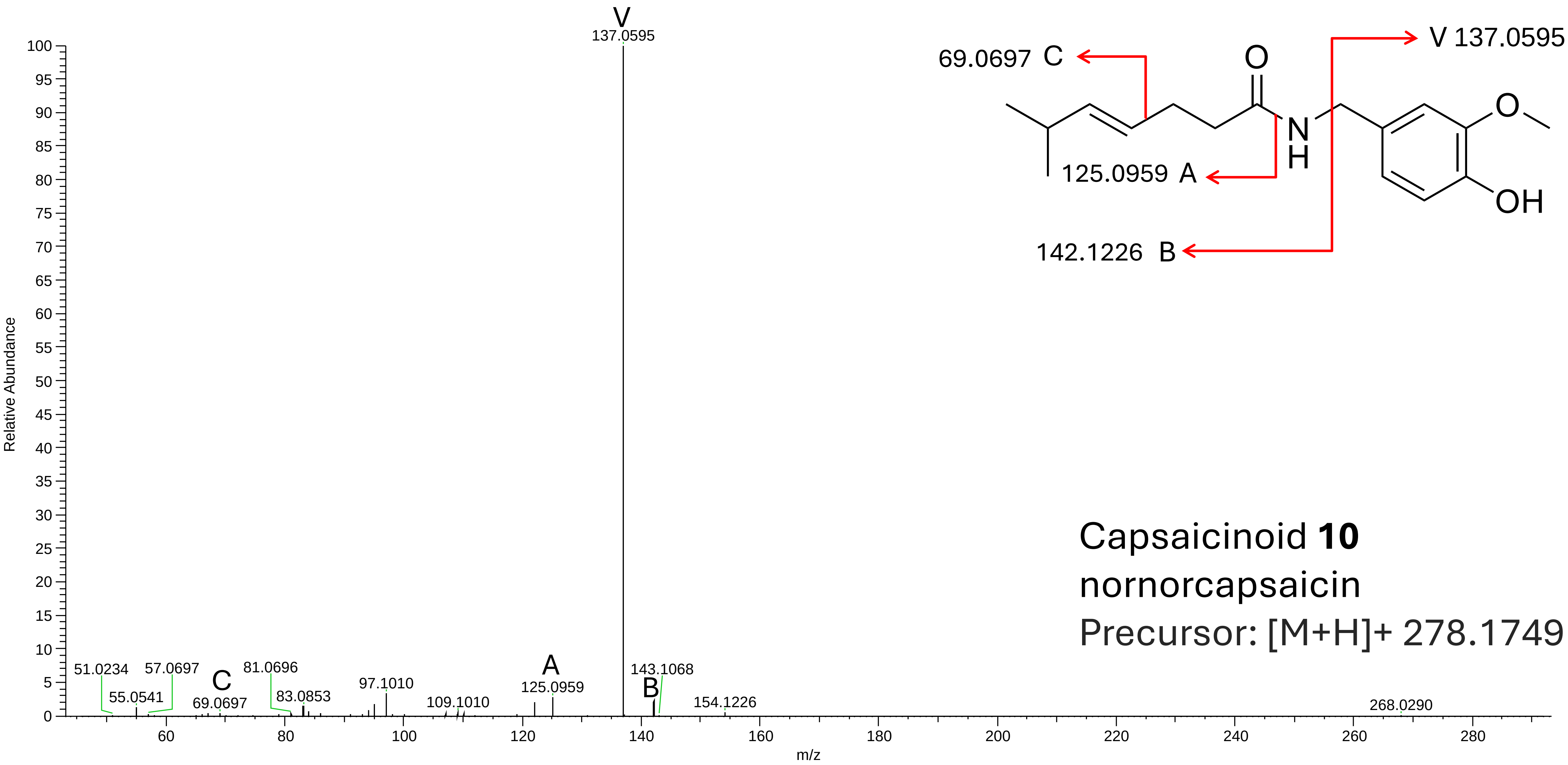
Capsaicinoid **10**
nornorcapsaicin
Precursor: [M+H]+ 278.1749
V 137.0595
69.0697 C
125.0959 A
142.1226 B
Relative Abundance
m/z
137.0595
51.0234
55.0541
57.0697
69.0697
81.0696
83.0853
97.1010
109.1010
125.0959
143.1068
154.1226
268.0290

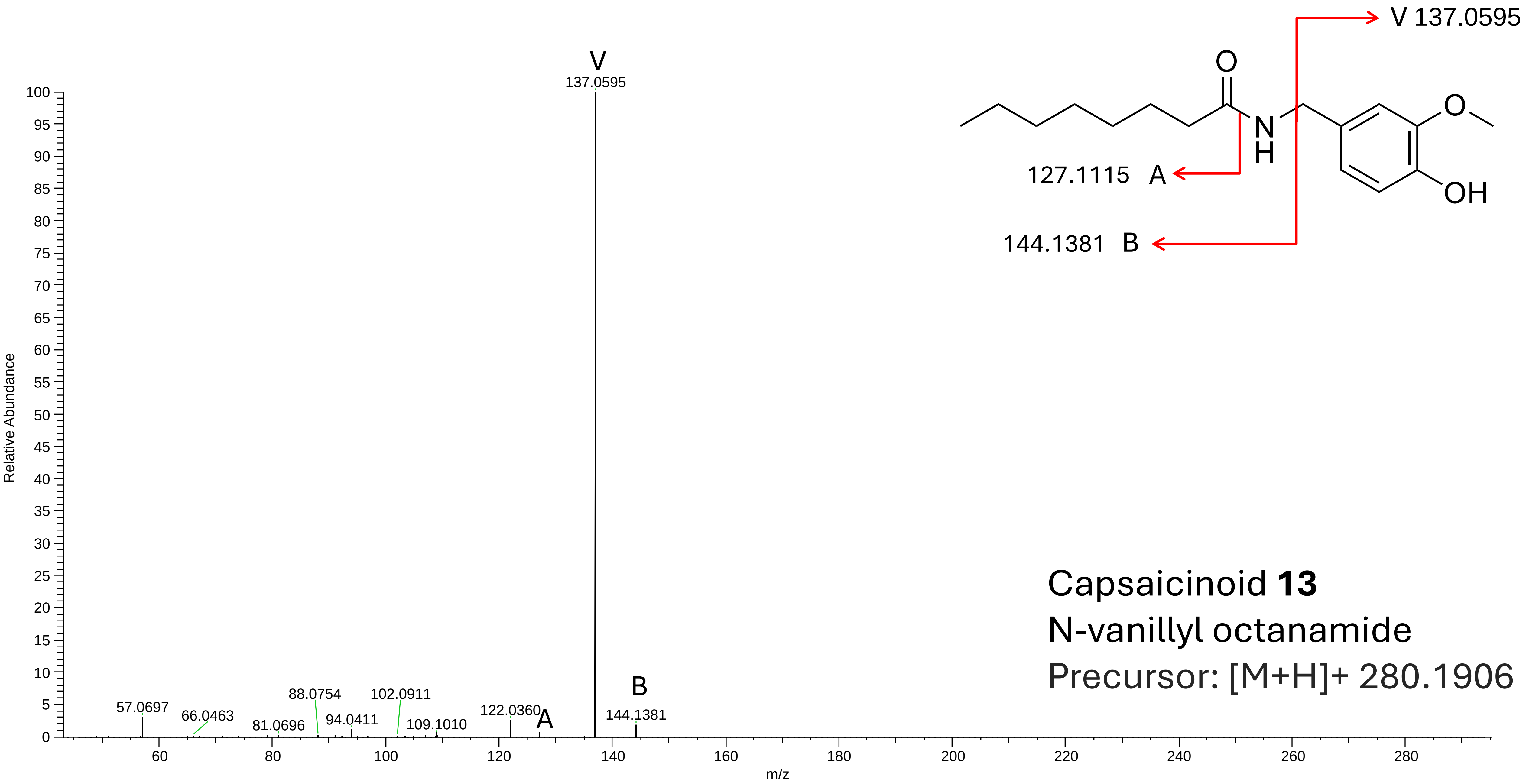


Capsaicinoid **13**

N-vanillyl octanamide

Precursor: [M+H]+ 280.1906

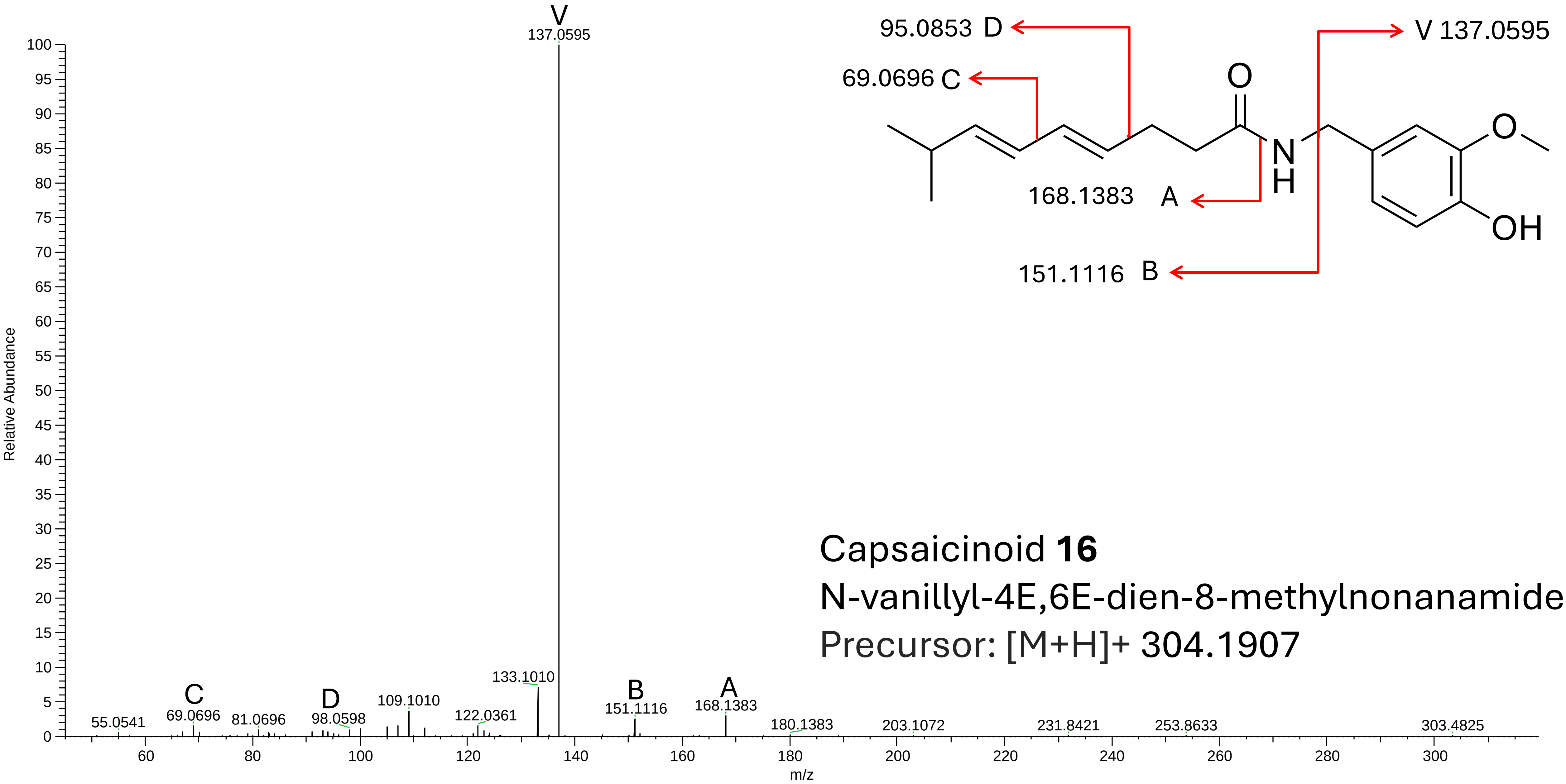


Capsaicinoid **16**

N-vanillyl-4E,6E-dien-8-methylnonanamide

Precursor: [M+H]+ 304.1907

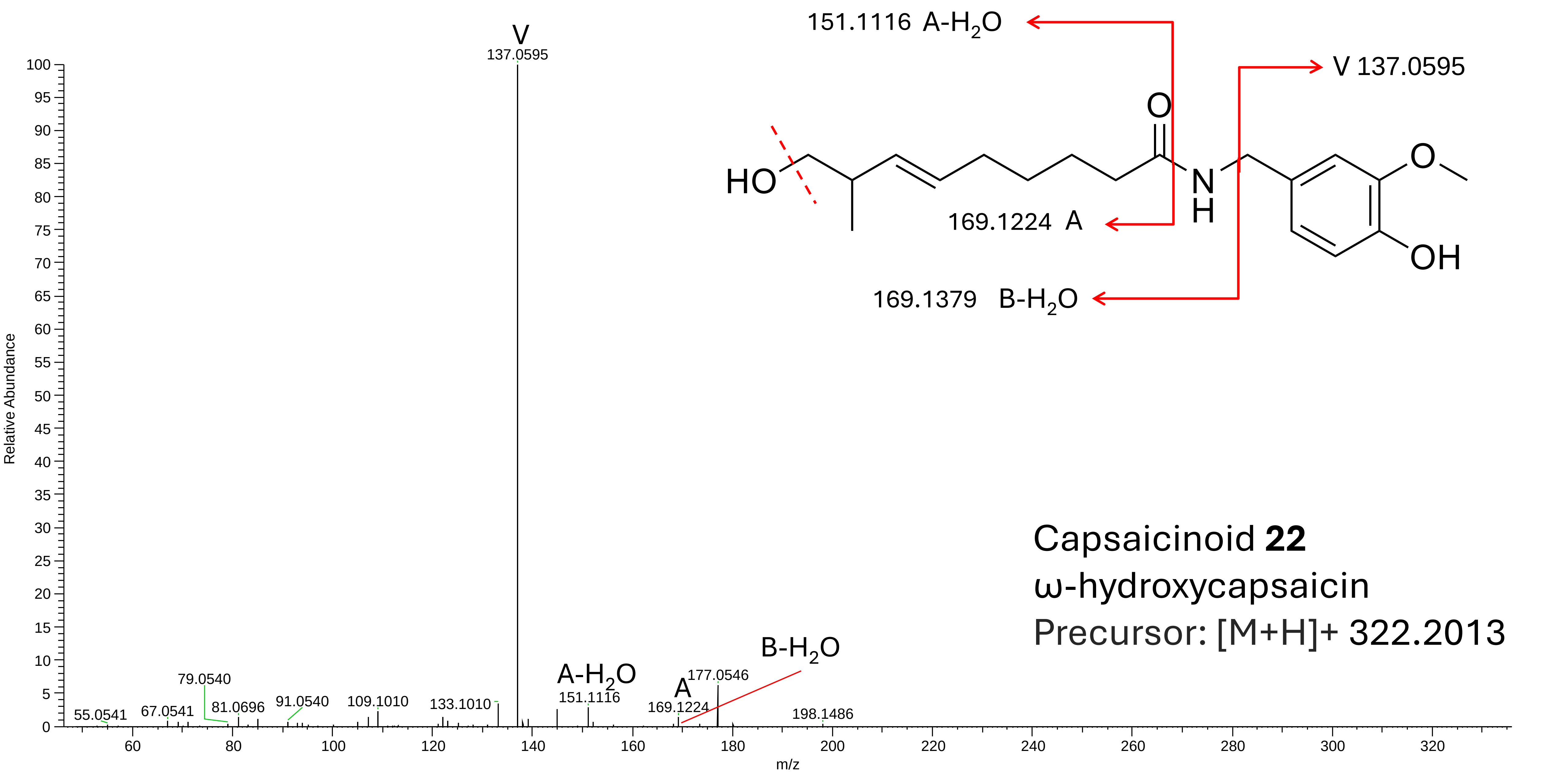
V
137.0595
151.1116 A-H2O
V 137.0595
169.1224 A
169.1379 B-H2O
HO
O
N
H
O
OH
Relative Abundance
m/z
55.0541
67.0541
79.0540
81.0696
91.0540
109.1010
133.1010
A-H2O
151.1116
A
169.1224
177.0546
B-H2O
198.1486
Capsaicinoid 22
ω-hydroxycapsaicin
Precursor: [M+H]+ 322.2013

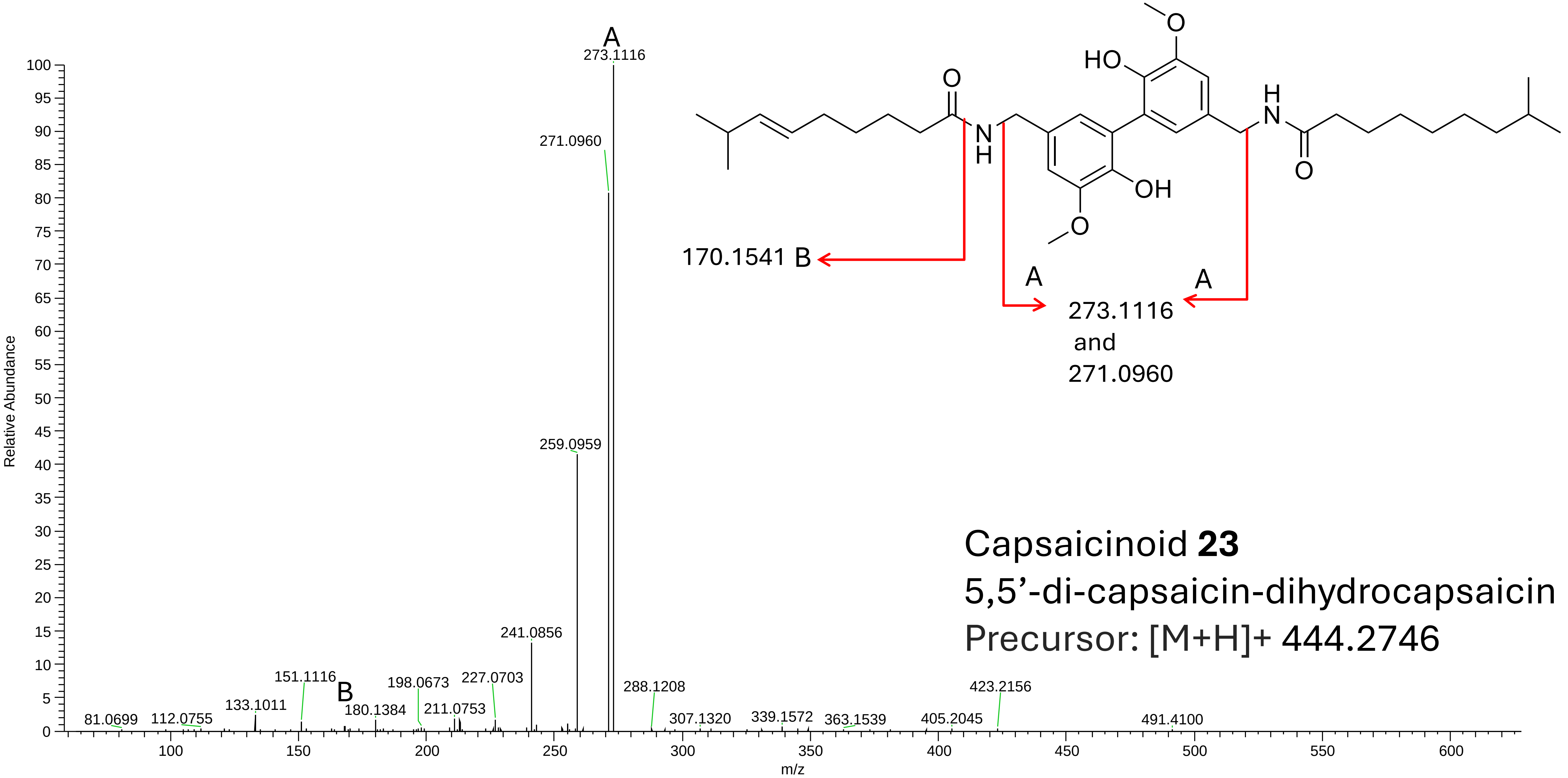


Capsaicinoid **23**
5,5’-di-capsaicin-dihydrocapsaicin
Precursor: [M+H]+ 444.2746

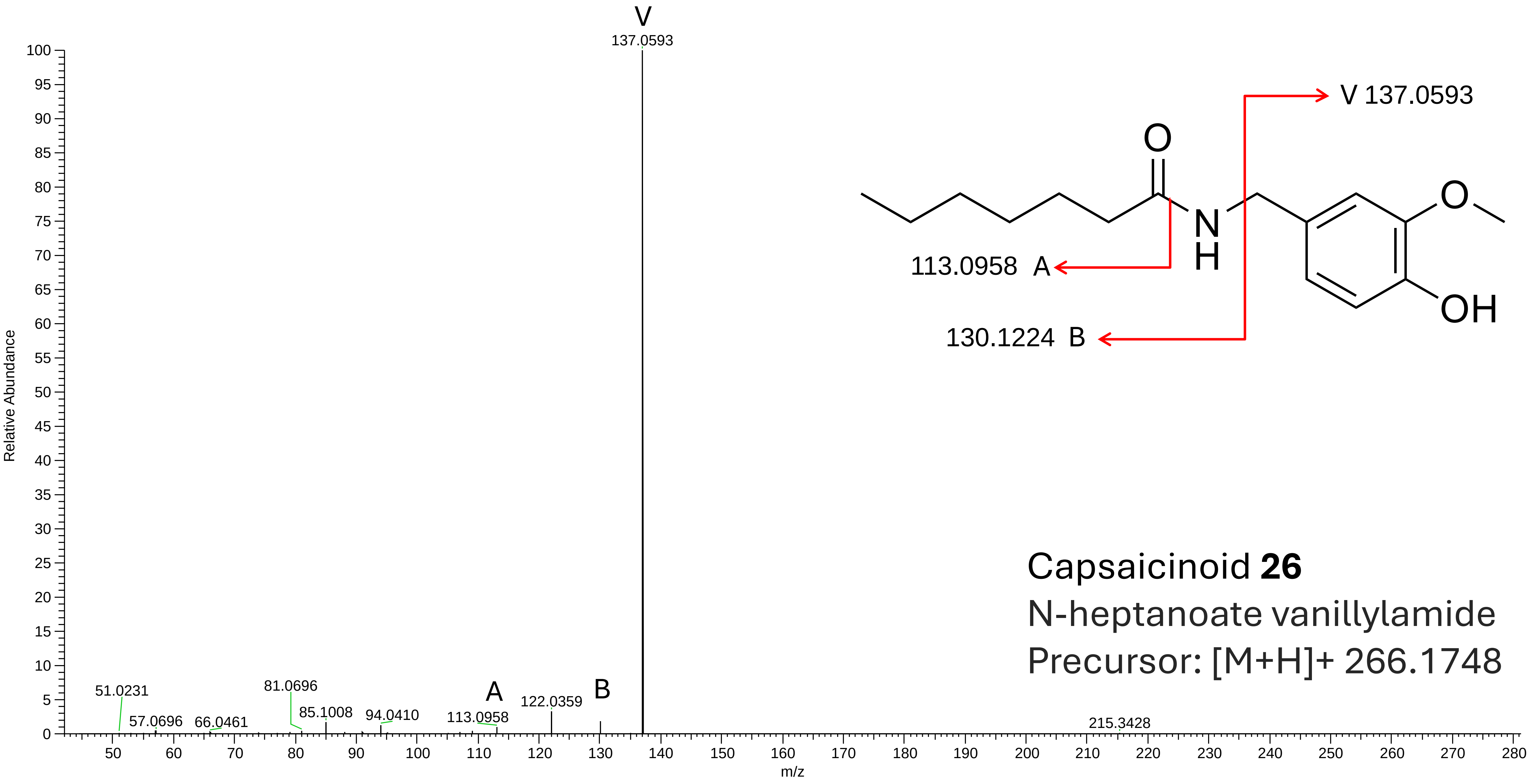
V
137.0593
Relative Abundance
m/z
51.0231
57.0696
66.0461
81.0696
85.1008
94.0410
A
113.0958
122.0359
B
215.3428
V 137.0593
O
N
H
O
OH
113.0958 A
130.1224 B
Capsaicinoid **26**
N-heptanoate vanillylamide
Precursor: [M+H]+ 266.1748

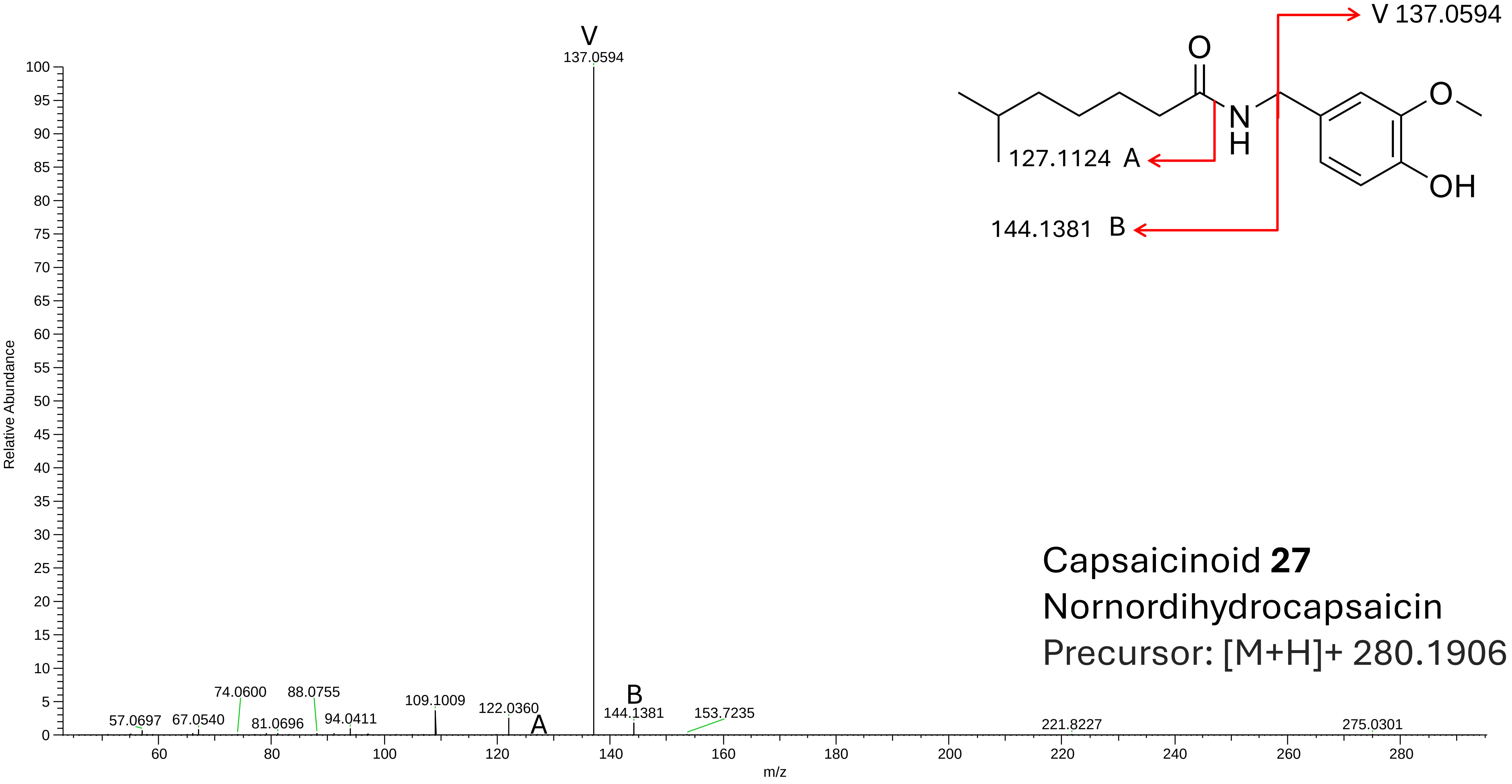
V
137.0594
100
95
90
85
80
75
70
65
60
55
50
45
40
35
30
25
20
15
10
5
0
Relative Abundance
57.0697
67.0540
74.0600
81.0696
88.0755
94.0411
109.1009
122.0360
A
B
144.1381
153.7235
221.8227
275.0301
60
80
100
120
140
160
180
200
220
240
260
280
m/z
V 137.0594
O
N
H
O
OH
127.1124 A
144.1381 B
Capsaicinoid **27**
Nornordihydrocapsaicin
Precursor: [M+H]+ 280.1906

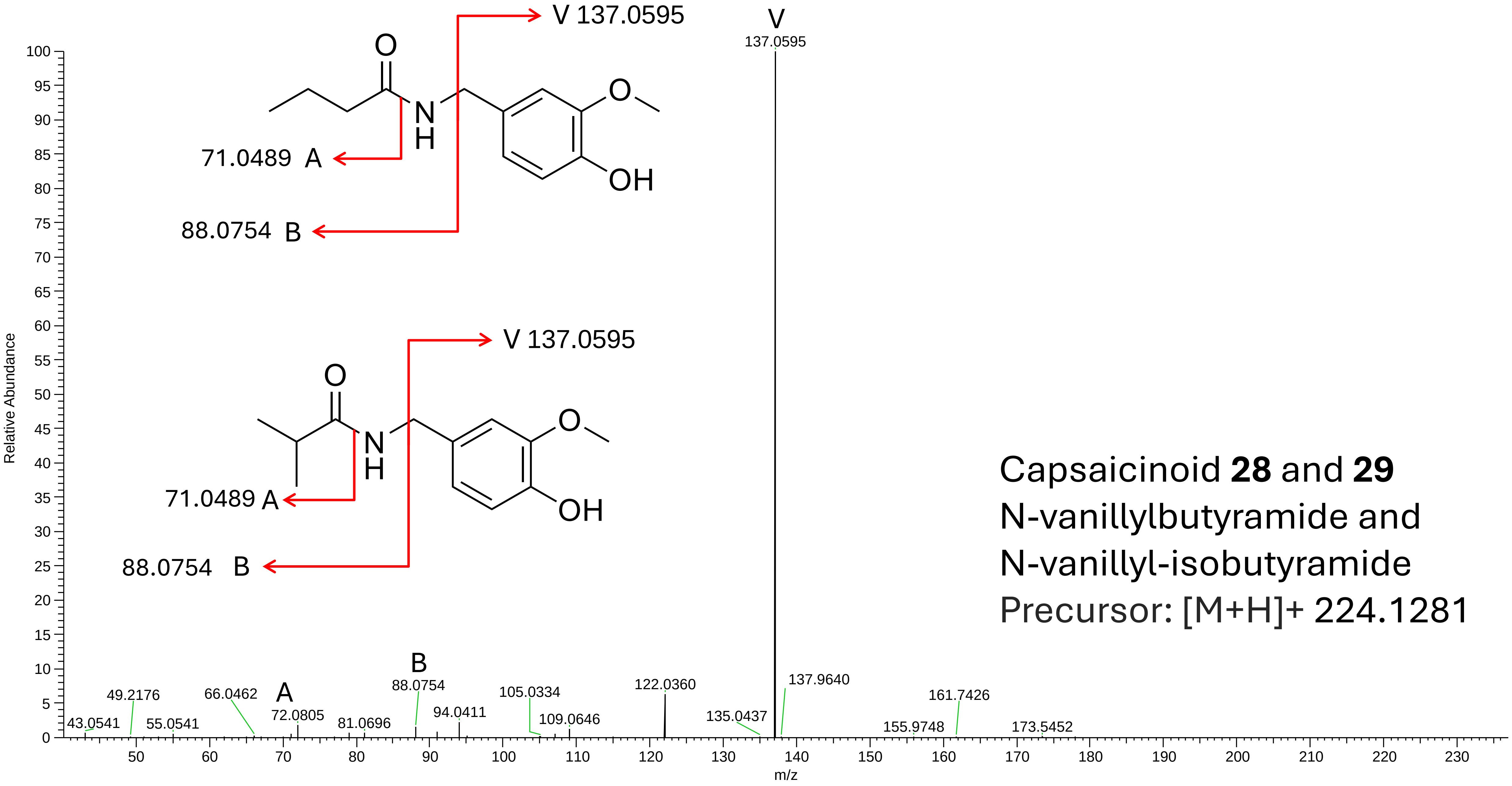


Capsaicinoid **28** and **29**
N-vanillylbutyramide and
N-vanillyl-isobutyramide
Precursor: [M+H]+ 224.1281

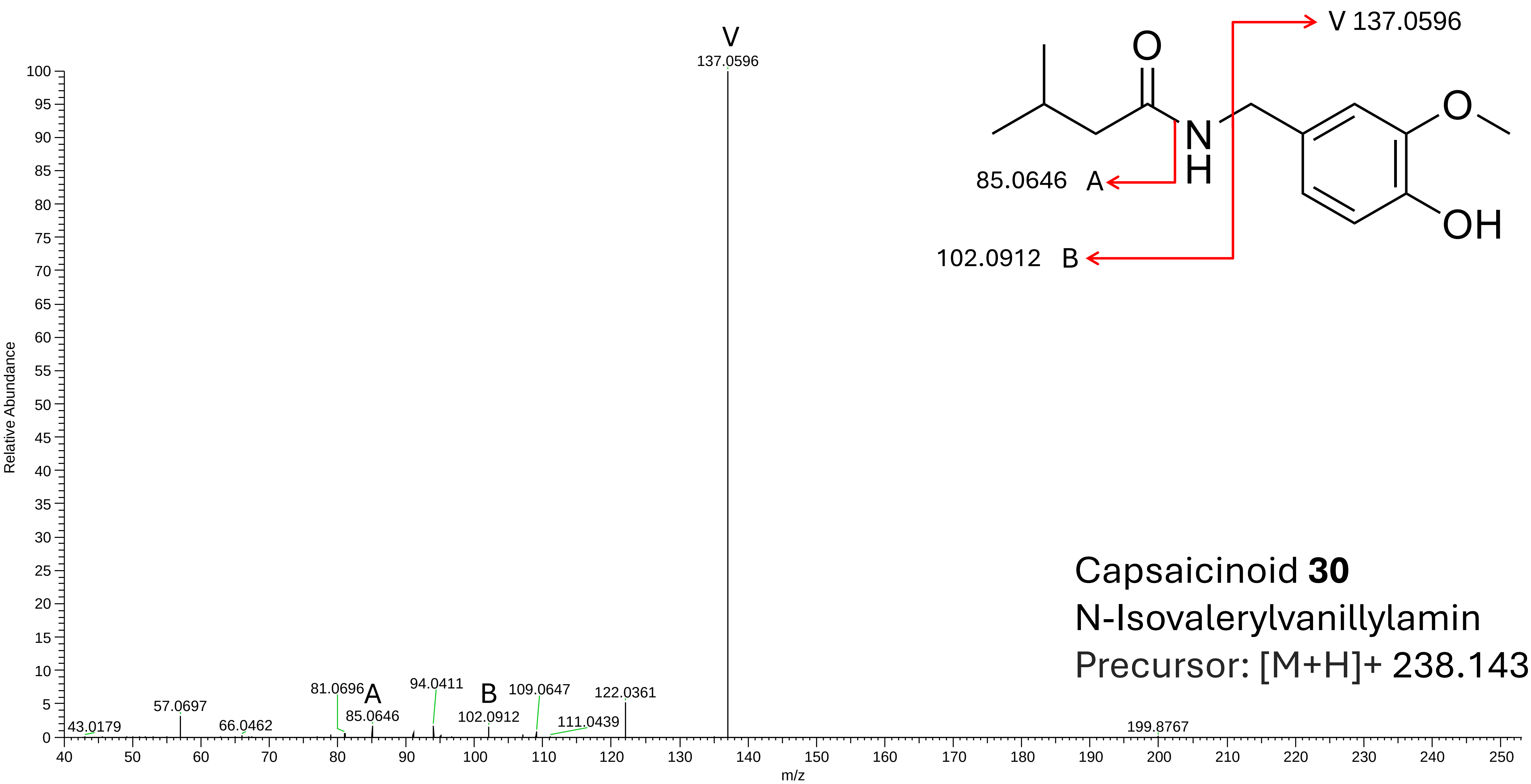


Capsaicinoid **30**
N-Isovalerylvanillylamin
Precursor: [M+H]+ 238.1437

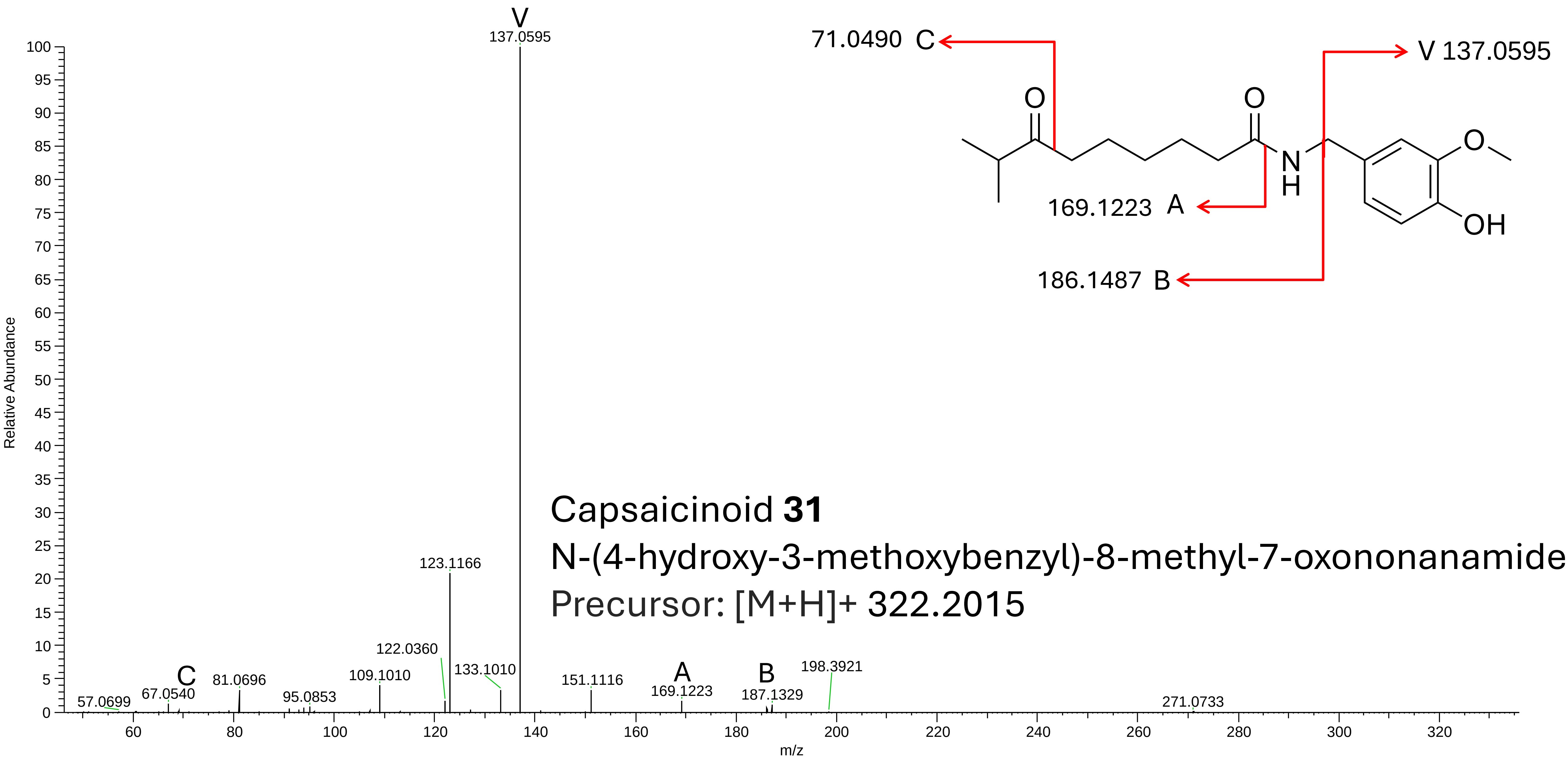


Capsaicinoid **31**

N-(4-hydroxy-3-methoxybenzyl)-8-methyl-7-oxononanamide

Precursor: [M+H]+ 322.2015

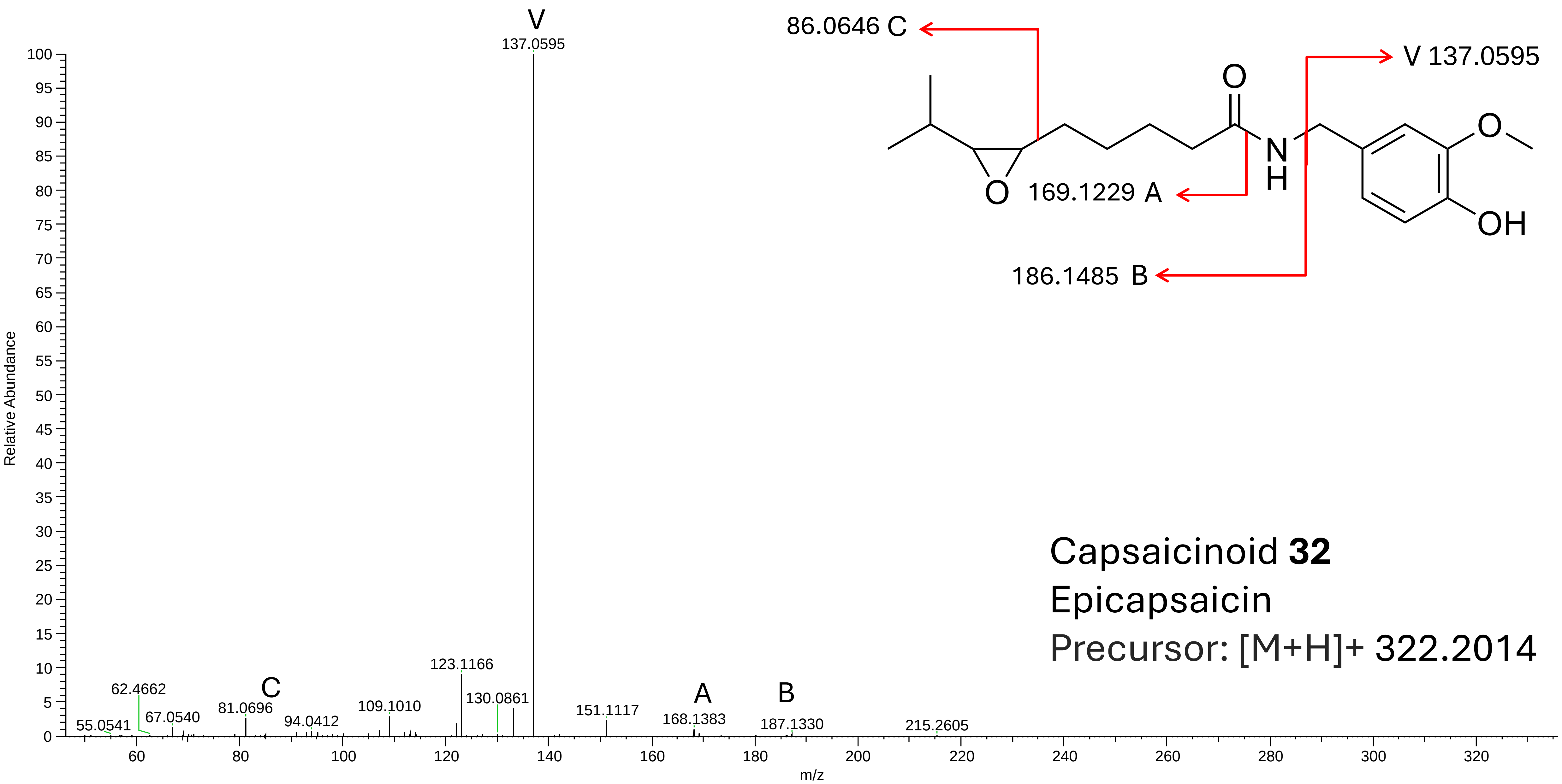

V
137.0595
123.1166
130.0861
62.4662
C
81.0696
109.1010
151.1117
A
B
55.0541
67.0540
94.0412
168.1383
187.1330
215.2605
Relative Abundance
m/z
86.0646 C
V 137.0595
169.1229 A
186.1485 B
O
N
H
OH
Capsaicinoid 32
Epicapsaicin
Precursor: [M+H]+ 322.2014

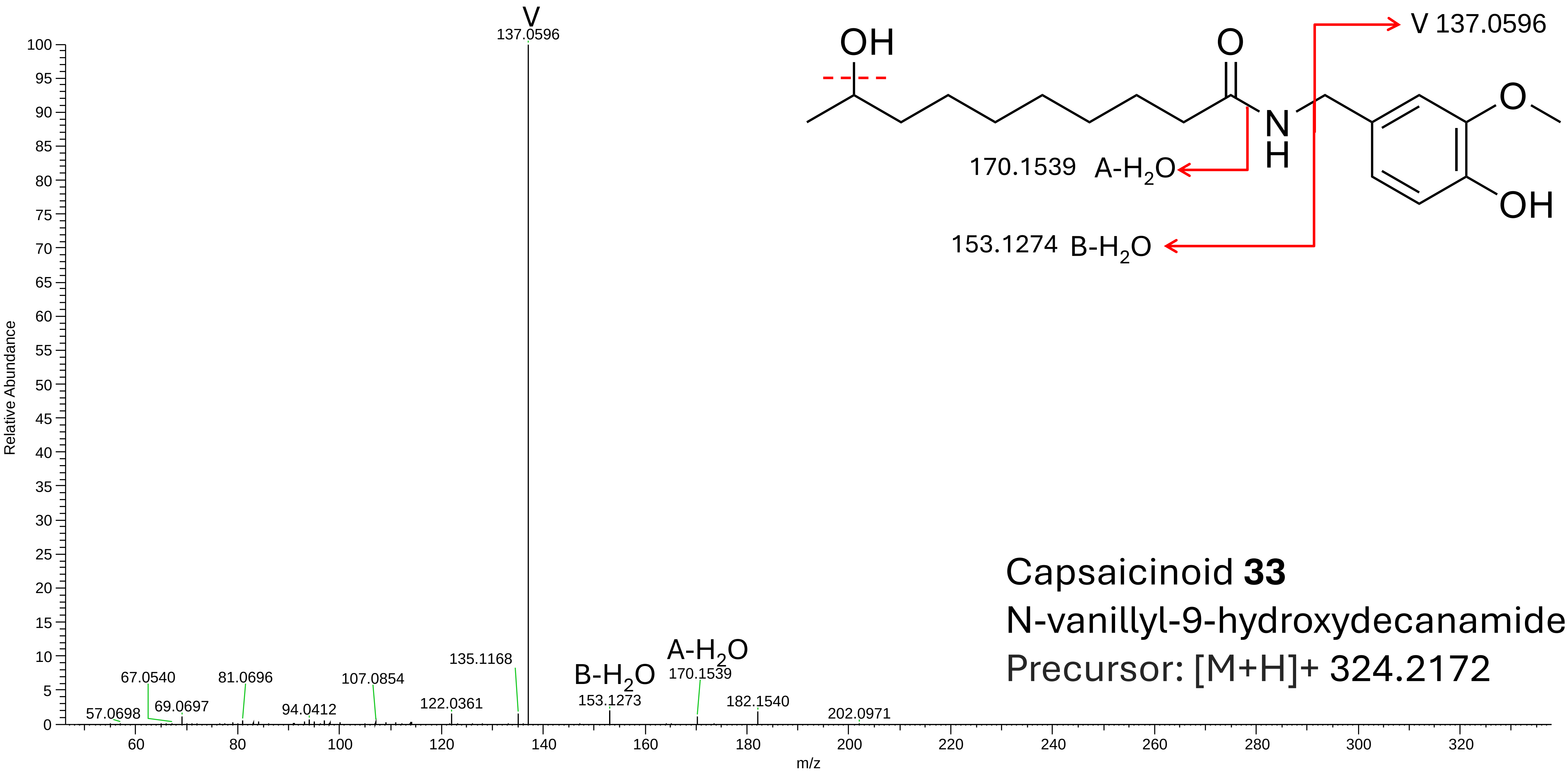


Capsaicinoid **33**
N-vanillyl-9-hydroxydecanamide
Precursor: [M+H]+ 324.2172

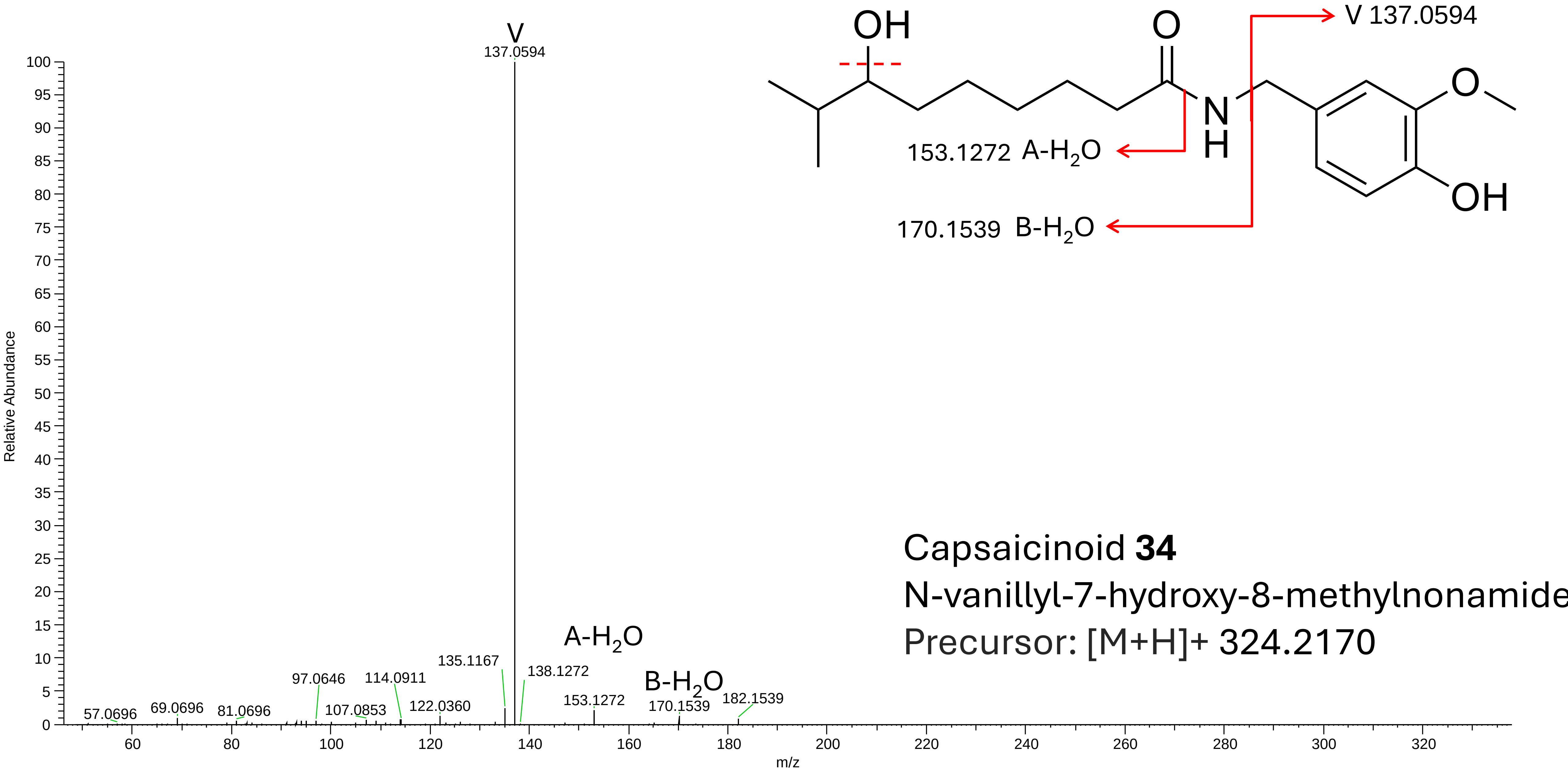
V
137.0594
Relative Abundance
m/z
57.0696
69.0696
81.0696
97.0646
107.0853
114.0911
122.0360
135.1167
138.1272
153.1272
170.1539
182.1539
A-H2O
B-H2O
OH
O
N
H
OH
O
V 137.0594
153.1272 A-H2O
170.1539 B-H2O
Capsaicinoid 34
N-vanillyl-7-hydroxy-8-methylnonamide
Precursor: [M+H]+ 324.2170

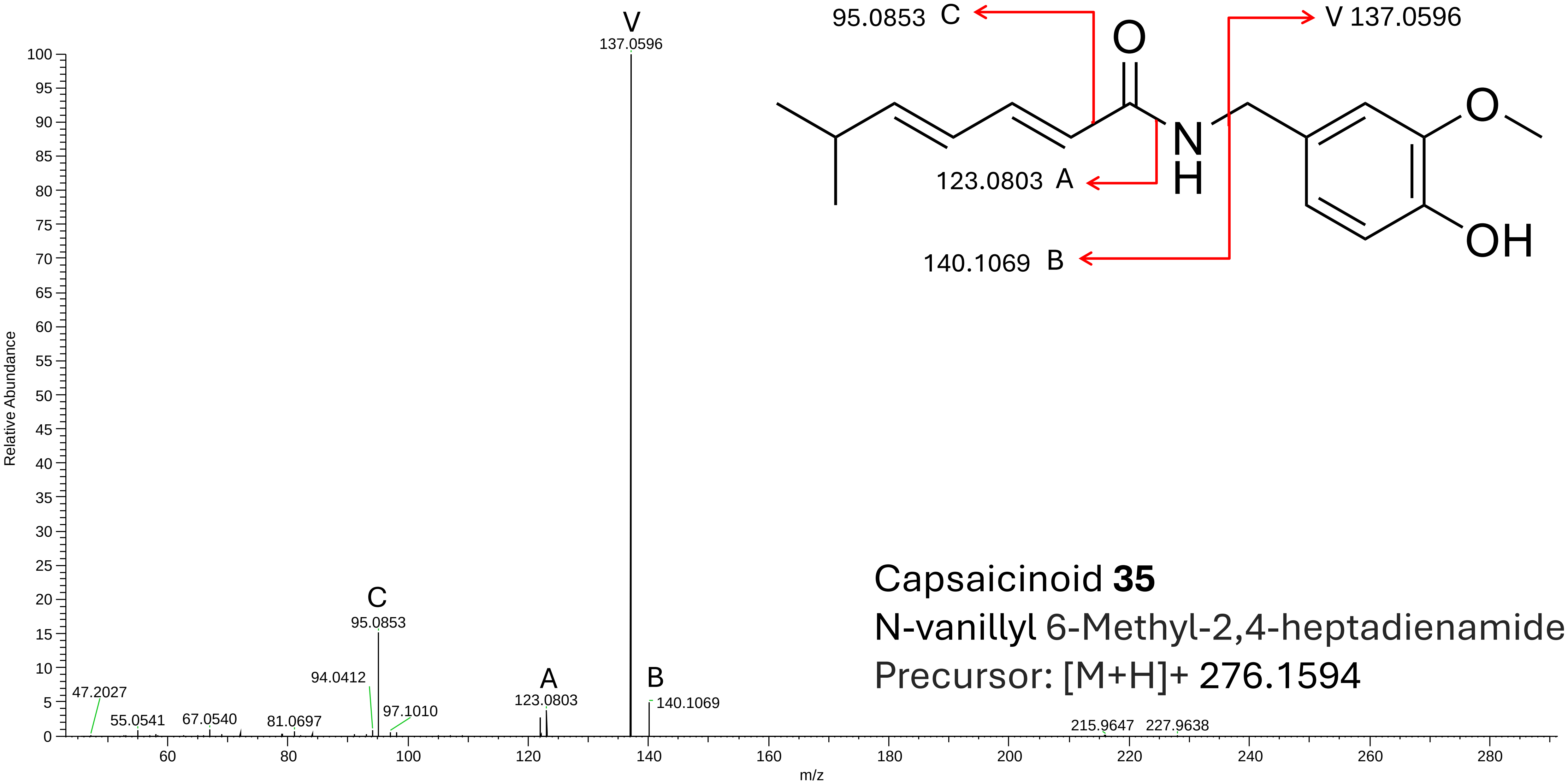


Capsaicinoid **35**

N-vanillyl 6-Methyl-2,4-heptadienamide

Precursor: [M+H]+ 276.1594

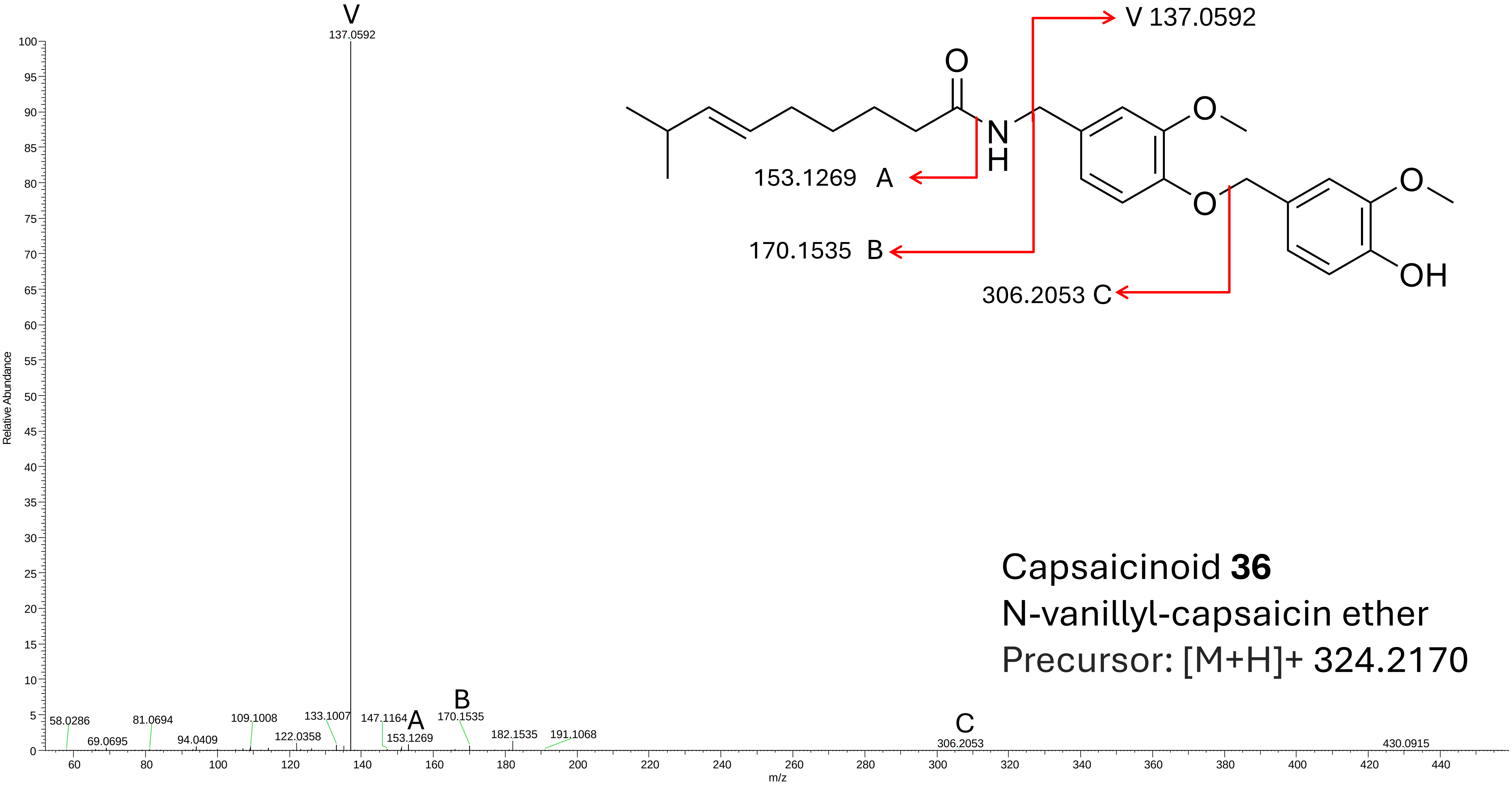


Capsaicinoid **36**
N-vanillyl-capsaicin ether
Precursor: [M+H]+ 324.2170

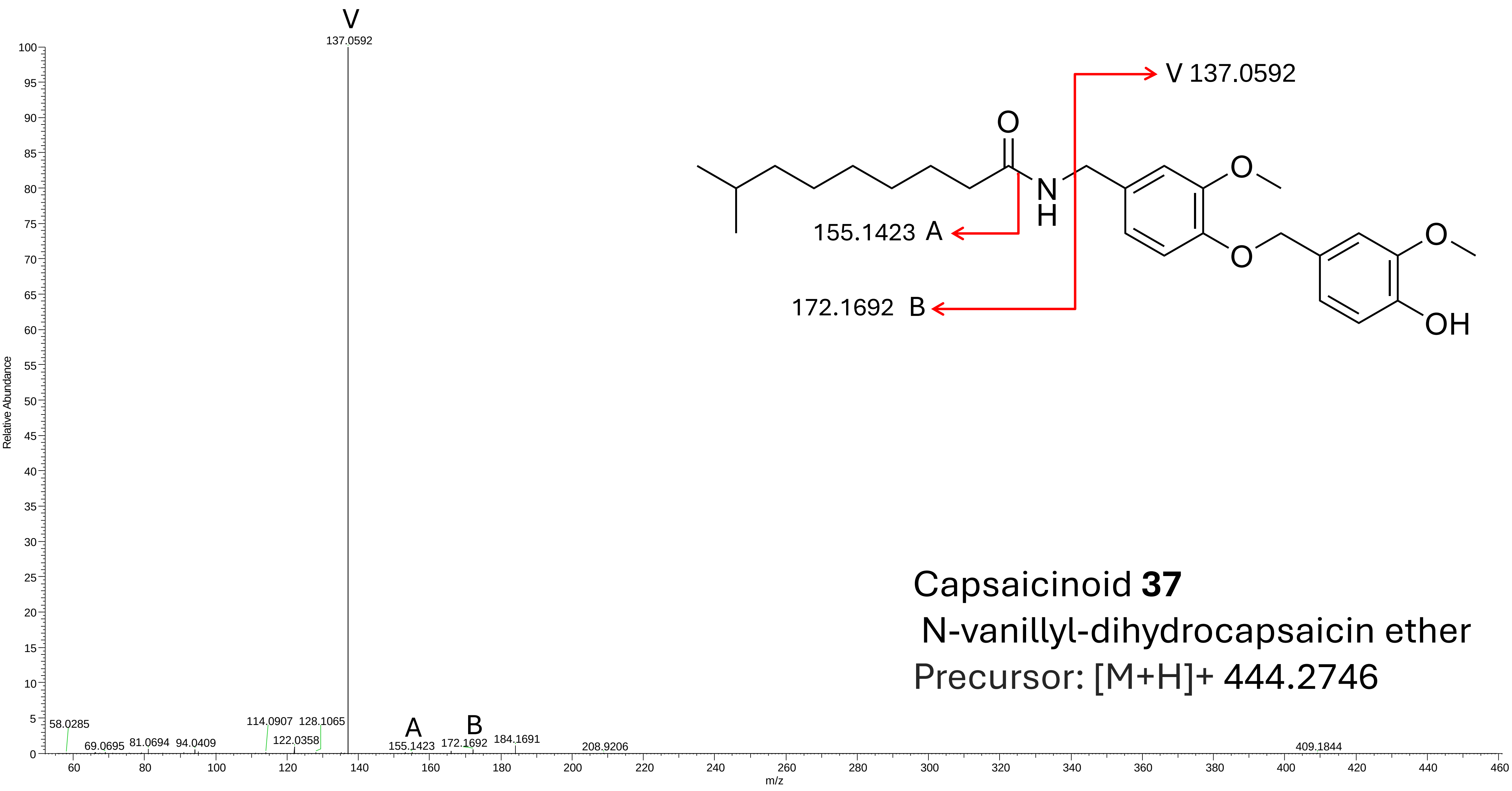


Capsaicinoid **37**

N-vanillyl-dihydrocapsaicin ether

Precursor: [M+H]+ 444.2746

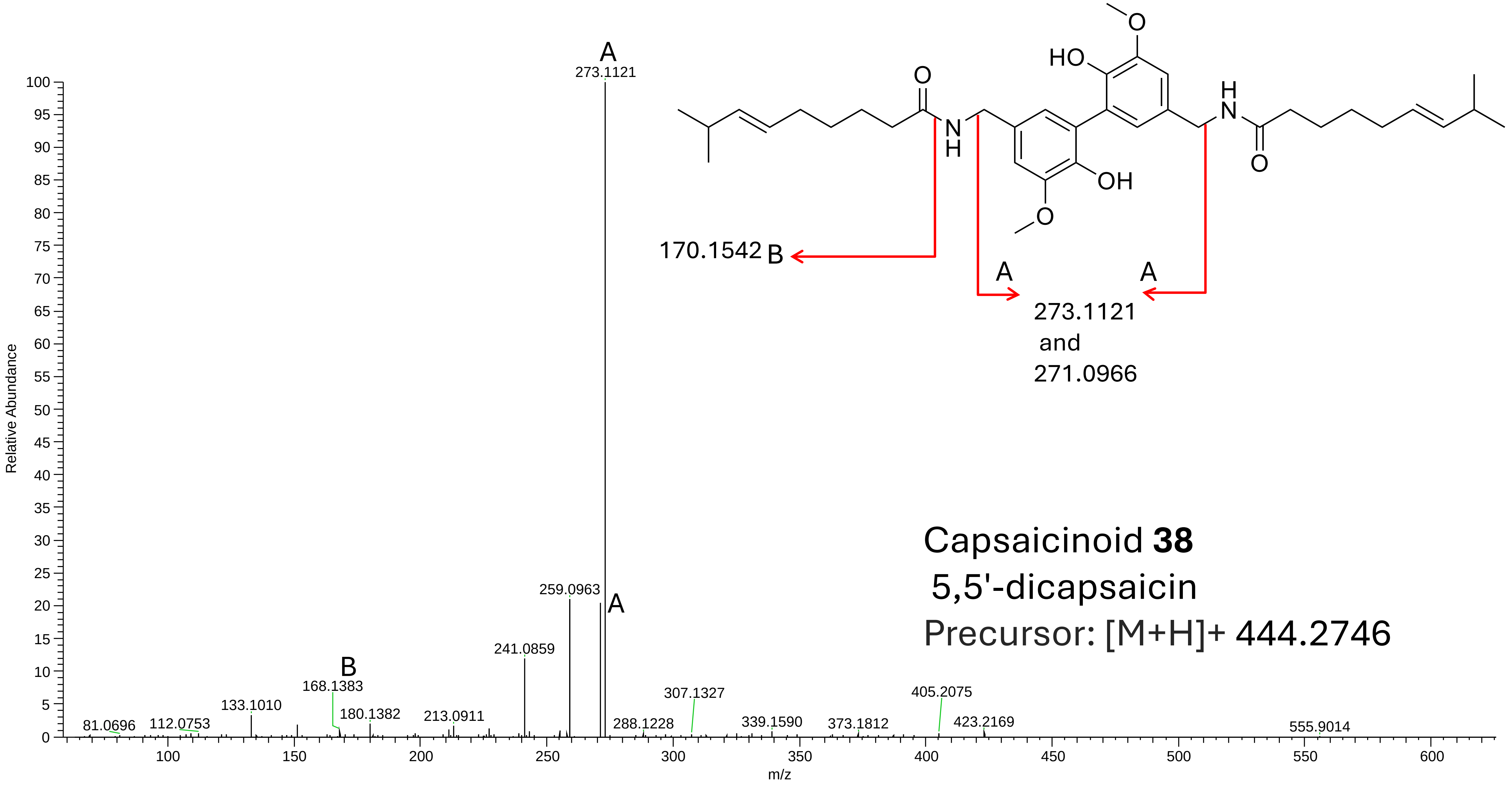
Capsaicinoid **38**
5,5'-dicapsaicin
Precursor: [M+H]+ 444.2746
170.1542 B
A
A
273.1121
and
271.0966
HO
OH
O
N
H
Relative Abundance
m/z
273.1121
259.0963
241.0859
168.1383
133.1010
180.1382
213.0911
112.0753
81.0696
288.1228
307.1327
339.1590
373.1812
405.2075
423.2169
555.9014
