## Supplementary material for "Discovery and isolation of novel capsaicinoids and their TRPV1-related activity": SI_File_2_NMR

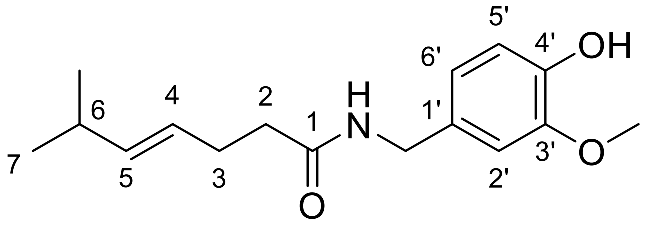

Nornorcapsaicin **10**, Capsicum8-P4,P5

$^{1}$H NMR (500.0 MHz, $CD_3CN$) $\delta$ = 6.86 (d, 1H, $J_{2',6'}$ = 1.9, H2′), 6.74 (d, 1H, $J_{5',6'}$ = 8.0, H5′), 6.71 (ddt, 1H, $J_{6',5'}$ = 8.0, $J_{6',2'}$ = 1.9, $J_{6',CH2}$ = 0.7, H6′), 6.65 (bs, 1H, NH), 6.42 (bs, 1H, OH), 5.34–5.47 (m, 2H, H4 and H5), 4.22 (d, 2H, $J_{CH2,NH}$ = 6.0, 1′–C**H₂**), 3.82 (s, 3H, $OCH_3$), 2.16–2.26 (m, 5H, H2, H3 and H6).

$^{13}$C NMR (125.7 MHz, $CD_3CN$) $\delta$ = 172.8 (C1), 148.1 (C3′), 146.1 (C4′), 139.2 (C5), 132.3 (C1′), 126.7 (C4), 121.0 (C6′), 115.4 (C5′), 112.1 (C2′), 56.6 ($OCH_3$), 43.3 (1′–C**H₂**), 37.0 (C2), 31.7 (C6), 29.4 (C3), 22.8 (C7).

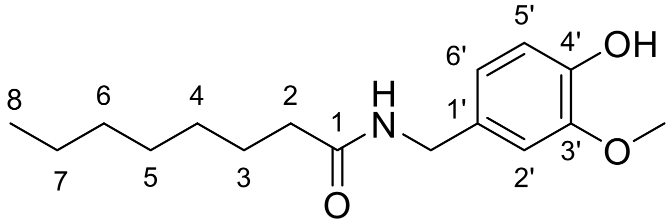

**13**, Capsicum8-P10

$^{1}$H NMR (500.0 MHz, $CD_3CN$) $\delta$ = 6.85 (d, 1H, $J_{2',6'}$ = 1.9, H2′), 6.75 (d, 1H, $J_{5',6'}$ = 7.9, H5′), 6.71 (ddt, 1H, $J_{6',5'}$ = 8.0, $J_{6',2'}$ = 1.9, $J_{6',CH2}$ = 0.7, H6′), 6.65 (bs, 1H, NH), 6.42 (bs, 1H, OH), 4.22 (d, 2H, $J_{CH2,NH}$ = 6.0, 1′–C**H₂**), 3.82 (s, 3H, $OCH_3$), 2.13 (m, 2H, H2), 1.57 (m, 2H, H3), 1.21–1.33 (m, 8H, H4, H5, H6 and H7), 0.88 (m, 3H, H8).

$^{13}$C NMR (125.7 MHz, $CD_3CN$) $\delta$ = 173.6 (C1), 148.1 (C3′), 146.1 (C4′), 132.4 (C1′), 121.0 (C6′), 115.4 (C5′), 112.0 (C2′), 56.6 ($OCH_3$), 43.2 (1′–C**H₂**), 36.9 (C2), 32.5 (C6), 29.9 (C4 or C5), 29.8 (C4 or C5), 26.6 (C3), 23.3 (C7), 14.4 (C8).

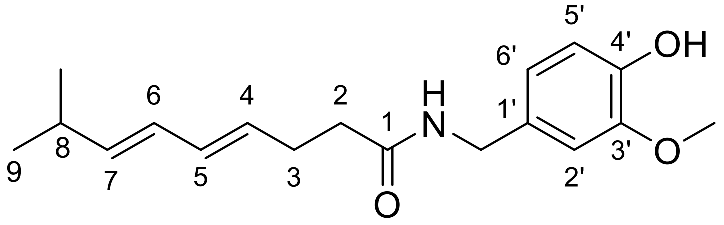

**16**, Capsicum8-P14

$^{1}$H NMR (500.0 MHz, $CD_3CN$) $\delta$ = 6.85 (d, 1H, $J_{2',6'}$ = 2.0, H2′), 6.74 (d, 1H, $J_{5',6'}$ = 8.0, H5′), 6.70 (ddt, 1H, $J_{6',5'}$ = 8.0, $J_{6',2'}$ = 1.9, $J_{6',CH2}$ = 0.7, H6′), 6.67 (bs, 1H, NH), 6.44 (bs, 1H, OH), 5.93–6.05 (m, 2H, H5 and H6), 5.53–5.60 (m, 2H, H4 and H7), 4.22 (d, 2H, $J_{CH2,NH}$ = 6.0, 1′–C**H₂**), 3.82 (s, 3H, $OCH_3$), 2.20–2.35 (m, 5H, H2, H3 and H8), 0.98 (d, 6H, $J_{9,8}$ = 6.7, H9).

$^{13}$C NMR (125.7 MHz, $CD_3CN$) $\delta$ = 172.8 (C1), 148.1 (C3′), 146.1 (C4′), 140.8 (C7), 132.3 (C1′), 132.1 (C5), 131.7 (C4), 128.3 (C6), 121.0 (C6′), 115.4 (C5′), 112.1 (C2′), 56.6 ($OCH_3$), 43.3 (1′–C**H₂**), 36.5 (C2), 31.8 (C8), 29.3 (C3), 22.6 (C9).

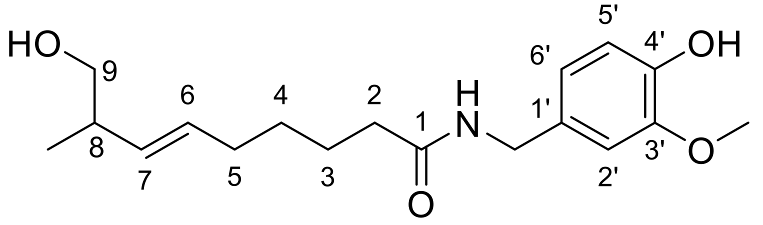

ω-hydroxycapsaicin **22**, Capsicum11-P8

$^1$H NMR (500.0 MHz, $CD_3CN$) $\delta$ = 6.85 (d, 1H, $J_{2',6'}$ = 2.0, H2′), 6.75 (d, 1H, $J_{5',6'}$ = 8.1, H5′), 6.70 (ddt, 1H, $J_{6',5'}$ = 8.1, $J_{6',2'}$ = 2.0, $J_{6',CH2}$ = 0.7, H6′), 6.69 (bs, 1H, NH), 6.44 (bs, 1H, 4′–OH), 5.44 (dtd, 1H, $J_{6,7}$ = 15.4, $J_{6,5}$ = 6.7, $J_{6,8}$ = 1.0, H6), 5.32 (ddt, 1H, $J_{7,6}$ = 15.4, $J_{7,8}$ = 7.4, $J_{6,5}$ = 1.3, H7), 4.22 (d, 2H, $J_{CH2,NH}$ = 6.0, 1′–C**H₂**), 3.82 (s, 3H, $OCH_3$), 3.26–3.34 (m, 2H, H9), 2.56 (t, 1H, $J_{OH,9}$ = 5.9, 9-O**H**), 2.11–2.22 (m, 3H, H2 and H8), 2.00 (m, 2H, H5), 1.57 (m, 2H, H3), 1.35 (m, 2H, H4), 0.93 (d, 3H, $J_{CH3,8}$ = 6.8, 8-C**H₃**).

$^{13}$C NMR (125.7 MHz, $CD_3CN$) $\delta$ = 173.6 (C1), 148.2 (C3′), 146.0 (C4′), 134.4 (C7), 132.4 (C1′), 130.9 (C6), 121.0 (C6′), 115.4 (C5′), 112.0 (C2′), 67.82 (C9), 56.6 ($OCH_3$), 43.2 (1′–C**H₂**), 40.4 (C8), 36.6 (C2), 33.0 (C5), 29.7 (C4), 26.0 (C3), 17.1 (8-**C**H₃).

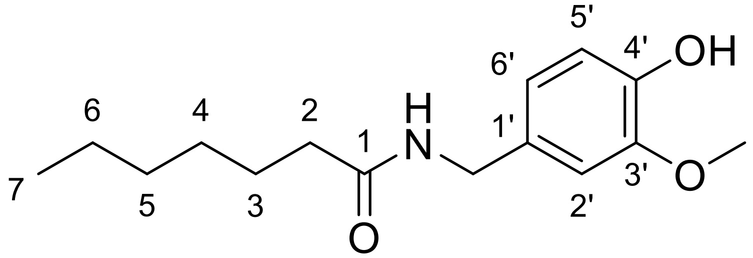

**26**, Capsicum10-P3

$^1$H NMR (500.0 MHz, $CD_3CN$) $\delta$ = 6.85 (d, 1H, $J_{2',6'}$ = 2.0, H2′), 6.75 (d, 1H, $J_{5',6'}$ = 8.0, H5′), 6.70 (ddt, 1H, $J_{6',5'}$ = 7.9, $J_{6',2'}$ = 1.9, $J_{6',CH2}$ = 0.7, H6′), 6.65 (bs, 1H, NH), 6.41 (bs, 1H, OH), 4.22 (d, 2H, $J_{CH2,NH}$ = 6.0, 1′–C**H₂**), 3.82 (s, 3H, $OCH_3$), 2.13 (m, 2H, H2), 1.56 (m, 2H, H3), 1.26–1.33 (m, 6H, H4, H5 and H6), 0.88 (m, 3H, H7).

Small amount of the sample did not allow for $^{13}$C NMR measurement.

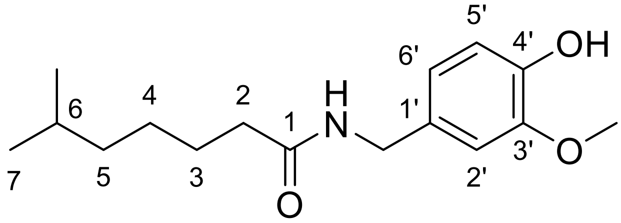

**27**, Capsicum8-P9

$^1$H NMR (500.0 MHz, $CD_3CN$) $\delta$ = 6.86 (d, 1H, $J_{2',6'}$ = 2.0, H2′), 6.74 (d, 1H, $J_{5',6'}$ = 8.1, H5′), 6.71 (ddt, 1H, $J_{6',5'}$ = 8.0, $J_{6',2'}$ = 1.9, $J_{6',CH2}$ = 0.6, H6′), 6.65 (bs, 1H, NH), 6.41 (bs, 1H, OH), 4.22 (d, 2H, $J_{CH2,NH}$ = 6.0, 1′–C**H₂**), 3.82 (s, 3H, $OCH_3$), 2.13 (m, 2H, H2), 1.47–1.60 (m, 3H, H3 and H6), 1.29 (m, 2H, H4), 1.18 (m, 2H, H5), 0.86 (d, 6H, $J_{7,6}$ = 6.7, H7).

$^{13}$C NMR (125.7 MHz, $CD_3CN$) $\delta$ = 173.7 (C1), 148.2 (C3′), 132.4 (C1′), 121.0 (C6′), 115.4 (C5′), 112.0 (C2′), 56.6 ($OCH_3$), 43.2 (1′–C**H₂**), 39.4 (C5), 36.9 (C2), 28.6 (C6), 27.7 (C4), 26.8 (C3), 22.9 (C7).

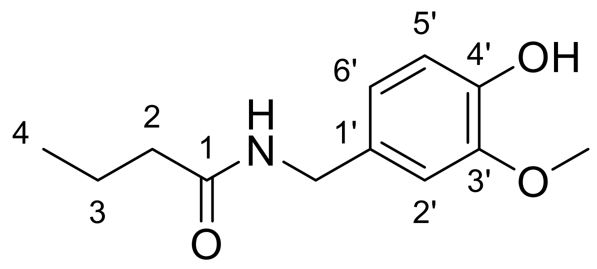

**28**, Capsicum10-P3

Structural analysis performed for a mixture of compounds **28** and **29**.

$^{1}$H NMR (500.0 MHz, $CD_3CN$) $\delta$ = 6.86 (d, 1H, $J_{2',6'}$ = 1.9, H2′), 6.75 (d, 1H, $J_{5',6'}$ = 8.1, H5′), 6.69–6.72 (m, 1H, H6′), 6.64 (bs, 1H, NH), 6.41 (bs, 1H, OH), 4.23 (d, 2H, $J_{CH2,NH}$ = 6.0, 1′–C**H₂**), 3.82 (s, 3H, $OCH_3$), 2.13 (m, 2H, H2), 1.60 (sextet, 2H, $J_{3,2}$ = 7.4, H3), 0.91 (t, 3H, $J_{4,3}$ = 7.4, H4).

$^{13}$C NMR (125.7 MHz, $CD_3CN$) $\delta$ = 173.5 (C1), 148.3 (C3′), 146.1 (C4′), 132.5 (C1′), 121.0 (C6′), 115.4 (C5′), 112.0 (C2′), 56.6 ($OCH_3$), 43.2 (1′–C**H₂**), 38.8 (C2), 19.9 (C3), 14.0 (C4).

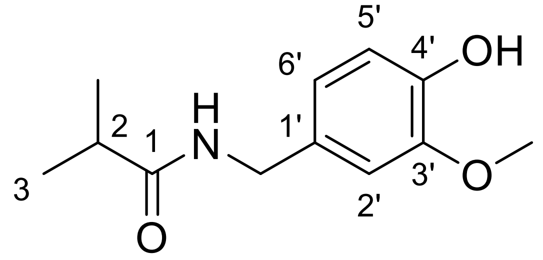

**29**, Capsicum10-P3

Structural analysis performed for a mixture of compounds **28** and **29**.

$^{1}$H NMR (500.0 MHz, $CD_3CN$) $\delta$ = 6.85 (d, 1H, $J_{2',6'}$ = 1.9, H2′), 6.75 (d, 1H, $J_{5',6'}$ = 8.1, H5′), 6.69–6.72 (m, 1H, H6′), 6.64 (bs, 1H, NH), 6.41 (bs, 1H, OH), 4.23 (d, 2H, $J_{CH2,NH}$ = 6.0, 1′–C**H₂**), 3.82 (s, 3H, $OCH_3$), 2.37 (septet, 1H, $J_{2,3}$ = 6.8, H2), 1.08 (d, 6H, $J_{3,2}$ = 6.9, H3).

$^{13}$C NMR (125.7 MHz, $CD_3CN$) $\delta$ = 173.5 (C1), 148.3 (C3′), 146.1 (C4′), 132.5 (C1′), 120.8 (C6′), 115.4 (C5′), 111.9 (C2′), 56.6 ($OCH_3$), 43.1 (1′–C**H₂**), 35.9 (C2), 19.9 (C3).

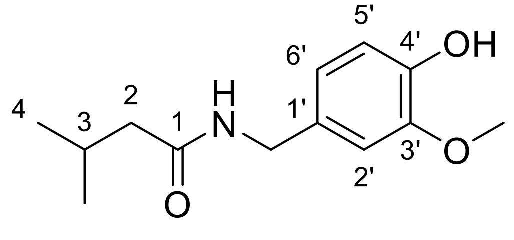

**30**, Capsicum10-P6

$^{1}$H NMR (500.0 MHz, $CD_3CN$) $\delta$ = 6.86 (d, 1H, $J_{2',6'}$ = 1.9, H2′), 6.75 (d, 1H, $J_{5',6'}$ = 8.1, H5′), 6.71 (ddt, 1H, $J_{6',5'}$ = 8.1, $J_{6',2'}$ = 2.0, $J_{6',CH2}$ = 0.7, H6′), 6.66 (bs, 1H, NH), 6.41 (bs, 1H, OH), 4.23 (d, 2H, $J_{CH2,NH}$ = 5.9, 1′–C**H₂**), 3.82 (s, 3H, $OCH_3$), 2.00–2.04 (m, 3H, H2 and H3), 0.91 (m, 6H, H4).

$^{13}$C NMR (125.7 MHz, $CD_3CN$) $\delta$ = 172.9 (C1), 148.1 (C3′), 146.0 (C4′), 132.5 (C1′), 121.0 (C6′), 115.4 (C5′), 112.0 (C2′), 56.6 ($OCH_3$), 46.2 (C2), 43.2 (1′–C**H₂**), 27.0 (C3), 22.7 (C4).

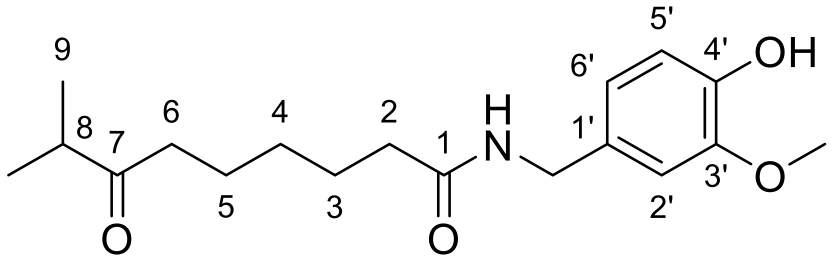


**31**, Capsicum-P10-16

$^{1}$H NMR (500.0 MHz, $CD_3CN$) $\delta$ = 6.85 (d, 1H, $J_{2',6'}$ = 2.0, H2′), 6.75 (d, 1H, $J_{5',6'}$ = 8.1, H5′), 6.70 (ddt, 1H, $J_{6',5'}$ = 8.1, $J_{6',2'}$ = 1.9, $J_{6',CH2}$ = 0.7, H6′), 6.65 (bs, 1H, NH), 6.42 (bs, 1H, OH), 4.22 (d, 2H, $J_{CH2,NH}$ = 6.0, 1′–C**H₂**), 3.82 (s, 3H, $OCH_3$), 2.59 (septet, 1H, $J_{8,9}$ = 6.9, H8), 2.45 (t, 2H, $J_{6,5}$ = 7.3, H6), 2.13 (m, 2H, H2), 1.57 (m, 2H, H3), 1.50 (m, 2H, H5), 1.26 (m, 2H, H4), 1.02 (d, 6H, $J_{9,8}$ = 6.9, H9).

$^{13}$C NMR (125.7 MHz, $CD_3CN$) $\delta$ = 215.4 (C7), 173.5 (C1), 148.1 (C3′), 146.1 (C4′), 132.4 (C1′), 121.0 (C6′), 115.4 (C5′), 112.0 (C2′), 56.6 ($OCH_3$), 43.2 (1′–C**H₂**), 41.3 (C8), 40.7 (C6), 36.7 (C2), 29.5 (C4), 26.4 (C3), 24.2 (C5), 18.5 (C9).

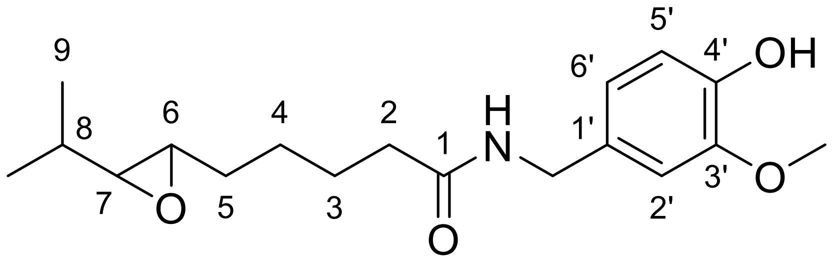


**32**, Capsicum-P10-17

$^{1}$H NMR (500.0 MHz, $CD_3CN$) $\delta$ = 6.86 (d, 1H, $J_{2',6'}$ = 2.0, H2′), 6.75 (d, 1H, $J_{5',6'}$ = 8.1, H5′), 6.71 (ddt, 1H, $J_{6',5'}$ = 8.0, $J_{6',2'}$ = 2.0, $J_{6',CH2}$ = 0.7, H6′), 6.66 (bs, 1H, NH), 6.42 (bs, 1H, OH), 4.23 (d, 2H, $J_{CH2,NH}$ = 6.0, 1′–C**H₂**), 3.82 (s, 3H, $OCH_3$), 2.66 (m, 1H, H6), 2.39 (m, 1H, H7), 2.15 (m, 2H, H2), 1.37–1.65 (m, 7H, H3, H4, H5 and H8), 0.95 (d, 3H, $J_{9,8}$ = 6.7, H9), 0.91 (d, 3H, $J_{9,8}$ = 6.9, H9).

$^{13}$C NMR (125.7 MHz, $CD_3CN$) $\delta$ = 173.4 (C1), 148.1 (C3′), 146.0 (C4′), 132.4 (C1′), 121.0 (C6′), 115.4 (C5′), 112.1 (C2′), 64.6 (C7), 58.1 (C6), 56.6 ($OCH_3$), 43.3 (1′–C**H₂**), 36.7 (C2), 32.6 (C5), 31.4 (C8), 26.4 (C3 or C4), 26.3 (C3 or C4), 19.3 and 18.6 (C9).

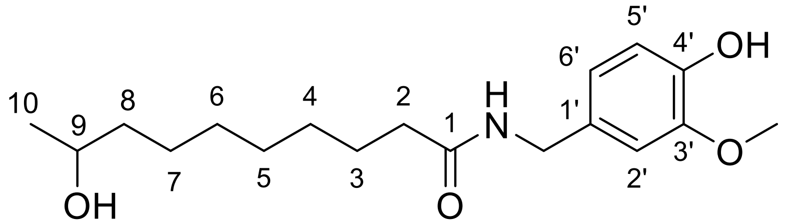


**33**, Capsicum11-P9

$^{1}$H NMR (500.0 MHz, $CD_3CN$) $\delta$ = 6.85 (d, 1H, $J_{2',6'}$ = 1.9, H2′), 6.75 (d, 1H, $J_{5',6'}$ = 8.0, H5′), 6.70 (ddt, 1H, $J_{6',5'}$ = 8.0, $J_{6',2'}$ = 2.0, H6′), 6.67 (bs, 1H, NH), 6.45 (bs, 1H, 4′–OH), 4.22 (d, 2H, $J_{CH2,NH}$ = 6.0, 1′–C**H₂**), 3.82 (s, 3H, $OCH_3$), 3.64 (m, 1H, H9), 2.48 (m, 1H, 9–O**H**), 2.13 (m, 2H, H2), 1.56 (m, 2H, H3), 1.35 (m, 2H, H8), 1.24–1.31 (m, 8H, H4, H5, H6 and H7), 1.07 (d, 3H, $J_{10,9}$ = 6.1, H10).

$^{13}C$ NMR (125.7 MHz, $CD_3CN$) $\delta$ = 173.6 (C1), 148.1 (C3′), 146.0 (C4′), 132.4 (C1′), 121.0 (C6′), 115.4 (C5′), 112.0 (C2′), 67.8 (C9), 56.6 ($OCH_3$), 43.2 (1′–C**H₂**), 40.1 (C8), 36.9 (C2), 30.3, 30.1 and 29.9 (C4, C5 and C6), 26.6 (C3 or C7), 26.5 (C3 or C7), 23.9 (C10).

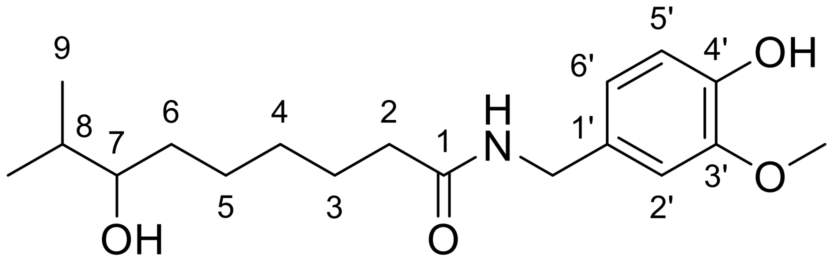


**34**, Capsicum11-P11

$^1H$ NMR (500.0 MHz, $CD_3CN$) $\delta$ = 6.85 (d, 1H, $J_{2',6'}$ = 1.9, H2′), 6.75 (d, 1H, $J_{5',6'}$ = 8.1, H5′), 6.71 (ddt, 1H, $J_{6',5'}$ = 8.0, $J_{6',2'}$ = 1.9, $J_{6',CH2}$ = 0.6, H6′), 6.67 (bs, 1H, NH), 6.44 (bs, 1H, 4′–OH), 4.22 (d, 2H, $J_{CH2,NH}$ = 6.0, 1′–C**H₂**), 3.82 (s, 3H, $OCH_3$), 3.22 (m, 1H, H7), 2.37 (bd, 1H, $J_{OH,7}$ = 5.5, 7–O**H**), 2.14 (m, 2H, H2), 1.52–1.61 (m, 3H, H3 and H8), 1.24–1.46 (m, 6H, H4, H5 and H6), 0.86 and 0.85 (2 × d, 2 × 3H, $J_{9,8}$ = 6.6, H9).

$^{13}C$ NMR (125.7 MHz, $CD_3CN$) $\delta$ = 173.6 (C1), 148.1 (C3′), 146.0 (C4′), 132.4 (C1′), 121.0 (C6′), 115.4 (C5′), 112.0 (C2′), 76.5 (C7), 56.6 ($OCH_3$), 43.2 (1′–C**H₂**), 36.9 (C2), 34.8 (C6), 34.4 (C8), 30.0 (C4), 26.6 (C3 or C5), 26.6 (C3 or C5), 19.3 and 17.6 (C9).

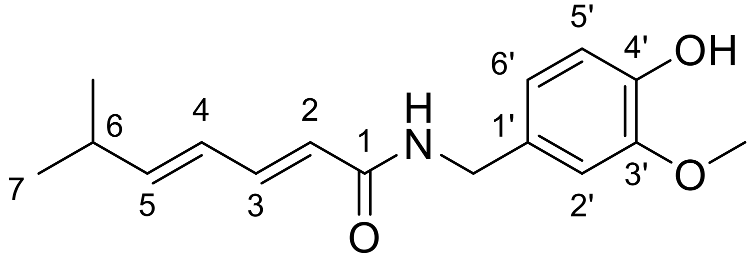


**35**, ISO-276

$^1H$ NMR (500.0 MHz, DMSO-*d*$_6$) $\delta$ = 8.33 (t, 1H, $J_{NH,CH2}$ = 5.9, NH), 7.01 (dd, 1H, $J_{3,2}$ = 15.1, $J_{3,4}$ = 10.6, H3), 6.82 (d, 1H, $J_{2',6'}$ = 2.0, H2′), 6.70 (d, 1H, $J_{5',6'}$ = 8.0, H5′), 6.64 (dd, 1H, $J_{6',5'}$ = 8.0, $J_{6',2'}$ = 2.0, H6′), 6.14 (dd, 1H, $J_{4,5}$ = 15.4, $J_{4,3}$ = 10.5, H4), 6.07 (dd, 1H, $J_{5,4}$ = 15.3, $J_{5,6}$ = 6.4, H5), 5.98 (d, 1H, $J_{2,3}$ = 15.1, H2), 4.22 (d, 2H, $J_{CH2,NH}$ = 5.9, 1′–C**H₂**), 3.73 (s, 3H, $OCH_3$), 2.40 (m, 1H, H6), 1.00 (d, 6H, $J_{7,6}$ = 6.8, H7).

Small amount of the sample did not allow for $^{13}C$ NMR measurement.

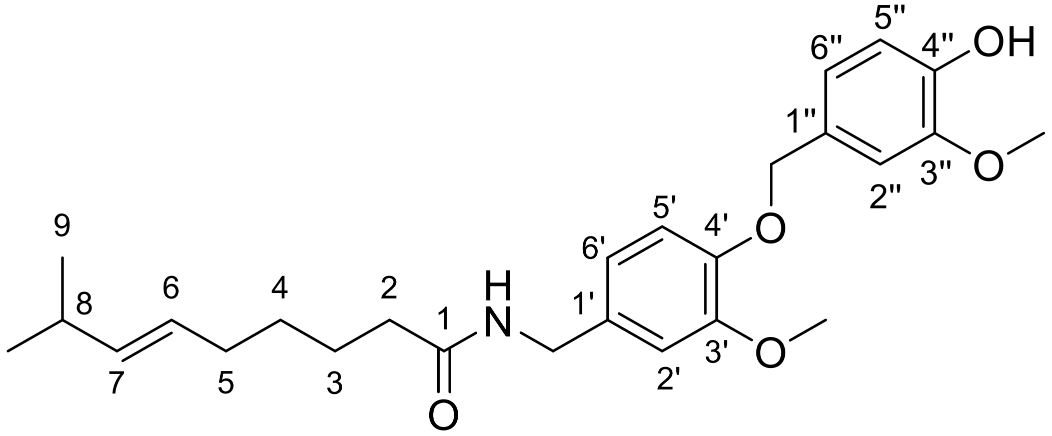

**36**, JS-ISO-21-20-442

$^{1}$H NMR (600.1 MHz, $CD_3CN$) $\delta$ = 7.03 (d, 1H, $J_{2'',6''}$ = 1.9, H2″), 6.92 (d, 1H, $J_{5',6'}$ = 8.1, H5′), 6.86–6.88 (m, 2H, H2′ and H6″), 6.80 (d, 1H, $J_{5'',6''}$ = 8.1, H5″), 6.76 (dd, 1H, $J_{6',5'}$ = 8.1, $J_{6',2'}$ = 2.0, H6′), 6.69 (bs, 1H, NH), 5.33–5.42 (m, 2H, H6 and H7), 4.94 (s, 2H, 1″–C**H₂**), 4.24 (d, 2H, $J_{CH2,NH}$ = 6.0, 1′–C**H₂**), 3.85 (s, 3H, 3″–$OCH_3$), 3.78 (s, 3H, 3′–$OCH_3$), 2.22 (m, 1H, H8), 2.15 (m, 2H, H2), 1.97 (m, 2H, H5), 1.57 (m, 2H, H3), 1.34 (m, 2H, H4), 0.95 (d, 6H, $J_{9,8}$ = 6.8, H9).

$^{13}$C NMR (150.9 MHz, $CD_3CN$) $\delta$ = 173.6 (C1), 150.7 (C3′), 148.2 (C3″), 148.2 (C4′), 147.0 (C4″), 138.7 (C7), 133.8 (C1′), 130.0 (C1″), 127.8 (C6), 122.3 (C6″), 120.4 (C6′), 115.5 (C5″), 114.9 (C5′), 112.9 (C2″), 112.5 (C2′), 71.8 (1″–C**H₂**), 56.7 (3″–O**C**$H_3$), 56.3 (3′–O**C**$H_3$), 43.1 (1′–C**H₂**), 36.7 (C2), 32.9 (C5), 31.8 (C8), 30.0 (C4), 26.1 (C3), 23.0 (C9).

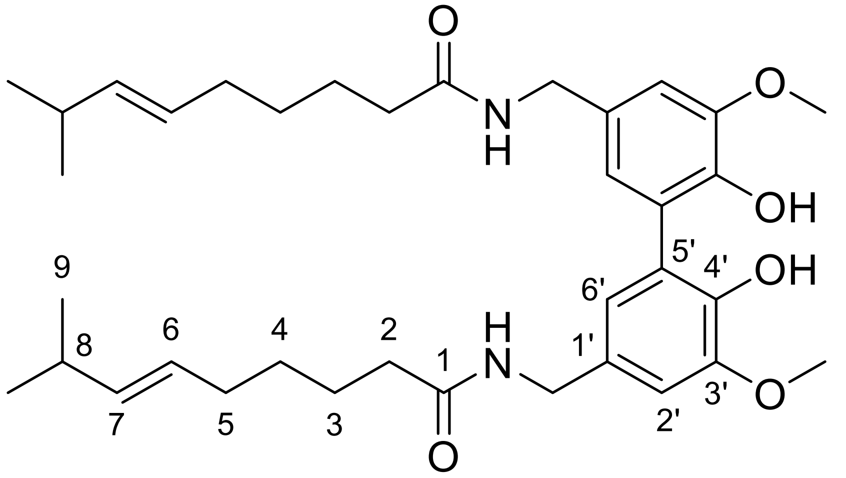

**38**, Capsicum11-24

$^{1}$H NMR (500.0 MHz, $CD_3CN$) $\delta$ = 6.85 (d, 2H, $J_{2',6'}$ = 2.0, H2′), 6.69 (d, 2H, $J_{6',2'}$ = 2.1, H6′), 6.68 (bs, 2H, NH), 5.31–5.41 (m, 4H, H6 and H7), 4.26 (d, 4H, $J_{CH2,NH}$ = 6.0, 1′–C**H₂**), 3.85 (s, 6H, $OCH_3$), 2.21 (m, 2H, H8), 2.13 (m, 4H, H2), 1.95 (m, 4H, H5), 1.57 (m, 4H, H3), 1.32 (m, 4H, H4), 0.93 (d, 12H, $J_{9,8}$ = 6.8, H9).

$^{13}$C NMR (125.7 MHz, $CD_3CN$) $\delta$ = 173.3 (C1), 148.6 (C3′), 144.2 (C4′), 138.5 (C7), 131.0 (C1′), 127.6 (C6), 126.0 (C5′), 122.7 (C6′), 110.8 (C2′), 56.7 ($OCH_3$), 43.5 (1′–C**H₂**), 36.7 (C2), 32.8 (C5), 31.8 (C8), 29.9 (C4), 26.0 (C3), 23.1 (C9).

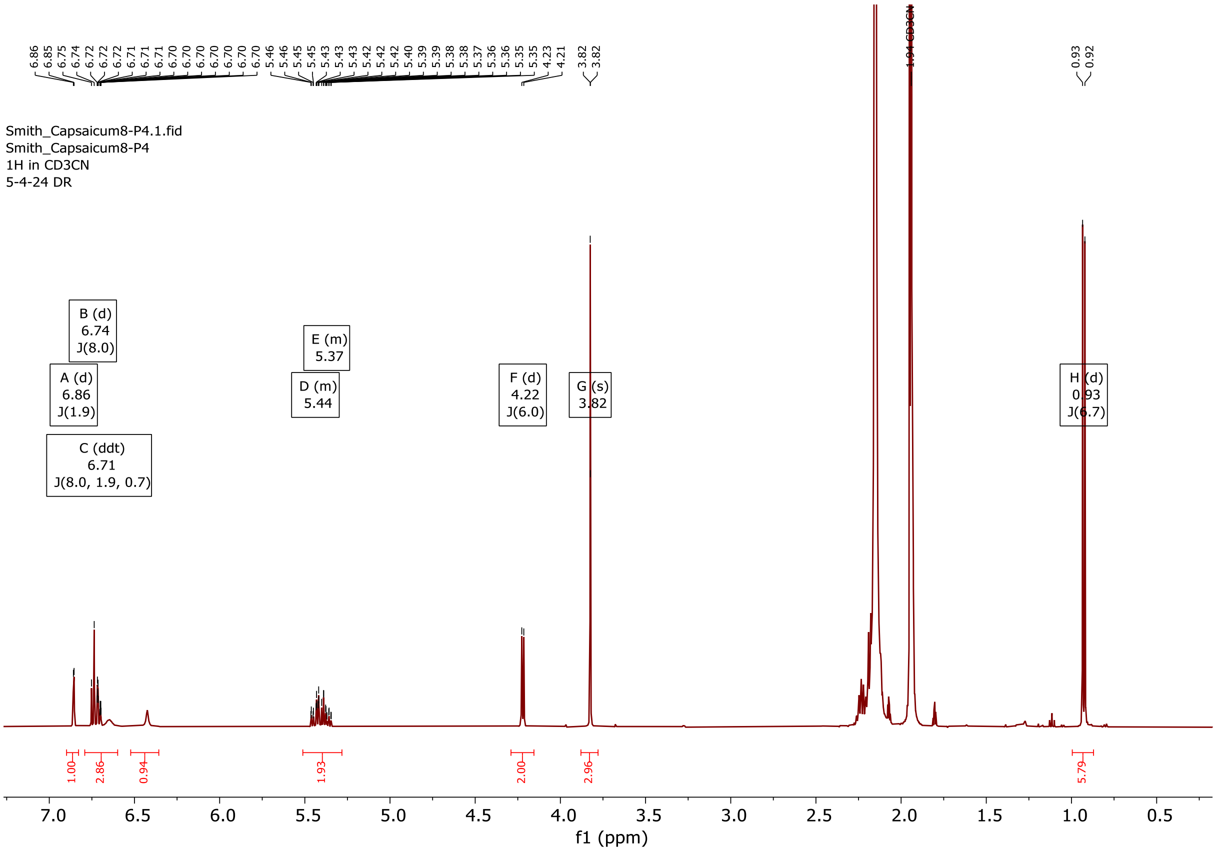


$^{1}$H NMR spectrum of compound **10**.

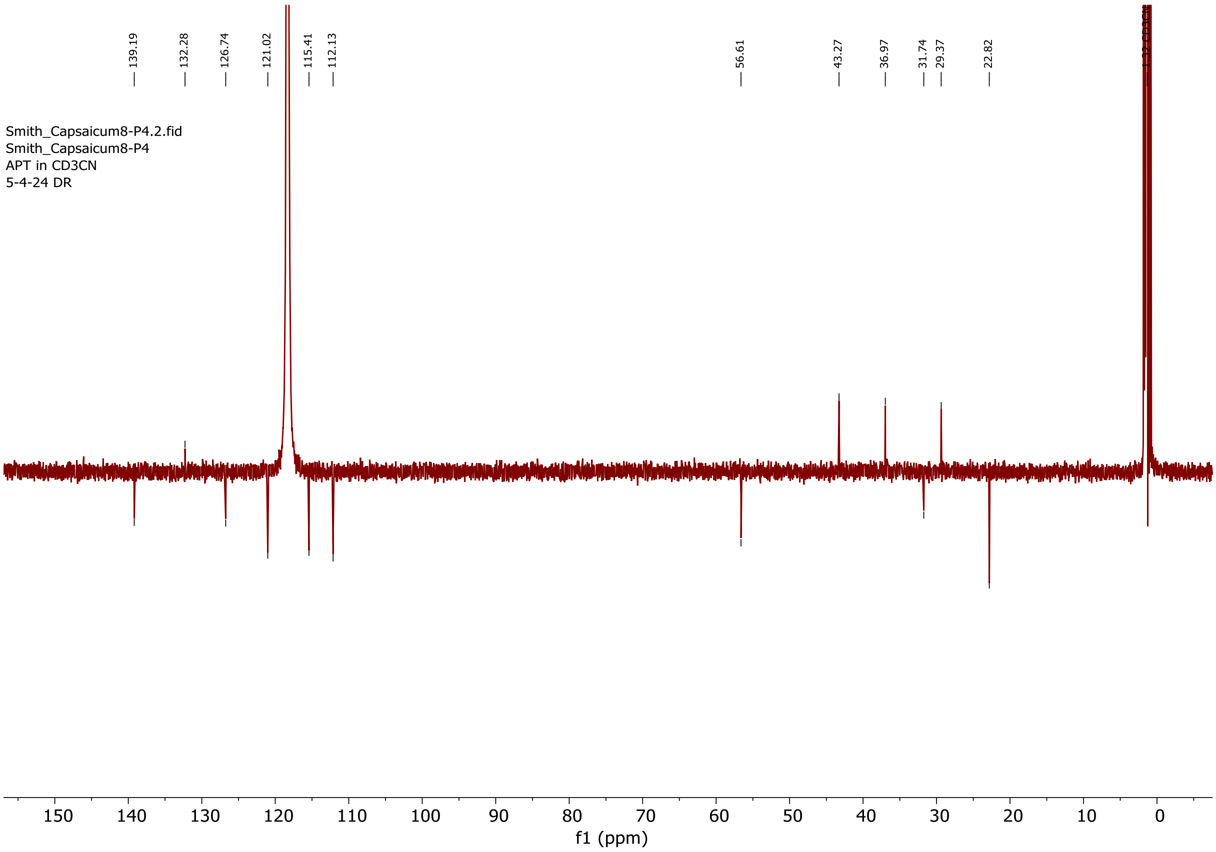


$^{13}$C NMR spectrum of compound **10**.

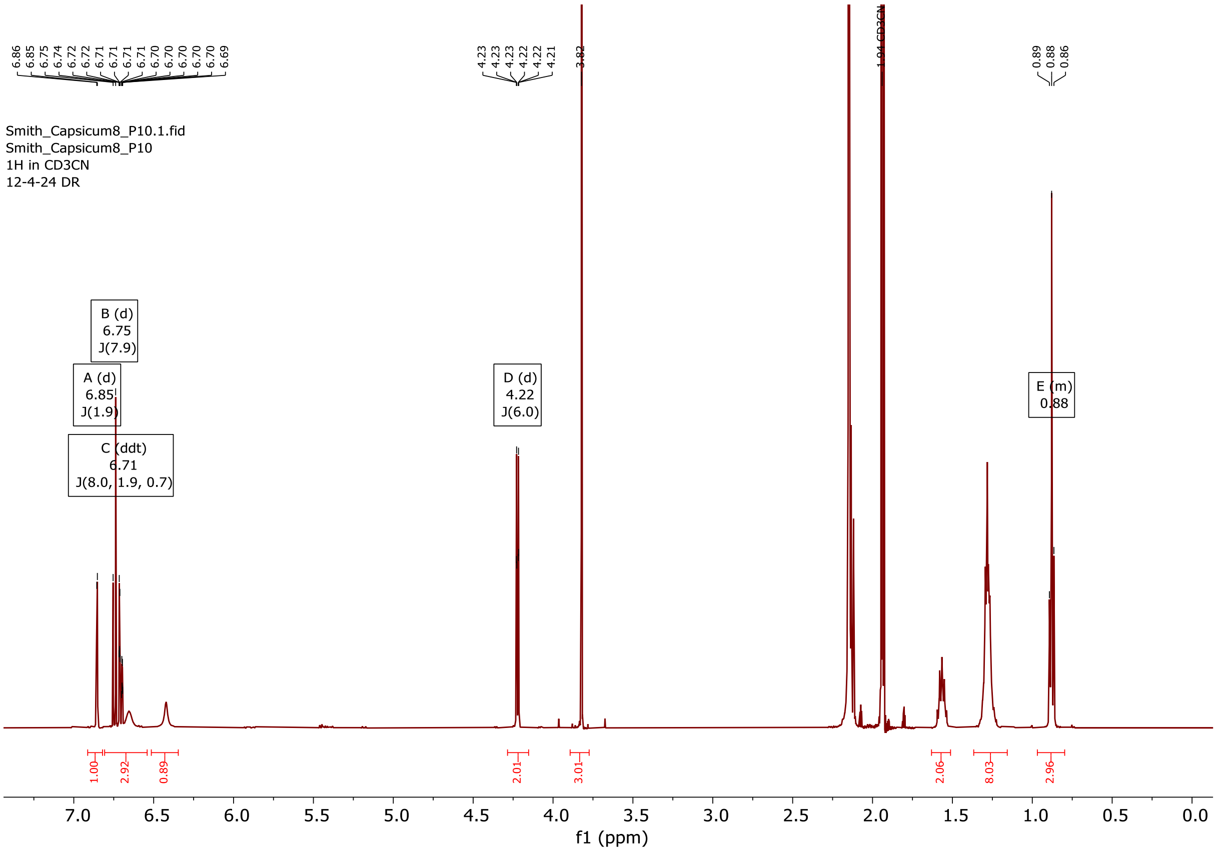


$^{1}H$ NMR spectrum of compound **13**.

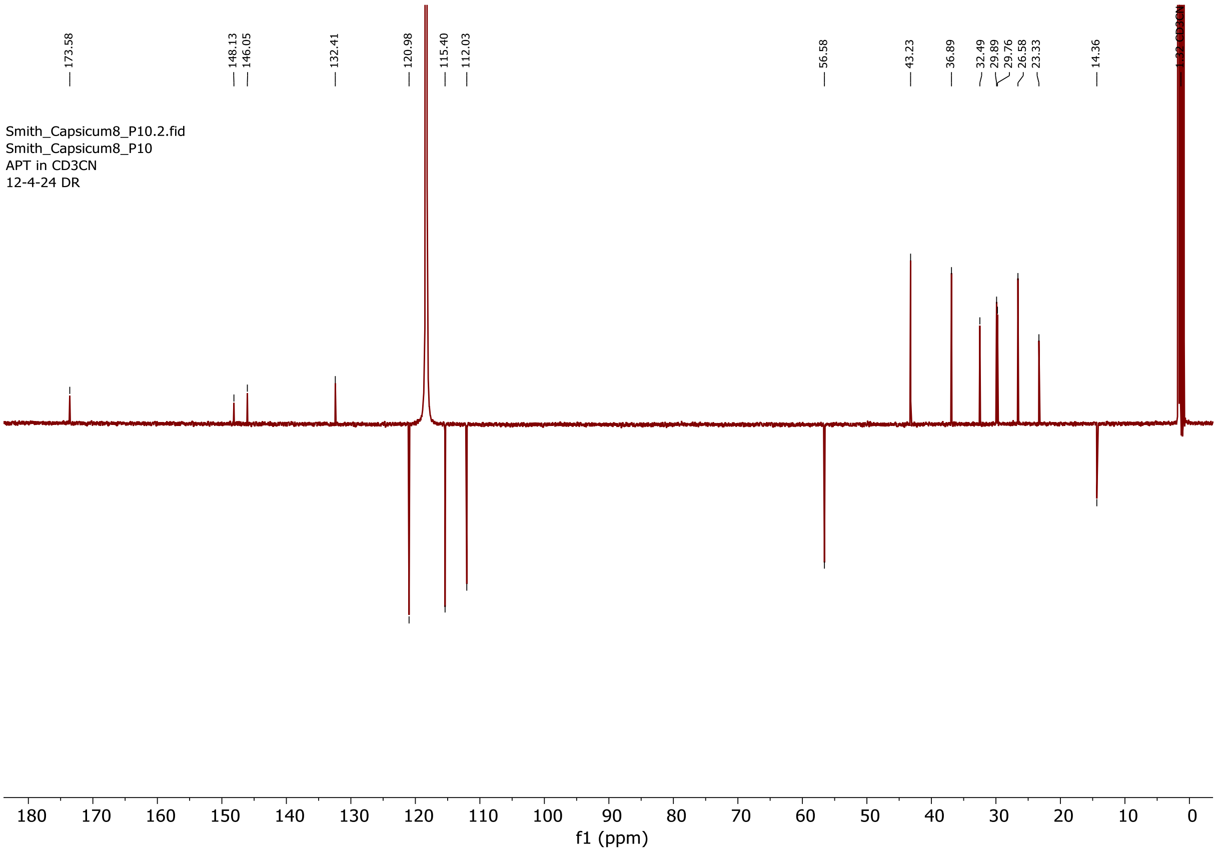


$^{13}C$ NMR spectrum of compound **13**.

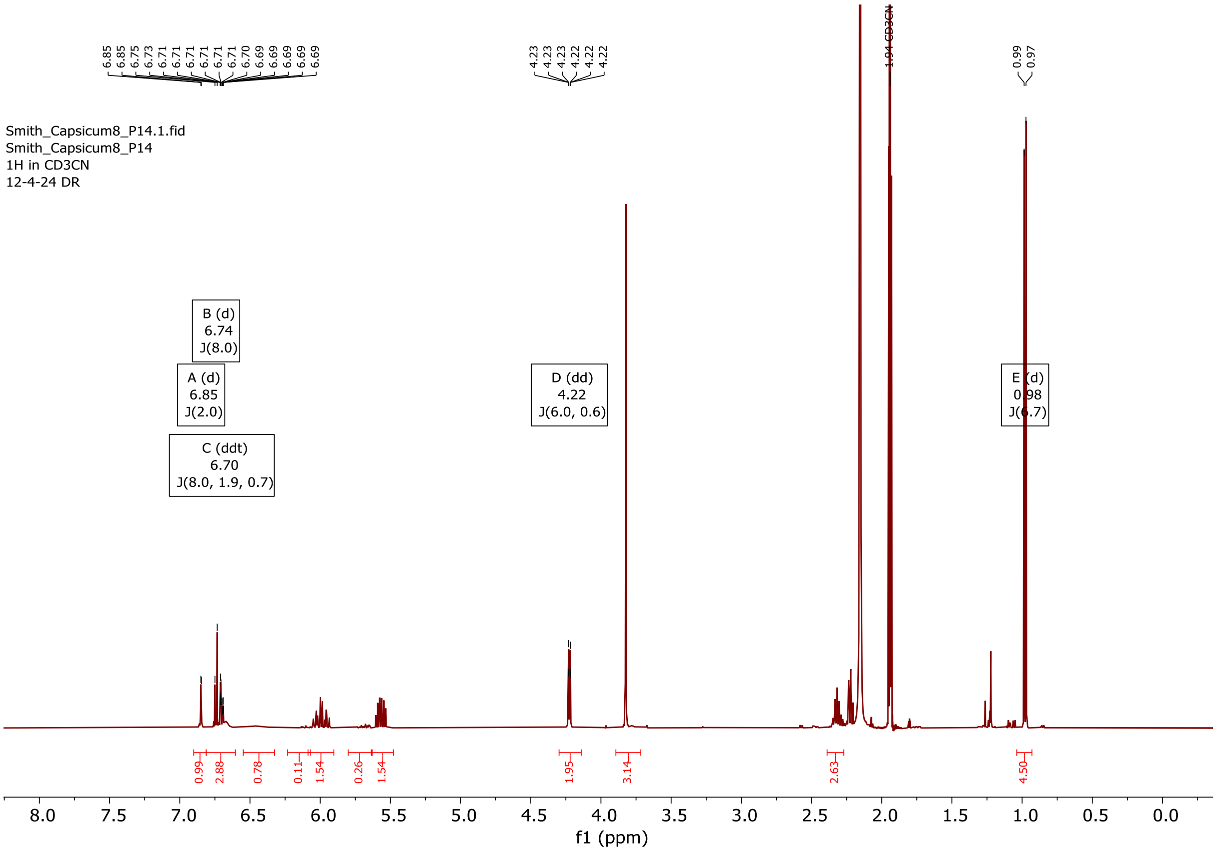


[1]H NMR spectrum of compound **16**.

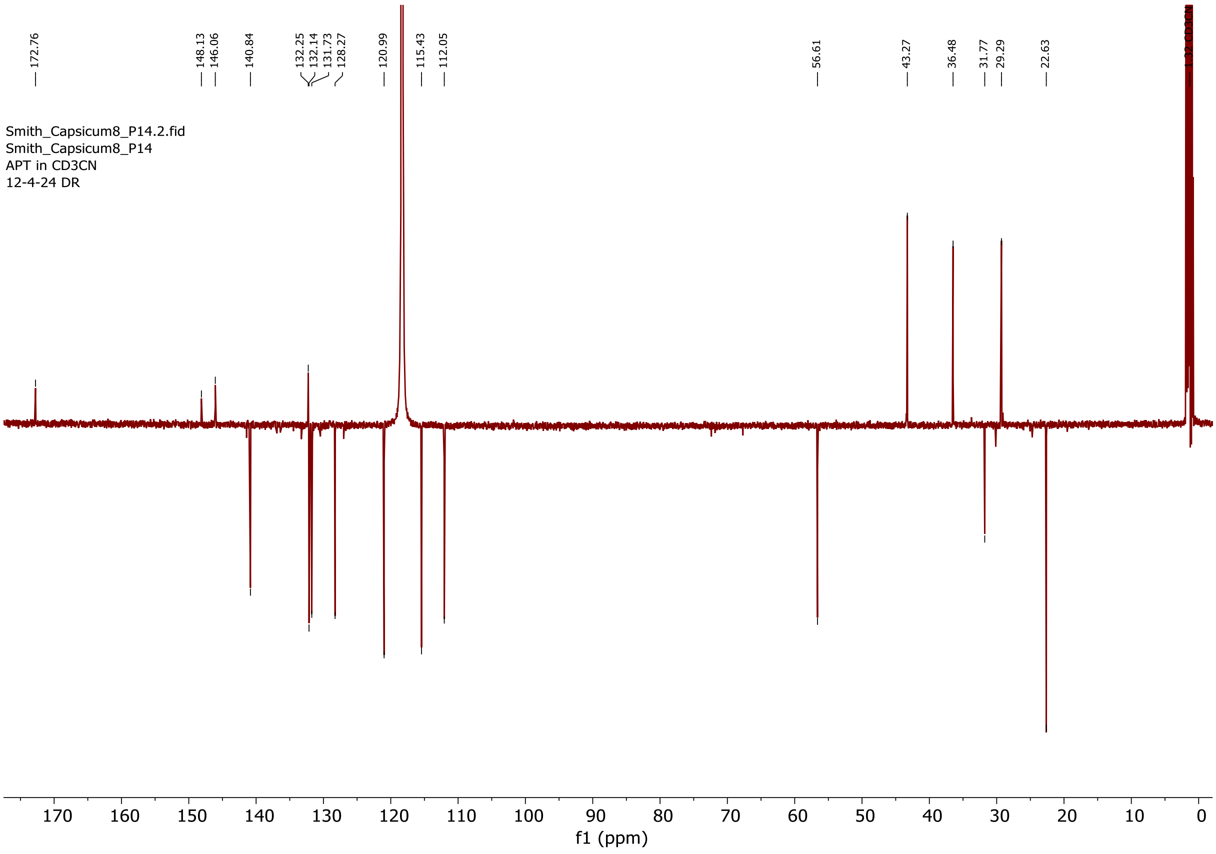


[13]C NMR spectrum of compound **16**.

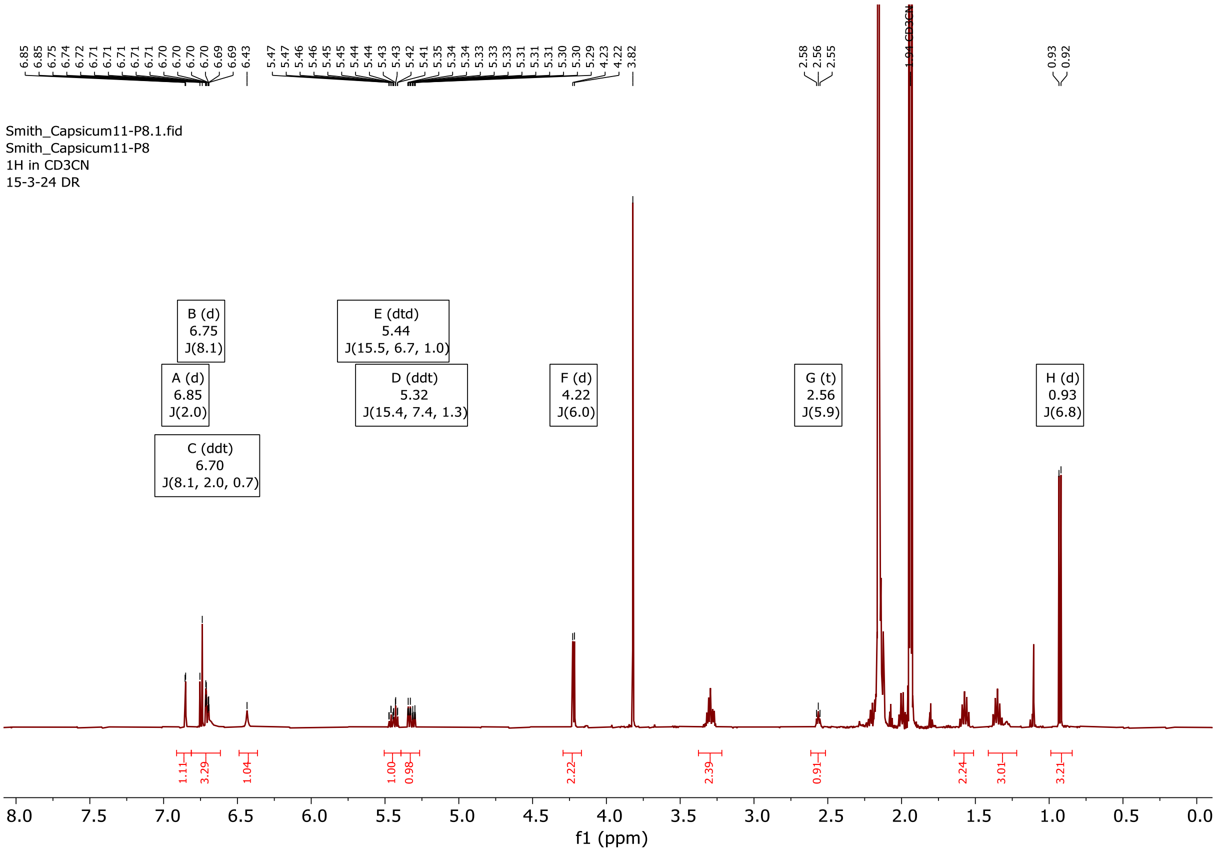


$^{1}H$ NMR spectrum of compound **22**.

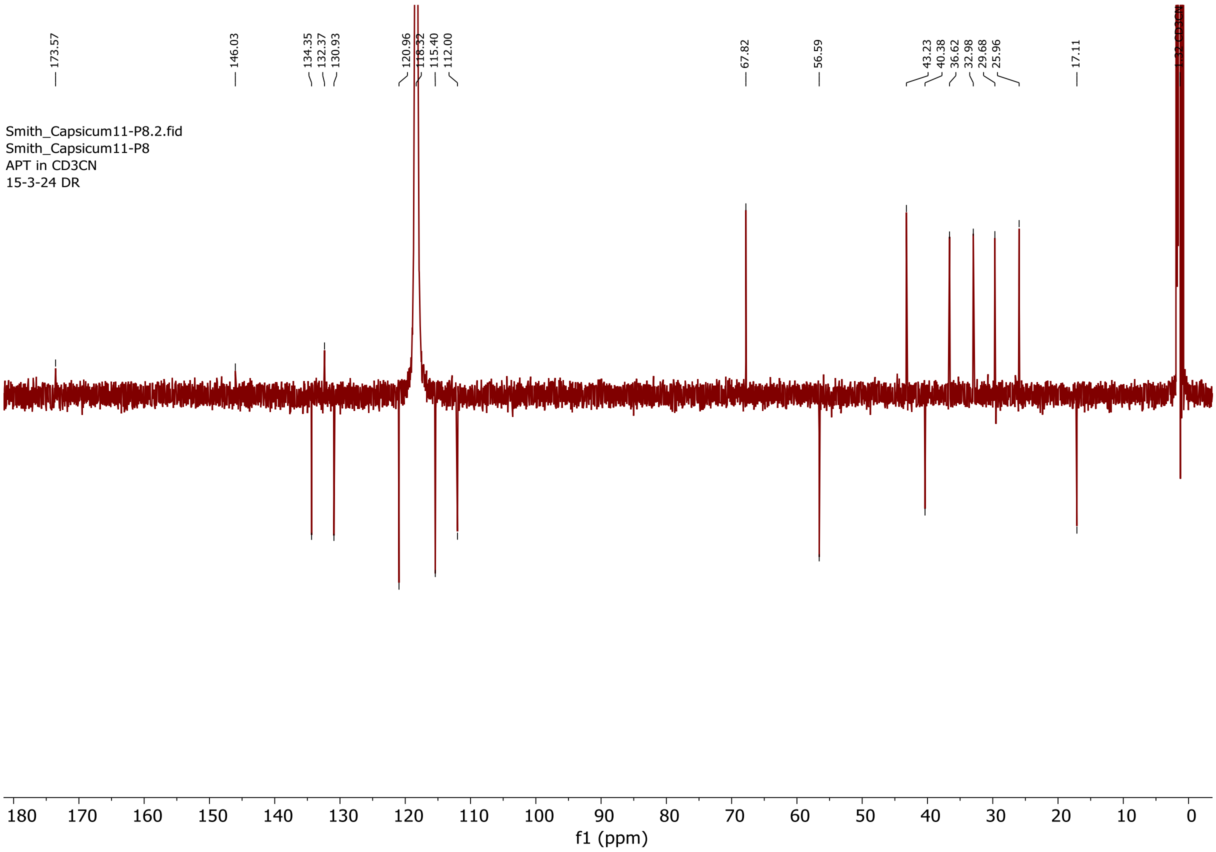


$^{13}C$ NMR spectrum of compound **22**.

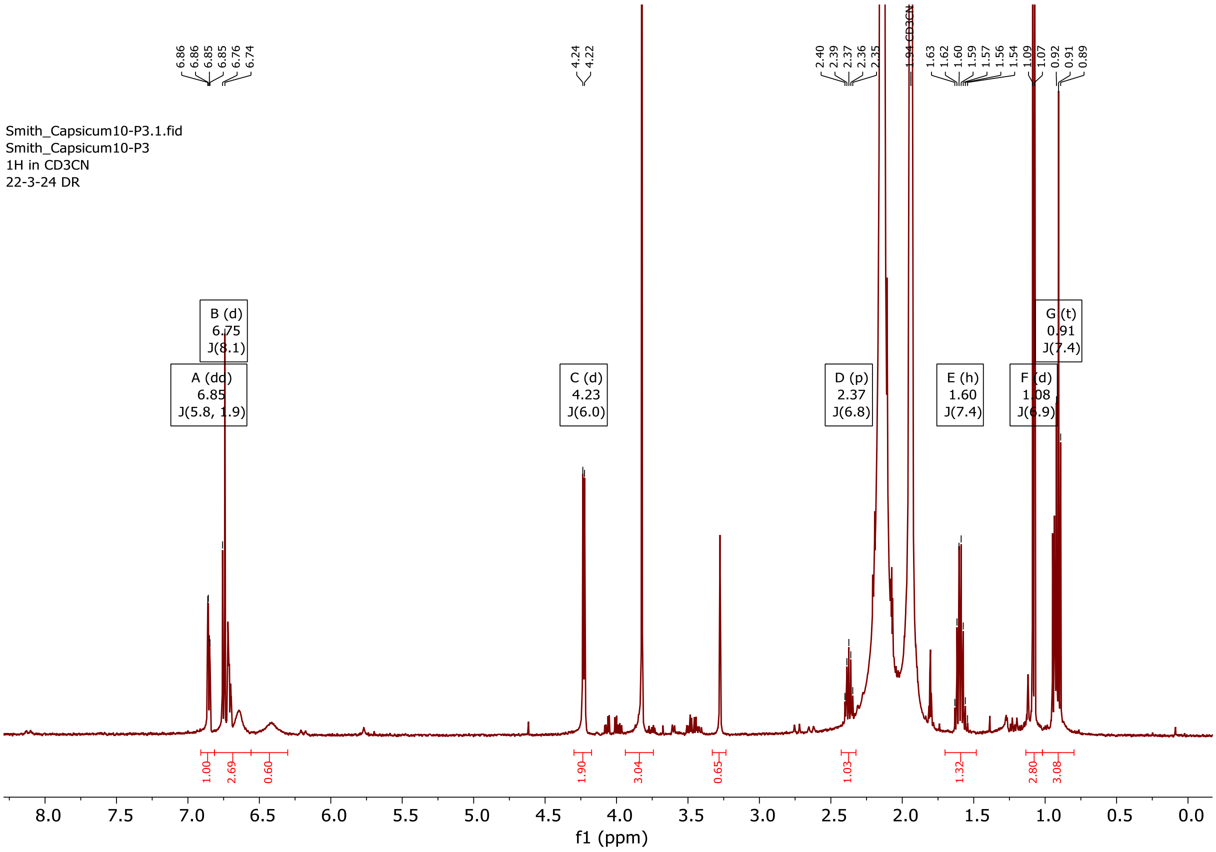


$^{1}H$ NMR spectrum of compound **26**.

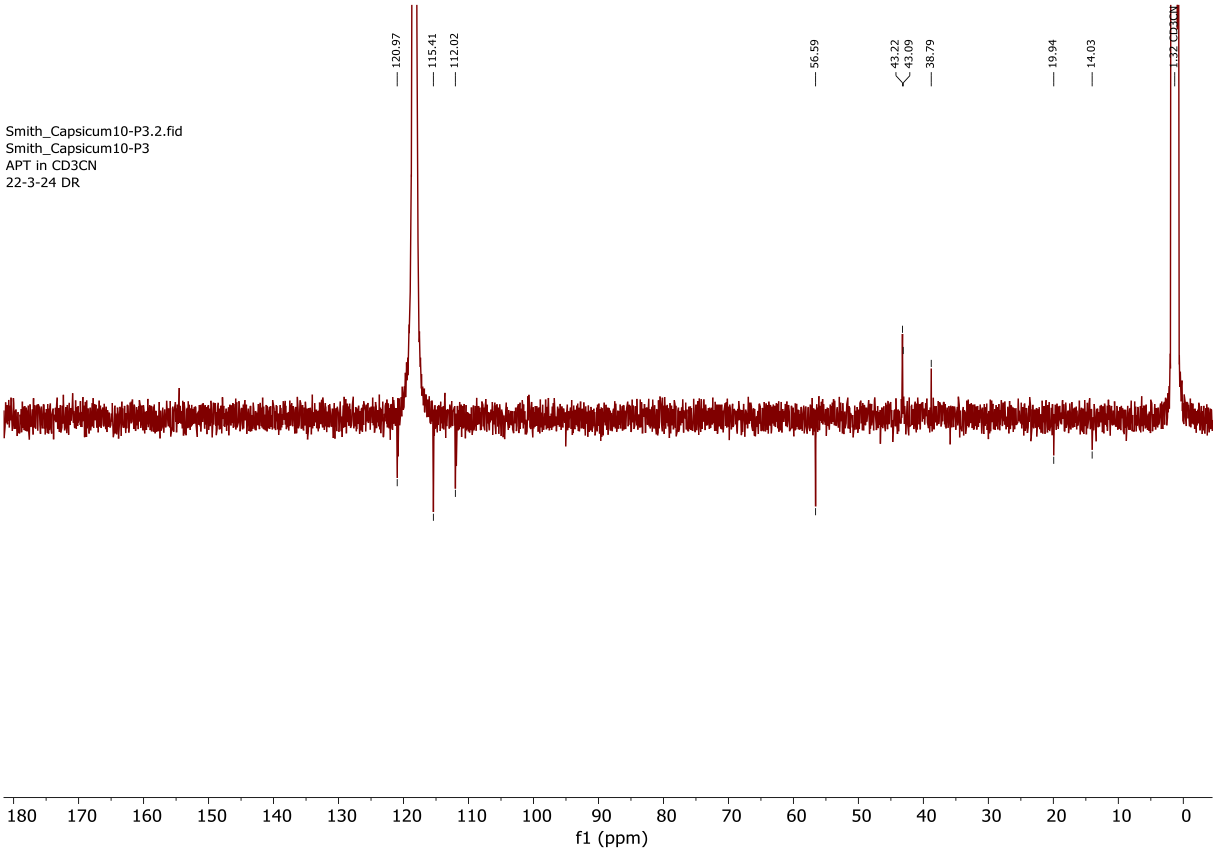


$^{13}C$ NMR spectrum of compound **26**.

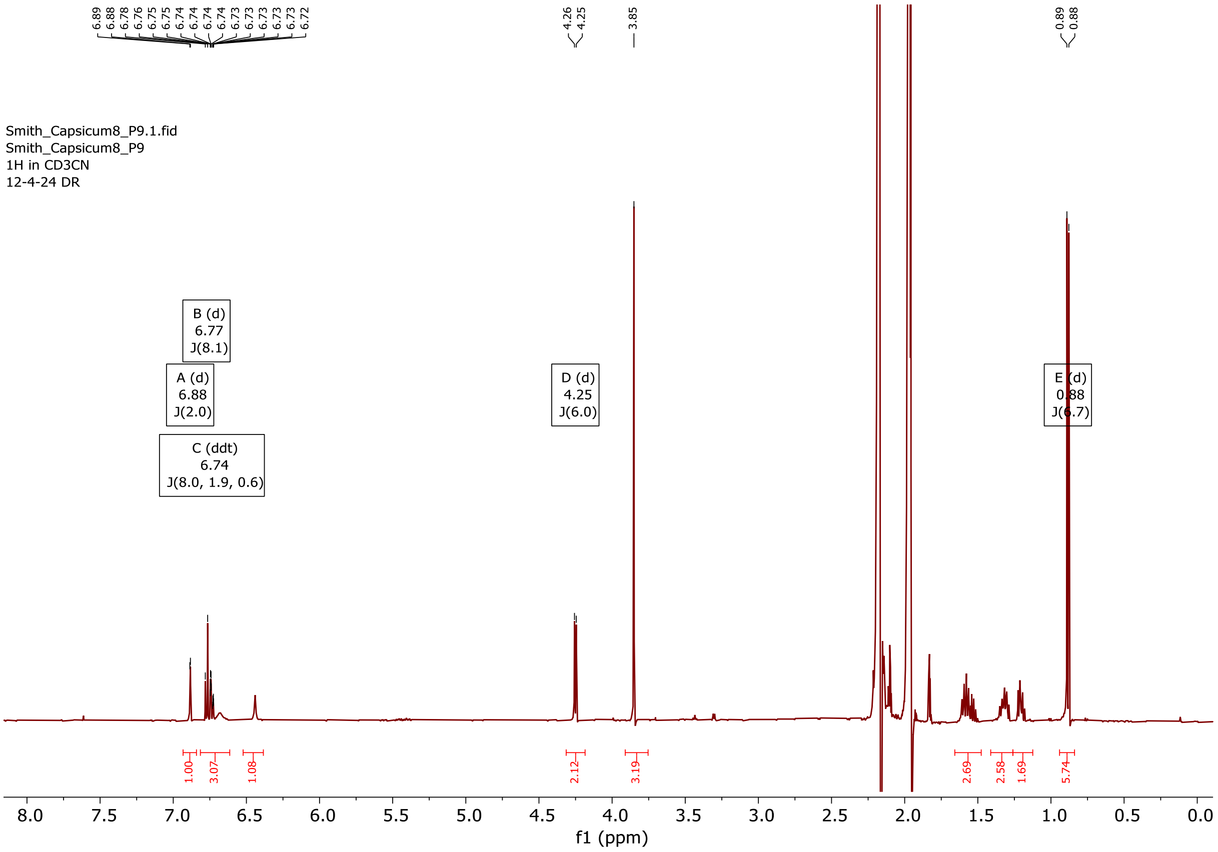


[1]H NMR spectrum of compound **27**.

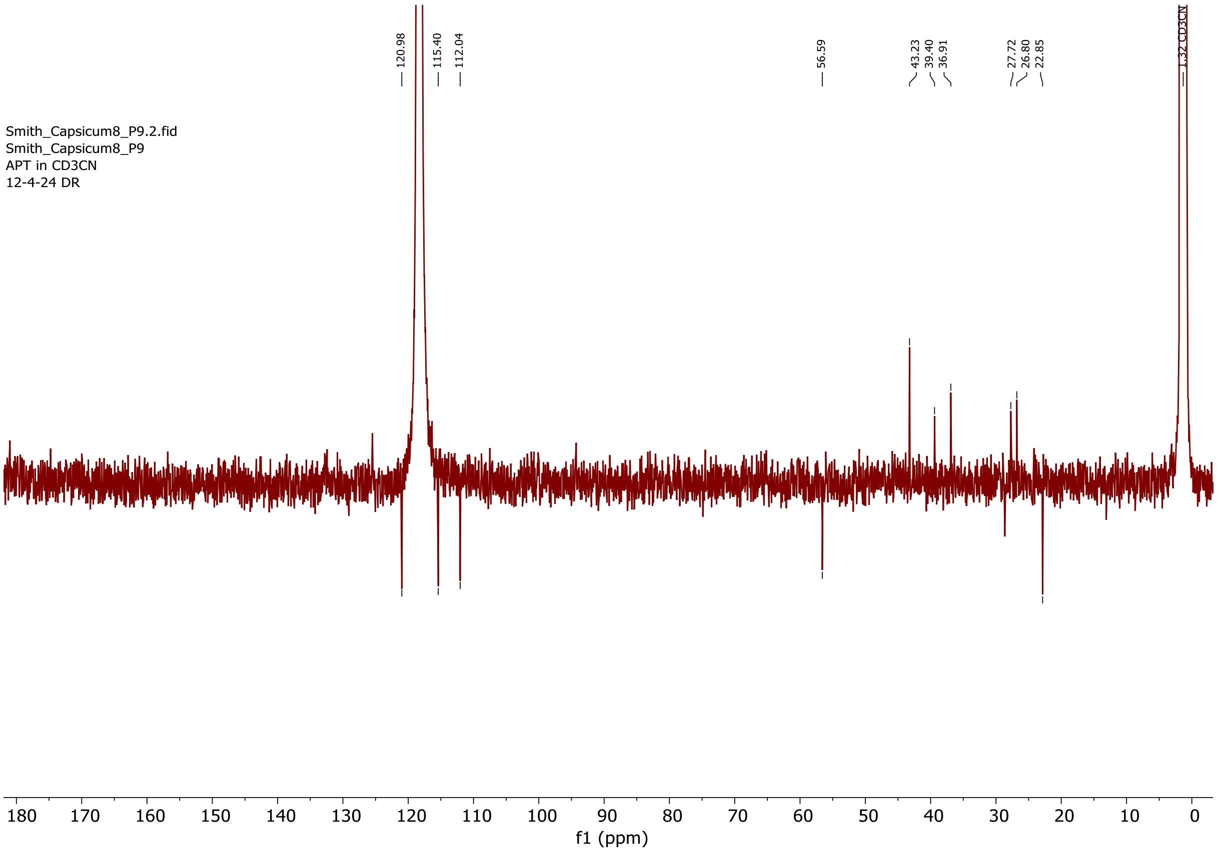


[13]C NMR spectrum of compound **27**.

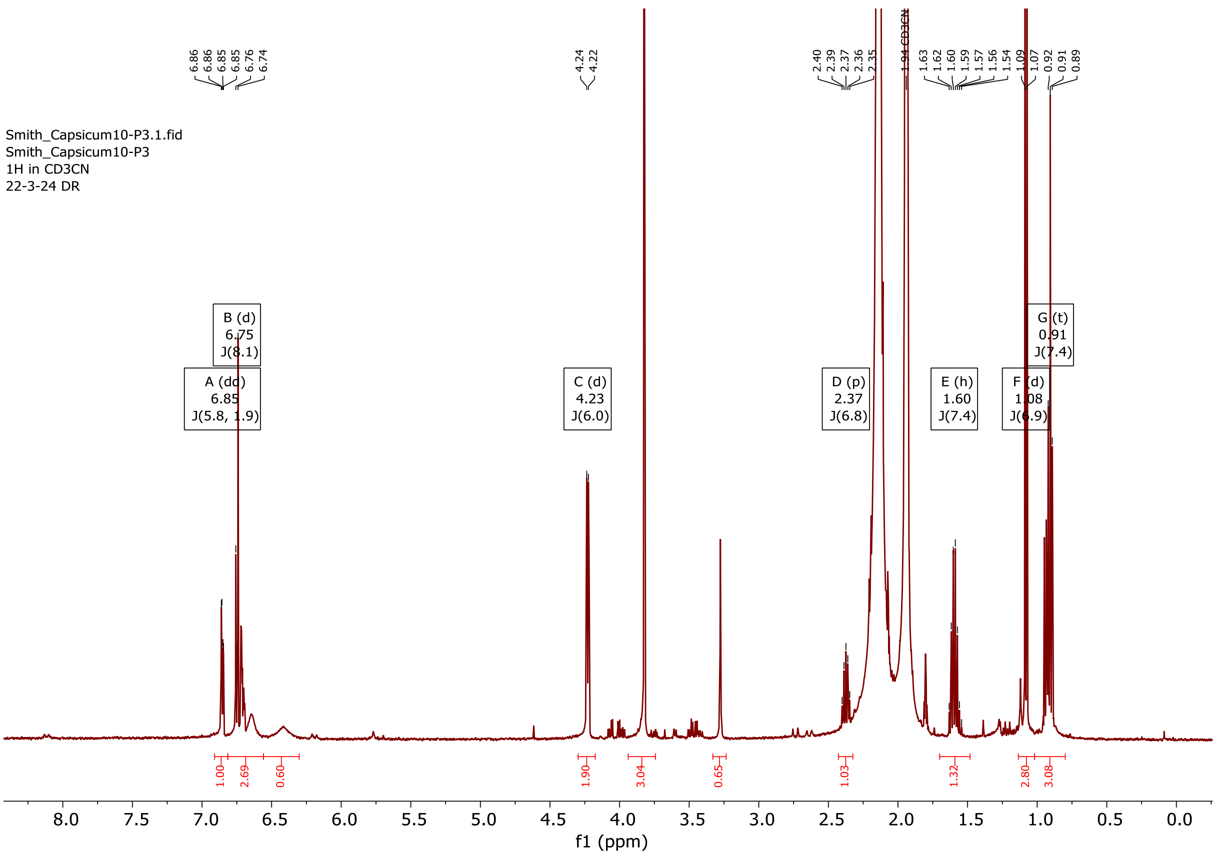


$^{1}H$ NMR spectrum of a mixture of compounds **28** and **29**.

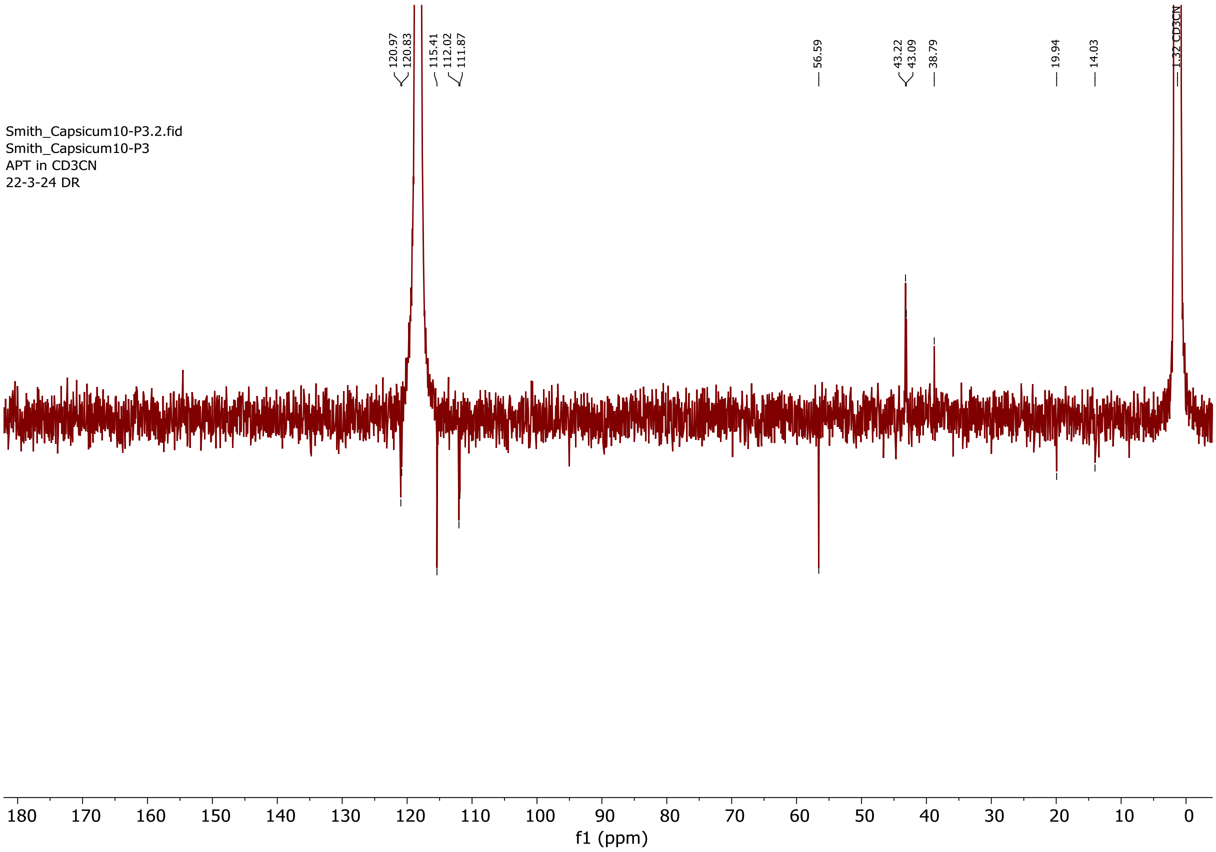


$^{13}C$ NMR spectrum of a mixture of compounds **28** and **29**.

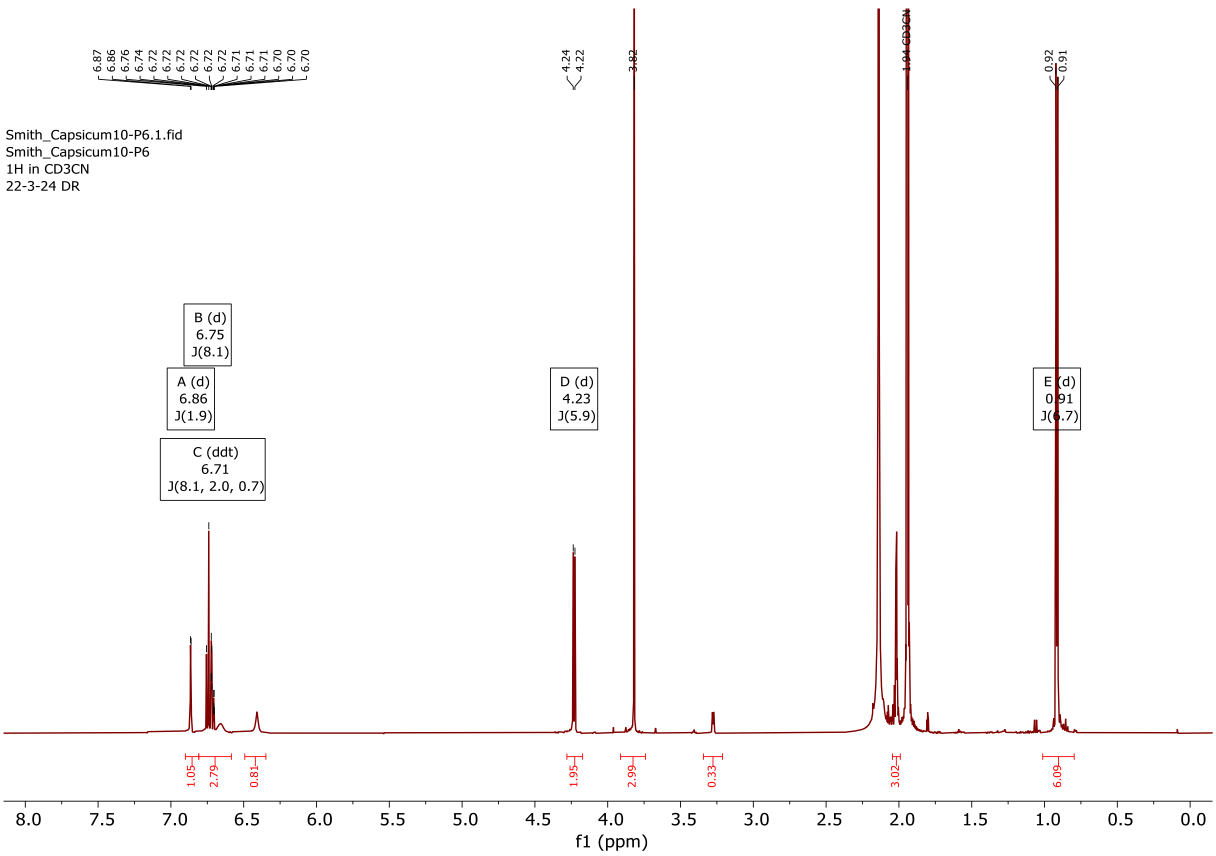


$^{1}H$ NMR spectrum of compound **30**.

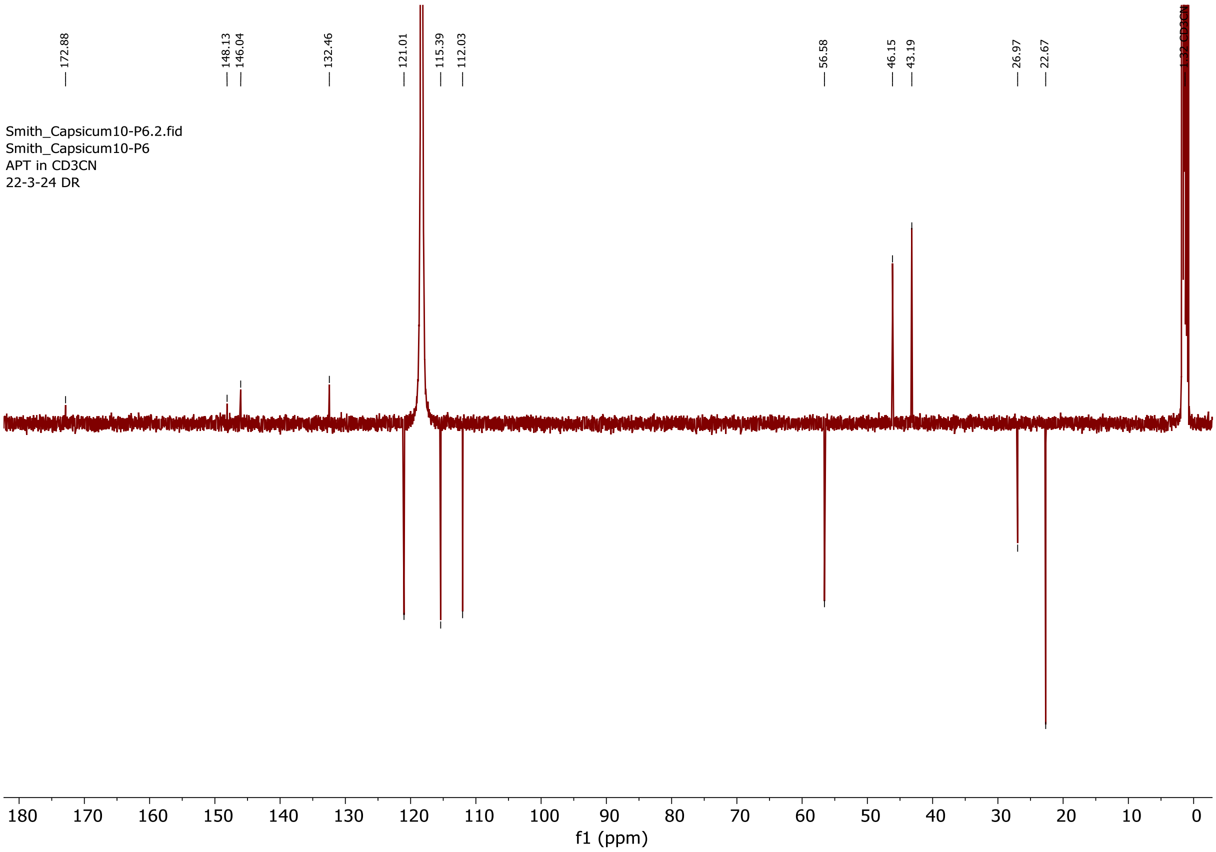


$^{13}C$ NMR spectrum of compound **30**.

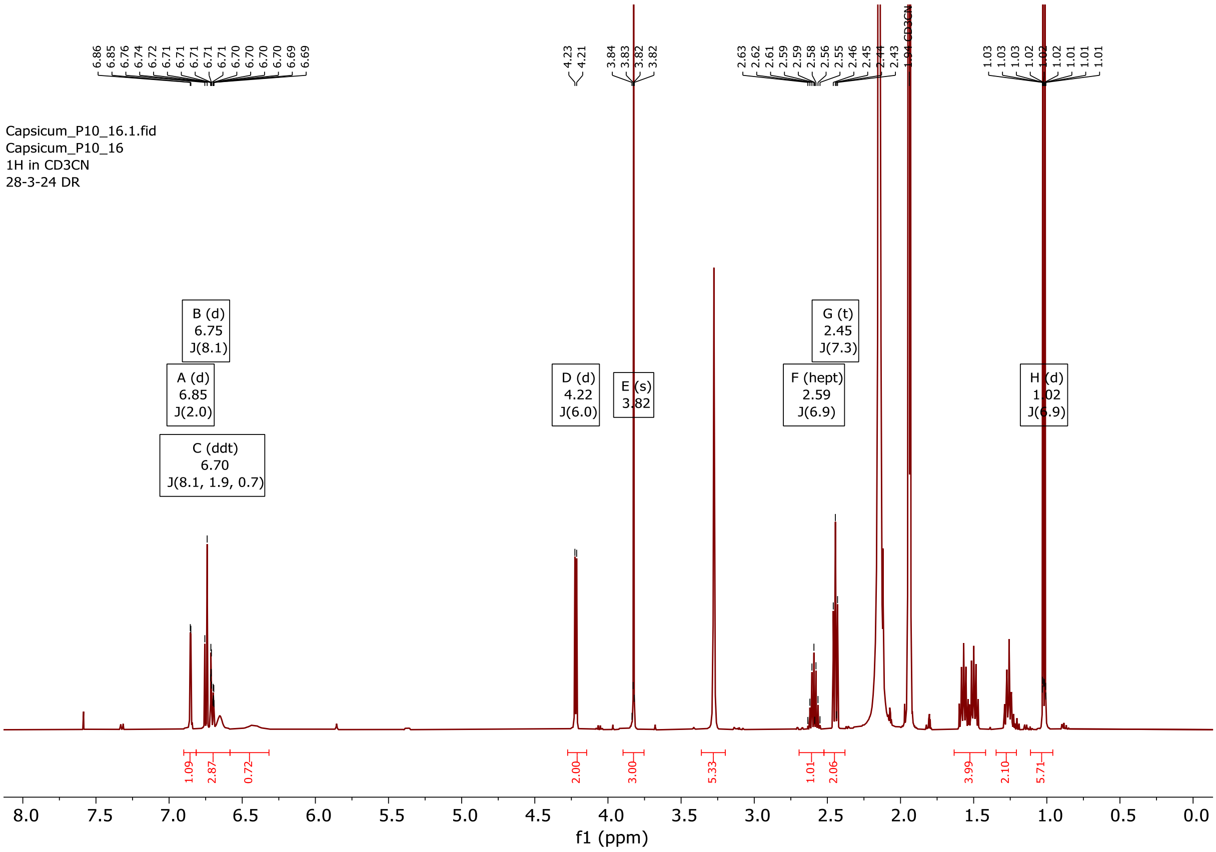


$^{1}$H NMR spectrum of compound **31**.

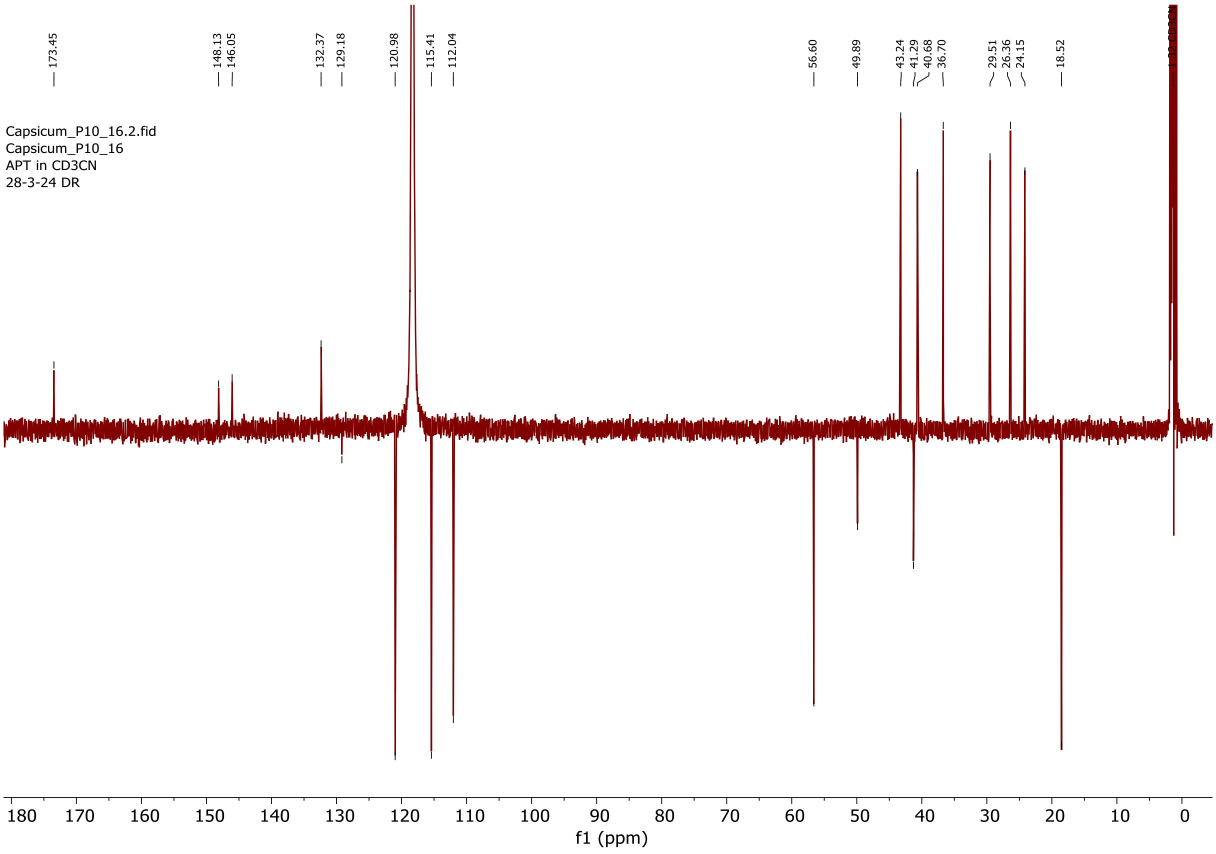


$^{13}$C NMR spectrum of compound **31**.

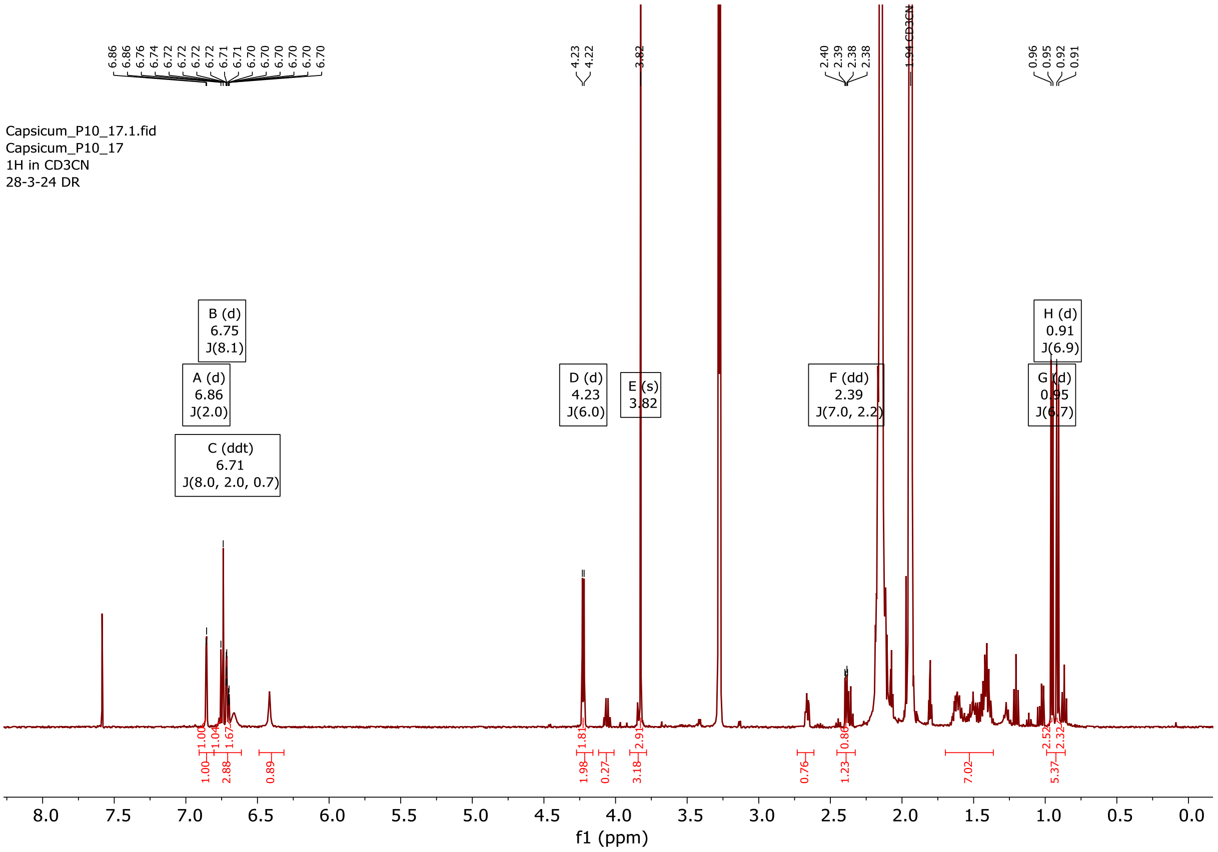


$^{1}H$ NMR spectrum of compound **32**.

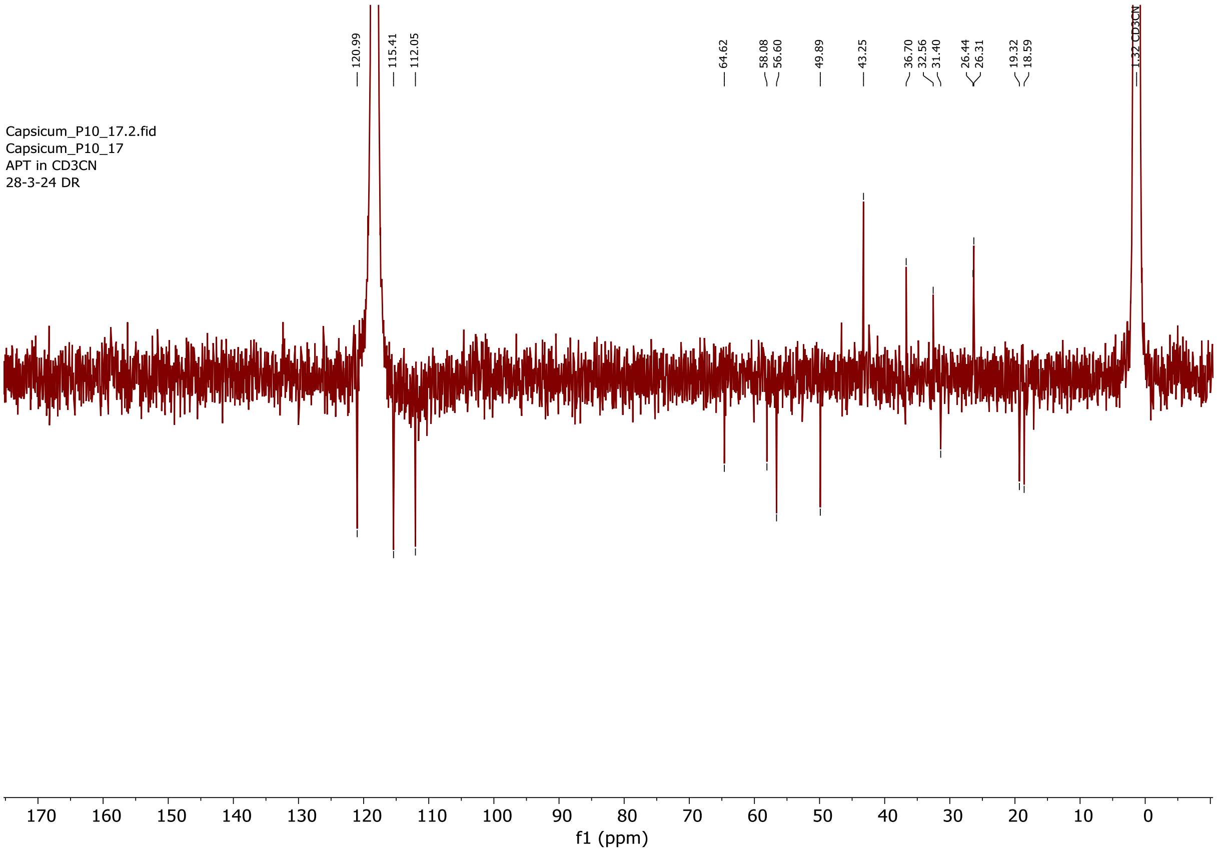


$^{13}C$ NMR spectrum of compound **32**.

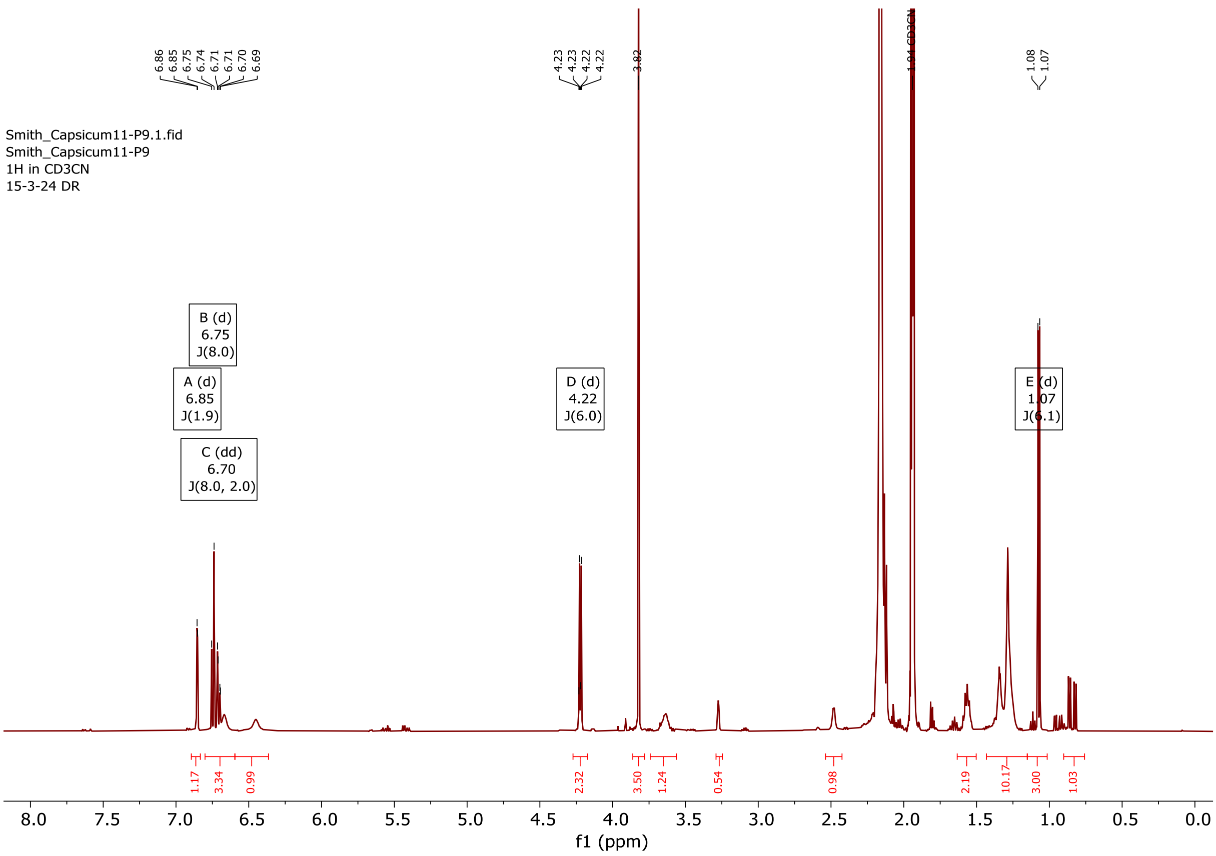


$^{1}$H NMR spectrum of compound **33**.

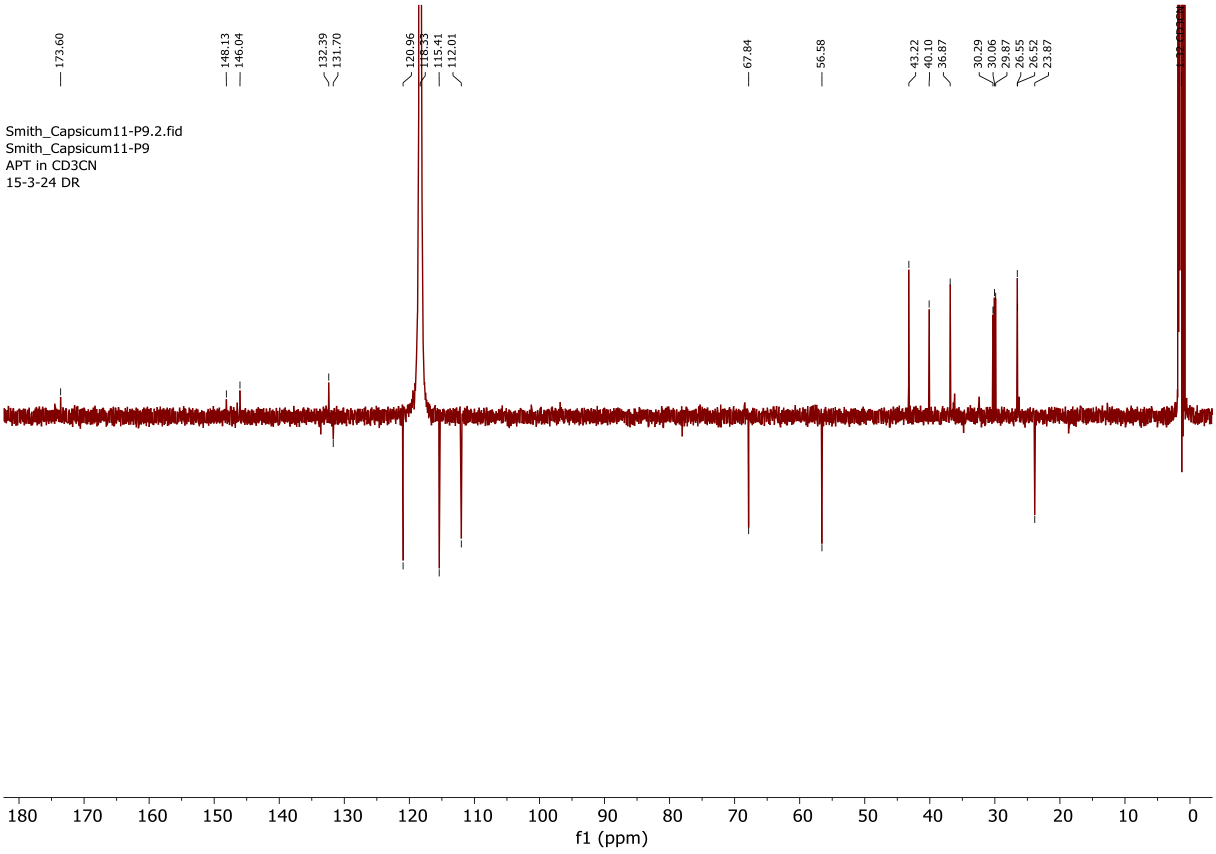


$^{13}$C NMR spectrum of compound **33**.

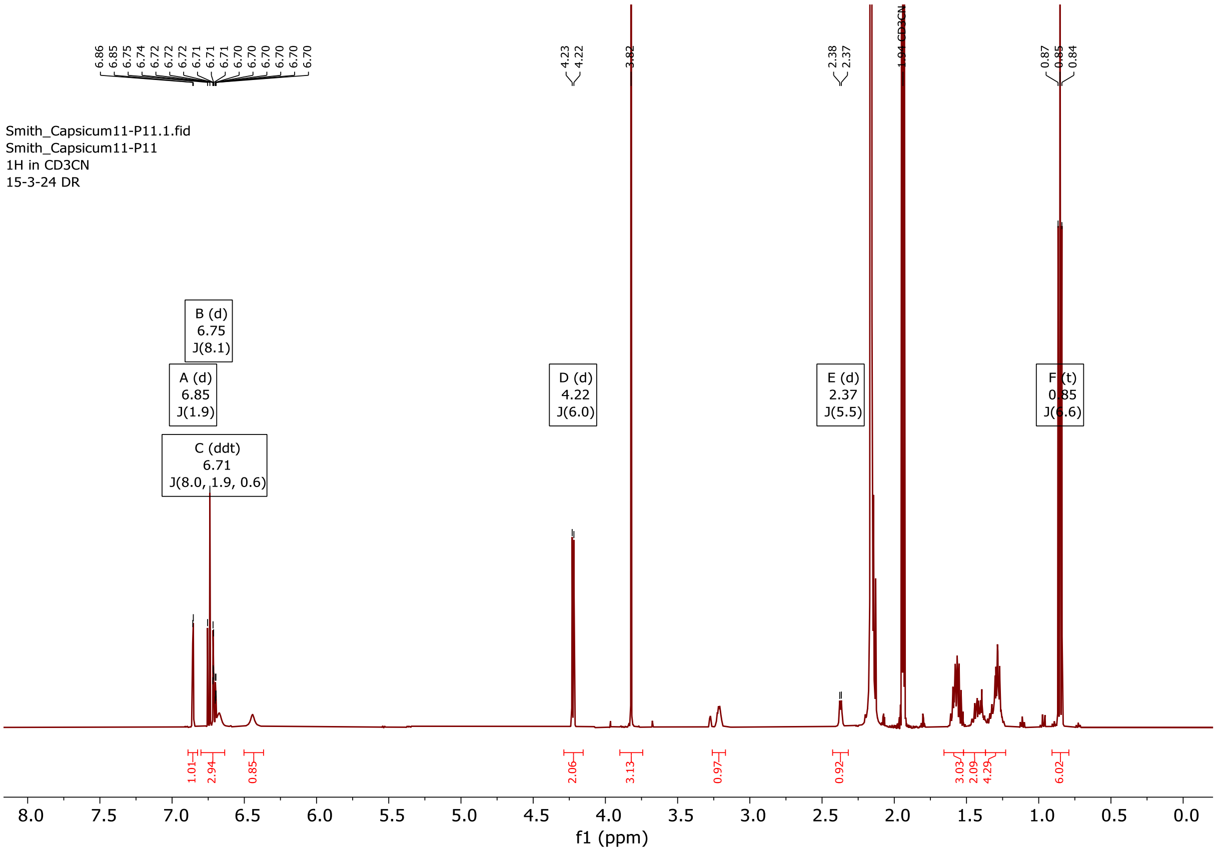

[1]H NMR spectrum of compound **34**.

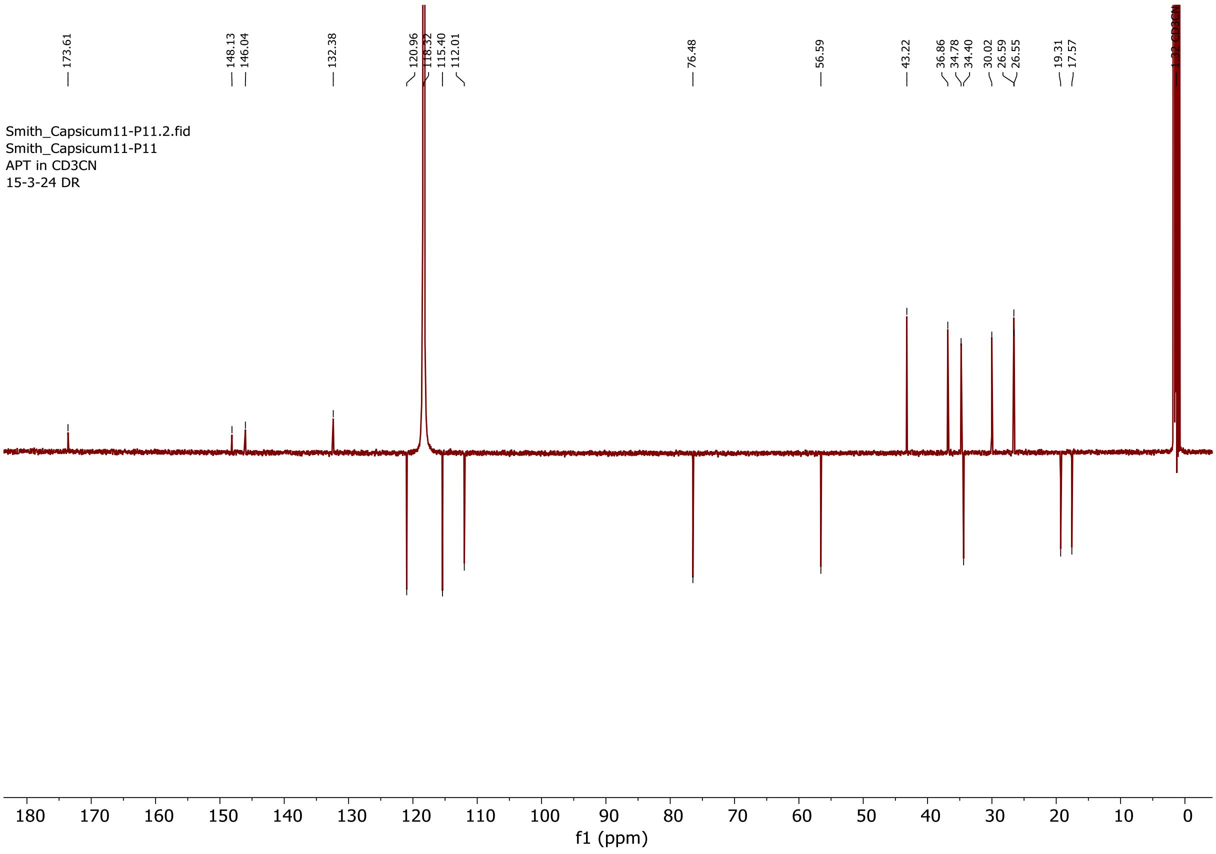

[13]C NMR spectrum of compound **34**.

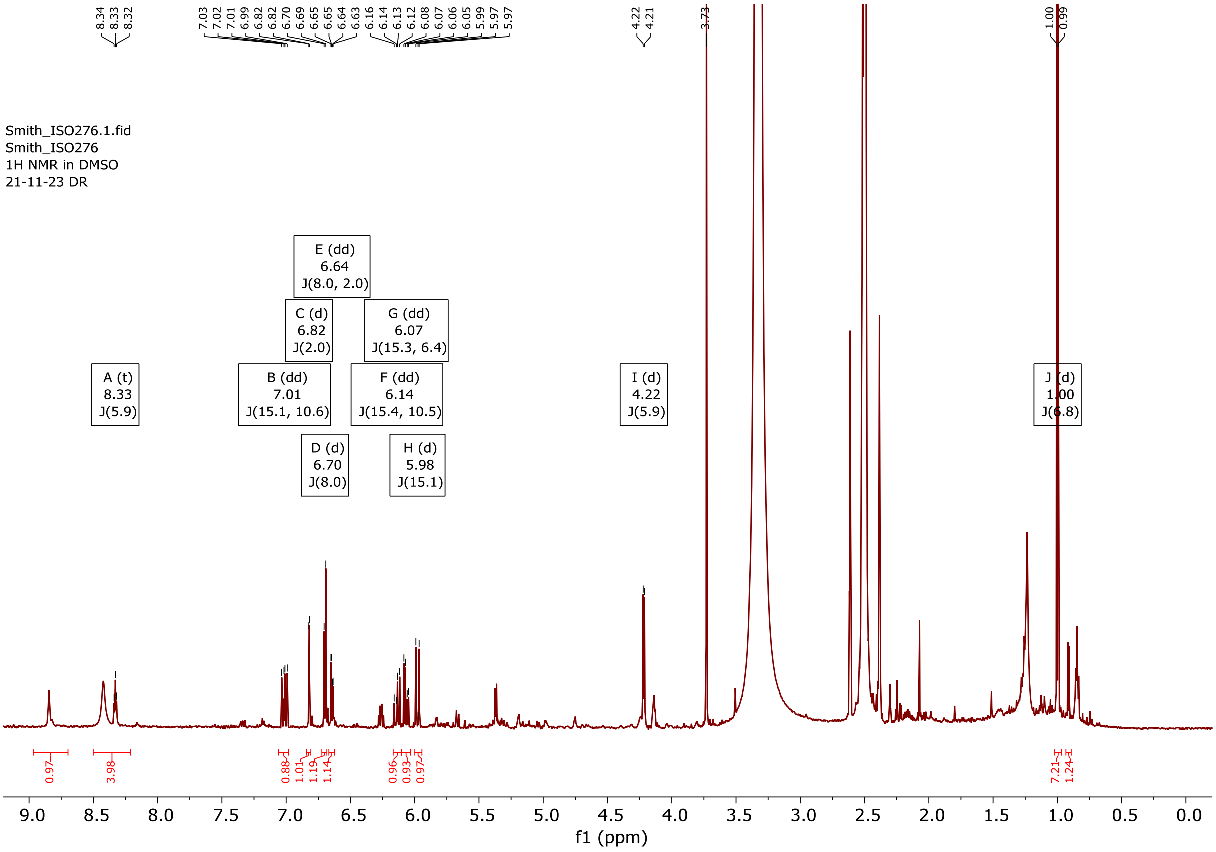


$^{1}$H NMR spectrum of compound **35**.

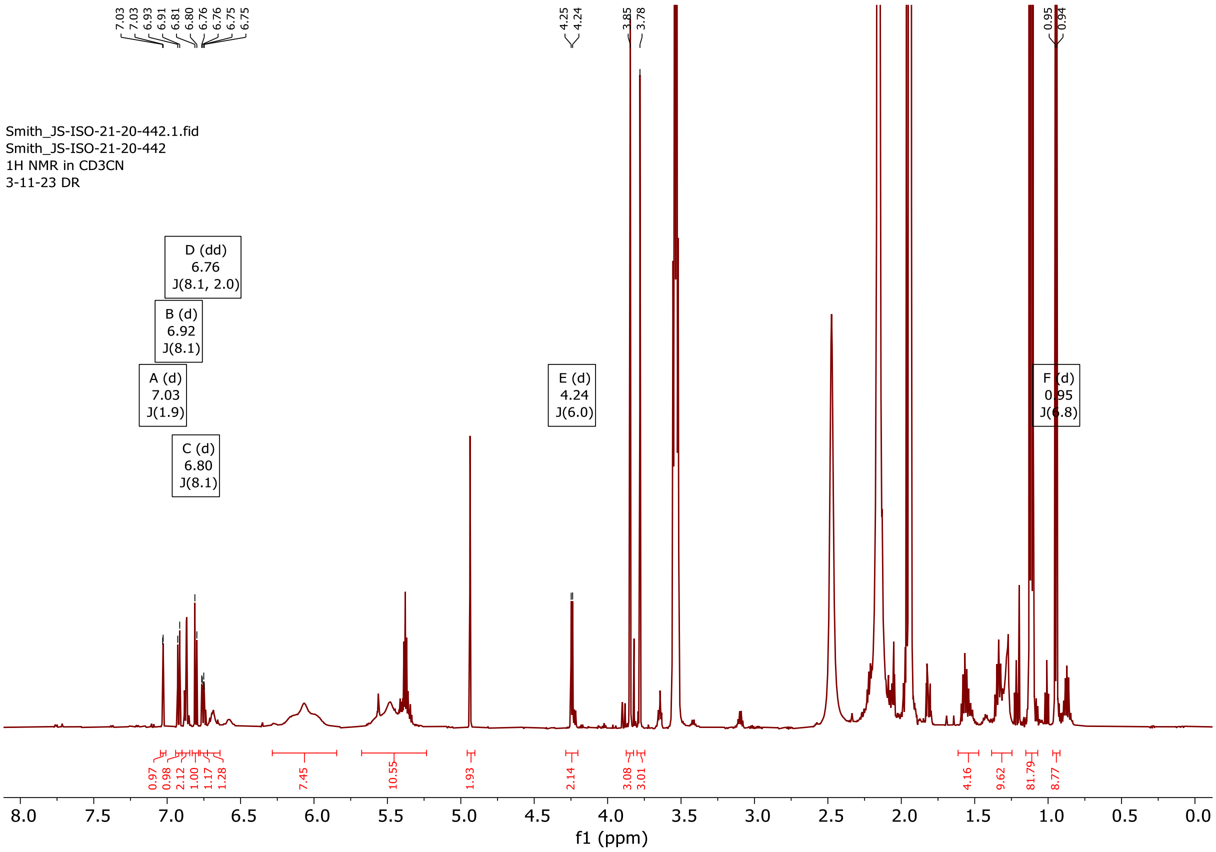


$^{1}$H NMR spectrum of compound **36**.

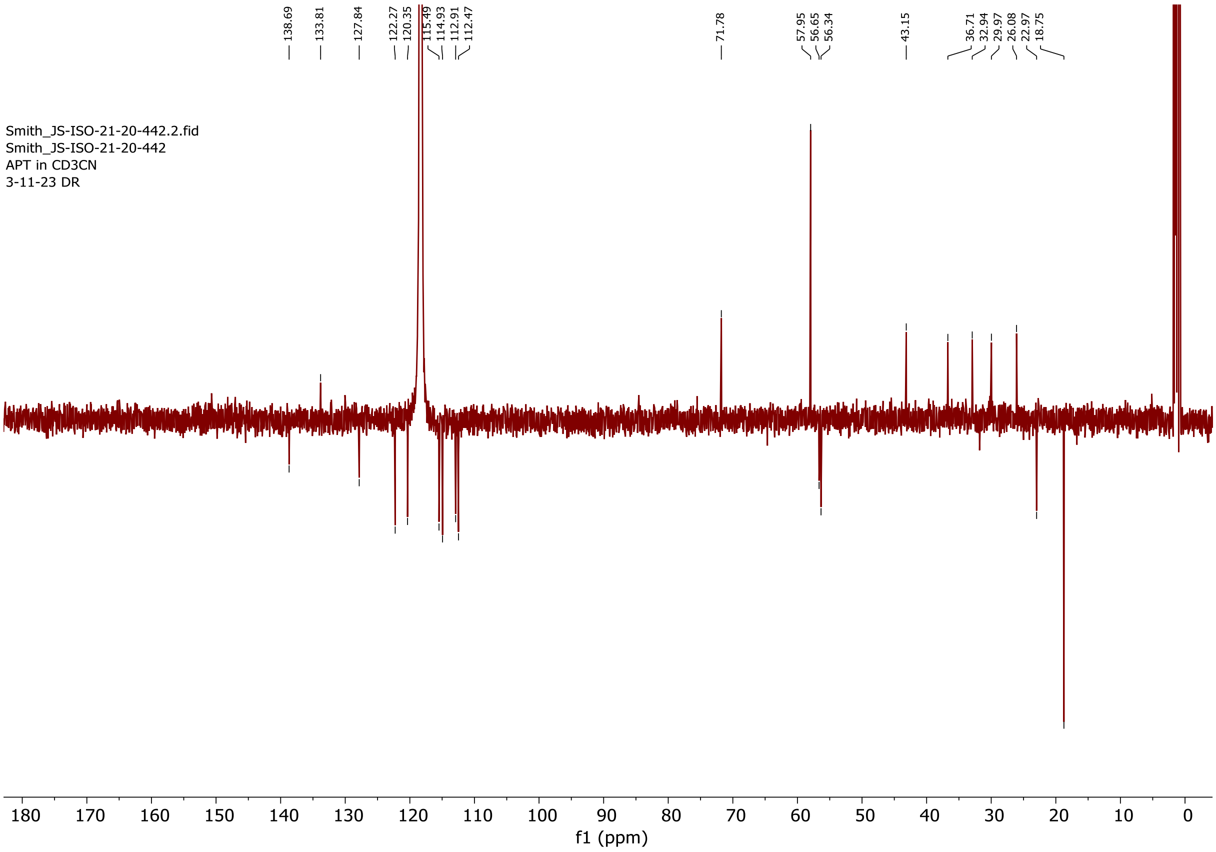


$^{13}$C NMR spectrum of compound **36**.

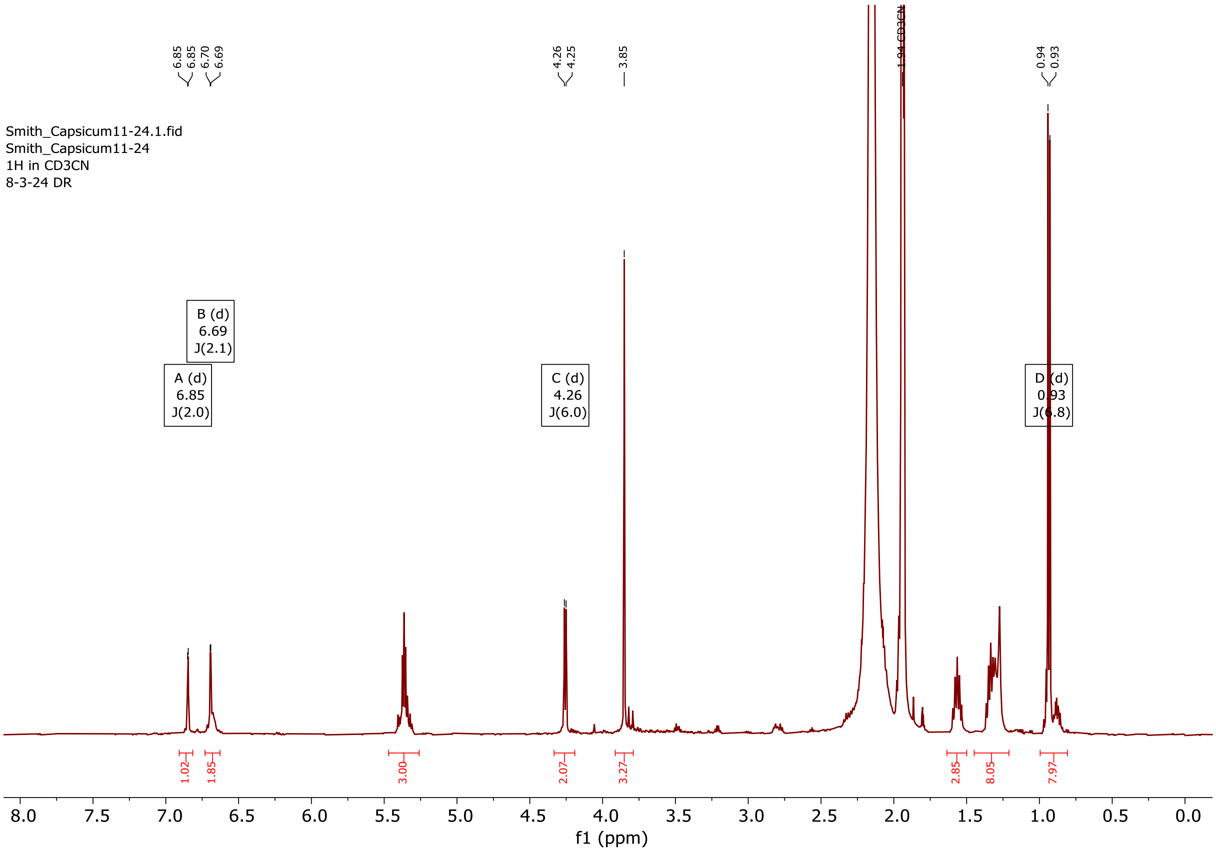


$^{1}$H NMR spectrum of compound **38**.
